# EvAM-Tools: tools for evolutionary accumulation and cancer progression models

**DOI:** 10.1101/2022.07.05.498481

**Authors:** Ramon Diaz-Uriarte, Pablo Herrera-Nieto

## Abstract

EvAM-Tools is an R package and web application that provides a unified interface to state-of-the-art cancer progression models (CPMs) and, more generally, evolutionary models of event accumulation. The output includes, in addition to the fitted models, the transition (and transition rate) matrices between genotypes and the probabilities of evolutionary paths. Generation of random cancer progression models is also available. Using the GUI in the web application, users can easily construct models (modifying Directed Acyclic Graphs —DAGs— of restrictions, matrices of mutual hazards, or specifying genotype composition), generate data from them (with user-specified observational/genotyping error), and analyze the data.

**Availability and Implementation:** Implemented in R and C; open source code available under the GNU Affero General Public License v3.0 at https://github.com/rdiaz02/EvAM-Tools. Docker images freely available from https://hub.docker.com/u/rdiaz02. Web app freely accessible at https://iib.uam.es/evamtools.

## 1 Introduction

Cancer progression models (CPMs) try to identify dependencies between discrete genetic events (e.g., mutations) from cross-sectional data (Beerenwinkel *et al*., 2015, 2016; Diaz-Uriarte, 2018). These dependencies can be deterministic, as in Oncogenetic Trees (OT) (Desper *et al*., 1999; Szabo and Boucher, 2008), Conjunctive Bayesian Networks (CBN) (Gerstung *et al*., 2009; Montazeri *et al*., 2016), OncoBN (Nicol *et al*., 2021), and Hidden Extended Suppes-Bayes Causal Networks (H-ESBCNs) (Angaroni *et al*., 2021), or stochastic as in Mutual Hazard Networks (MHN) (Schill *et al*., 2020). These models also implicitly encode the possible mutational trajectories with predictions about their probability, and have been used for predicting cancer evolution, both long-term (Diaz-Uriarte and Vasallo, 2019; Hosseini *et al*., 2019) and short-term (Diaz-Colunga and Diaz-Uriarte, 2021). Although developed in the field of computational oncology, these models are not limited to cancer: they can be applied to other questions involving the irreversible accumulation of discrete items (Gotovos *et al*., 2021), and have been used to examine tool use in animal taxa (Johnston and Røyrvik, 2020). Their wide utility derives from the assumptions these methods make about the cross-sectional data, which can apply to a broad range of domains. The key assumptions are that (Beerenwinkel *et al*., 2015, 2016; Diaz-Uriarte and Vasallo, 2019; Gerstung *et al*., 2011): a) events are gained one by one; b) once gained, events are not lost; c) we can consider the different individuals in the cross-sectional data as replicate evolutionary experiments where all individuals are under the same constraints (e.g., genetic constraints if we are dealing with mutations) —see further details in the additional documentation.

However, everyday use of these methods, and comparison of output from different ones, is hindered because they are implemented in different languages and with dependencies that make them difficult to install under some operating systems, and they use heterogeneous formats for input and output. In addition, output is often textual (not graphical) and most implementations do not provide the output users might be interested in, such as probabilities of evolutionary paths and genotypes. Finally, it is difficult to explore how changes in genotype composition affect the fitted models and to analyze with one model data generated under another. Here we present EvAM-Tools, an R package and web app for evolutionary and event accumulation models, that facilitates the use and comparison of CPMs, and their extension to other problems.

## 2 Functionality

Figure 1 shows an overview of the functionality of EvAM-Tools.

**Figure 1:**
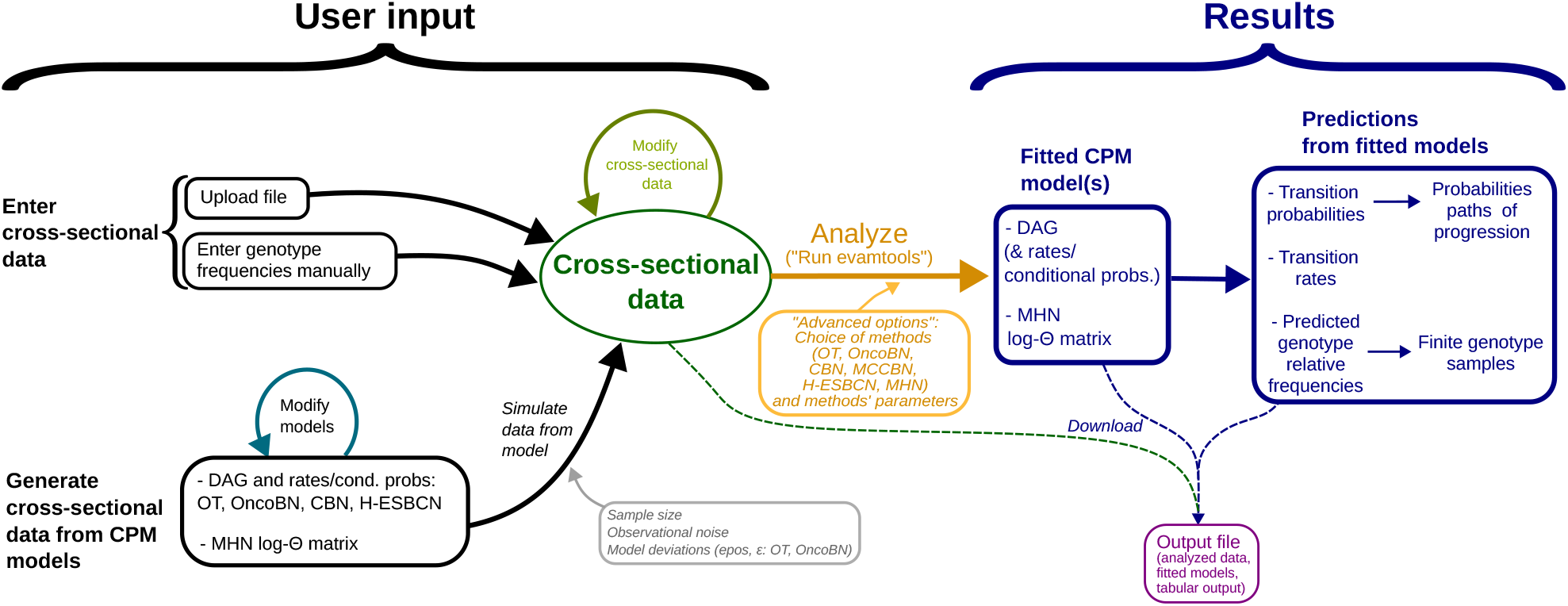
Overview of functionality and workflows with EvAM-Tools (including methods, input, and output); emphasis on functionality provided by the Shiny web app.

### 2.1 Methods available

EvAM-Tools is available as an R package, evamtools, that can also launch an interactive GUI as a Shiny (Chang *et al*., 2021) web app. A distinctive feature of EvAM-Tools is the unified input and usage of five CPMs, the available state-of-the-art procedures with existing implementations: OT (Desper *et al*., 1999; Szabo and Boucher, 2008), CBN (both H-CBN — Gerstung *et al*., 2009 — and MC-CBN — Montazeri *et al*., 2016), OncoBN (Nicol *et al*., 2021), H-ESBCN *(Angaroni et al*., 2021), MHN (Schill *et al*., 2020). OT represents restrictions as a tree (each event can have only one parent); they are untimed models, and weights along the edges are probabilities of transition by the time of observation. CBN represents restrictions with directed acyclic graphs (DAGs) so an event can have multiple parents with an AND relationship (all parent events must have occurred for the child event to appear); in contrast to OT, CBN is a timed model: the parameters of nodes are rates of the exponentially distributed times to event occurrence. OncoBN is untimed like OT but restrictions are represented as DAGs with either AND or OR dependencies (OR dependency: occurrence of one of the parent events is enough for the child to occur). H-ESBCN is timed like CBN, but allows for AND, OR, and XOR (exactly one of the parents must occur) in the DAG of dependencies. MHN uses a non-deterministic timed model where the rate of acquisition of an event is the product of the baseline hazard of the event and the multiplicative effects (promoting or inhibiting) that other events can have via pairwise interactions; there is no DAG as the model is fully specified by the Θ matrix of hazards and multiplicative effects. (See the additional documentation for a more complete description of the methods, including similarities and differences, assumptions, input, and output.)

### 2.2 Input

EvAM-Tools allows for different types of input (Figure 1). The analyses *sensu stricto* start from cross-sectional data, a matrix of individuals (or patients) by events (or genes, pathways, SNPs, CGH regions, etc), with a 1 denoting the event (e.g., a mutation in a gene) is present and a 0 denoting the event is not present. This cross-sectional data themselves can be:

- Uploaded from a file.
- Entered manually by specifying genotype frequencies.
- Generated from CPM models: users can create/modify the DAGs of restrictions (OT, CBN, OncoBN, H-ESBCN) or the log-Θ matrix (MHN) that define the models, and then generate synthetic data from the models (after specifying the size of the sample and the amount of noise, from both model error and observational noise).

File uploading and generation of synthetic data from CPM models are available in both the R package and the Shiny web app. The R package also provides a function for the generation of random CPMs/evam models (including OT, OncoBN in conjunctive and disjunctive forms, CBN, H-ESBCN with AND, XOR, OR dependencies, and MHN). The Shiny web app provides a **GUI** that allows to interactively modify: (a) the CPM models, (including the DAG of restrictions, the type of model —OT, OncoBN, CBN, H-ESBCN— and dependencies —AND, OR, XOR—, and their rates or conditional probabilities), and the log-Θ matrix for MHN models; (b) genotype frequencies, both from uploaded data and from data generated from CPM models. To facilitate exploration of the methods, the Shiny web app includes preloaded examples of cross-sectional data sets, DAGs, and MHN models.

### 2.3 Output

Output includes:

- DAGs of restrictions annotated with parameter values (OT, CBN, OncoBN, H-ESBCN) and log-Θ matrix (MHN).
- Plots and tables of the transition probabilities between genotypes and transition rate matrices from the continuous-time Markov Chain (for models where this exists: CBN, H-ESBCN, MHN). As visualizing the acquisition of mutations in a complex network can be challenging, we use the representation of the hypergraph transition graph from HyperTraPS (Greenbury *et al*., 2020); to enhance interpretability, the user can modify the number of paths to show and the type of annotation.
- Plots and tables of the predicted probabilities of genotypes and finite samples from the predicted probabilities with user-specified observational (e.g., genotyping) error levels.
- List of of evolutionary paths and their probabilities.

When using EvAM-Tools from R, output is available as standard R objects. The web app allows users to download data sets and analysis output.

### 2.4 Workflows

EvAM-Tools encompasses different workflows, including (Figure 1):

1. Inference of CPMs from user data uploaded from a file.
2. Exploration of the inferences that different CPM methods yield from manually constructed synthetic data.
3. Construction of CPM models (DAGs with their rates/probabilities and MHN models) and simulation of synthetic data from them.

3.1 Examination of the consequences of different CPM models and their parameters on the simulated data.
3.2 Analysis of data simulated under one model with methods that have different models (e.g., data simulated from CBN analyzed with OT).
3.3 Analysis of data simulated under one model with manual modification of specific genotype frequencies prior to analyses.

Examples that illustrate these workflows, with both real and simulated data, are available in the additional documentation.

## 3 Use cases

EvAM-Tools can be used, among others, as:

- A teaching tool to understand what CPMs imply about observed data and how different inputs affect the fitted models.
- A research tool to analyze cross-sectional data with state-of-the-art CPMs.
- A research tool in methodological work to compare the performance of different methods under data generated by other models or under specific evolutionary scenarios, and to evaluate new methods.

## 4 Tests, documentation, and availability

The package includes more than 1400 tests that run at every build/check cycle. These tests provide a code coverage of more than 90%. The Shiny web app includes usage documentation in the landing page, as well as in the application itself using: (a) short documentation strings next to each option; (b) tooltips; (c) longer help, via clickable “Help” buttons. The evamtools R package includes standard R function documentation. The repository README includes installation instructions. The additional documentation includes how to use OncoSimulR (Diaz-Uriarte, 2017) to compute transition probabilities between genotypes to provide additional testing and make fitness models explicit; details about CPMs —models, algorithms, and assumptions—, predicting genotype frequencies and generating random evam models; a FAQ; commented examples, with both real and simulated data, that illustrate the use and utility of EvAM-Tools.

The web app can be run on our servers or locally. In the latter case after installing the package from source or using one of the Docker images we provide, which include all dependencies and will run both the package and the web app.

## 5 Conclusion

EvAM-Tools provides a unified interface to state-of-the-art CPMs, a user-friendly GUI, output (graphical and tabular) that extends the usual output derived from these methods, and functionality that facilitates methodological work. EvAM-Tools should result in better understanding and increased use of these methods and encourage further methodological work and their extension to other scenarios, making it a tool of interest for computational oncologists and more broadly to evolutionary biologists.

## Supporting information

EvAM-Tools examples

EvAM-Tools methods' details and FAQ

## Acknowledgements

C. Lazaro-Perea and two anonymous reviewers for comments that helped improve the ms.

## Funding

Work supported by grant PID2019-111256RB-I00 funded by MCIN/AEI/10.13039/501100011033 to RDU. PHN supported by Comunidad de Madrid’s PEJ-2019-AI/BMD-13961 to RDU.

## Notes

https://github.com/rdiaz02/EvAM-tools

### Competing Interest Statement

The authors have declared no competing interest.

### Summary of Updates

- Figure showing workflows. - Explained type of input. - Section on workflows. - Brief description of methods in main ms. - Brief description of assumptions in main ms. - Supplementary data, examples: added many examples, and related to workflows figure. - Supplementary data, methods and FAQ: extended description of methods, added entries to FAQ.

