## Supplementary material for "EvAM-Tools: tools for evolutionary accumulation and cancer progression models": EvAM-Tools examples

Ramon Diaz-Uriarte*

2022-10-03
Version 44b7274

**5 Simulating random CPMs/evams** **41**

**6 Appendix: getting the BRCA and Ov data sets from the R console** **43**

**7 License and copyright** **44**

**8 References** **45**

### 1 Introduction

Here we present examples, with both real and simulated data, that illustrate the use and utility of EvAM-Tools. Most of the previous examples can be run in the web app, and that is what we use here. In the final section we discuss simulating random models, using the R package.

The objective of this document is not to provide complete analyses of any of the data sets, or address any of the questions mentioned above in full (that would require full papers). The objective is to illustrate the use of EvAM-Tools, especially its web app, and also to include some examples of output that, although not necessarily very common, are not unusual and can seem surprising at first (e.g., the variability among fitted H-ESBCN models, in *"Is it just sample size?", section 2.2*, or output with DAGs that are not transitively reduced in *"A model with AND", section 4.3*).

#### 1.1 Web app: overview of workflow and functionality, and relationship to these examples

On the web app landing page, under "About EvAM-tools" (`https://www.iib.uam.es/evamtools/#overview`) we provide an overview of the workflow and use cases. For completeness we repeat that material here, indicating, in bold, how the examples in this document relate to the major functionalities and workflows discussed there.

The figure below provides an overview of the major workflows with the web app:

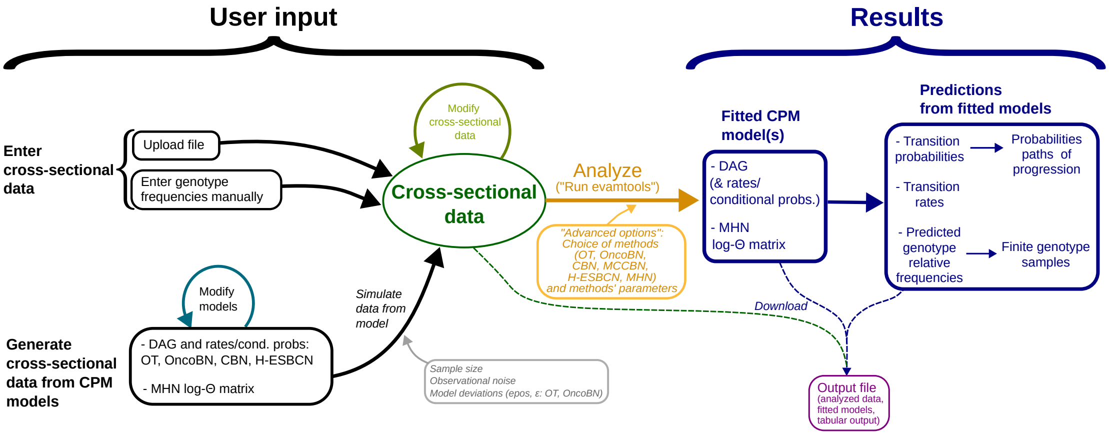

The web app encompasses, thus, different major functionalities and workflows, mainly:

1. Inference of CPMs from user data uploaded from a file. **Examples *"Analyzing the BRCA data set" (section 2.1)* and *"Analyzing the ovarian CGH data" (section 2.3)*.**

2. Exploration of the inferences that different CPM methods yield from manually constructed synthetic data. **Example *"Is it just sample size?" (section 2.2)* uses a modification of uploaded data to explore changes in sample size; example *"Analyzing manually constructed synthetic data" (section 3)* constructs synthetic data to examine the effect of aliasing of events.**

3. Construction of CPM models (DAGs with their rates/probabilities and MHN models) and simulation of synthetic data from them.

   3.1. Examination of the consequences of different CPM models and their parameters on the simulated data. **Examples *"What happens if we increase $\epsilon$ for OncoBN?" (section 4.1.1)*, *"A simple exploration of MHN" (section 4.1.2)*.**

3.2. Analysis of data simulated under one model with methods that have different models (e.g., data simulated from CBN analyzed with OT and OncoBN). **Examples** ***“A model with AND, XOR, OR” (section 4.2)*** **and** ***“A model with AND” (section 4.3)***.

3.3. Analysis of data simulated under one model with manual modification of specific genotype frequencies prior to analyses (e.g., data simulated under CBN but where, prior to analysis, we remove all observations with the WT genotype and the genotype with all loci mutated). **Example** ***“Modifying data generated from a CPM model before analysis” (section 4.4)***.

The figure below highlights the different major functionalities and workflows, as numbered above, over-imposed on the previous figure:

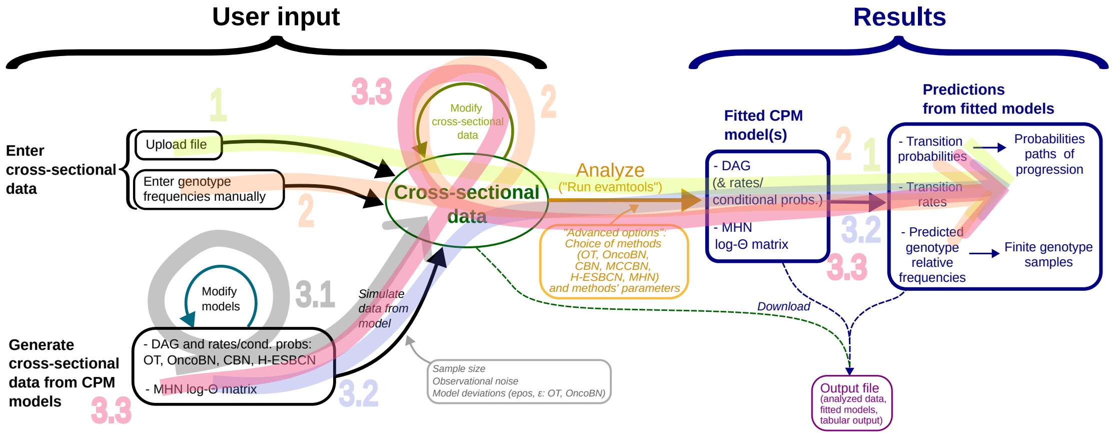

Furthermore, note that in all cases, when data are analyzed, in addition to returning the fitted models, the web app also returns the analysis of the CPMs in terms of their predictions such as predicted genotype frequencies and transition probabilities between genotypes. Most of the examples below illustrate this, showing, for example, the predicted genotype frequencies and transition probabilities.

##### 1.2 Additional documentation

See additional documentation in https://rdiaz02.github.io/EvAM-Tools. In particular, additional details about CPMs, error models, predicting genotype frequencies, generating random evam models, and a FAQ is available from https://rdiaz02.github.io/EvAM-Tools/pdfs/evamtools_methods_details_faq.pdf.

#### 2 Analysis of cross-sectional data

We will analyze two cancer data sets, the ovarian cancer CGH data included in the Oncotree package (Szabo and Pappas, 2022), and the BRCA data set for basal-like subtypes (from Cerami *et al.*, 2012; Gao *et al.*, 2013, originally from Cancer Genome Atlas Research Network, 2012; see Supplementary File S5_Text, https://doi.org/10.1371/journal.pcbi.1007246.s007 of Diaz-Uriarte and Vasallo, 2019 for full details about data origins and preprocessing).

So that you can use these data sets directly, we provide them in the repository (https://github.com/rdiaz02/EvAM-Tools/tree/main/examples_for_upload). The direct links are:

- BRCA_ba_s.csv: https://raw.githubusercontent.com/rdiaz02/EvAM-Tools/main/examples_for_upload/BRCA_ba_s.csv
- ov2.csv: https://raw.githubusercontent.com/rdiaz02/EvAM-Tools/main/examples_for_upload/ov2.csv

In section *"Appendix: getting the BRCA and Ov data sets from the R console" (section 6)* we show how to obtain the data from the R console.

#### 2.1 Analyzing the BRCA data set

We now import the BRCA csv data set, BRCA_ba_s.csv, into the web app, https://iib.uam.es/evamtools. We go to the "User input" tab, and click, on the left, on "Upload file"; we set the "Name for data" as "BRCA_ba"):

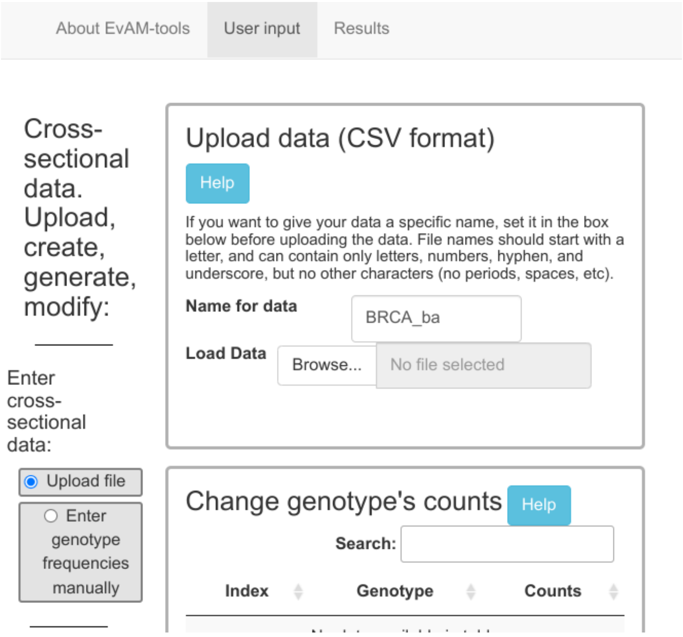

On "Load Data" we click on "Browse" and select the file from our file system; the data is uploaded, and the genotypes' frequencies are shown in the histogram on the right and the table at the bottom (where, if we wanted, we could modify the genotype counts):

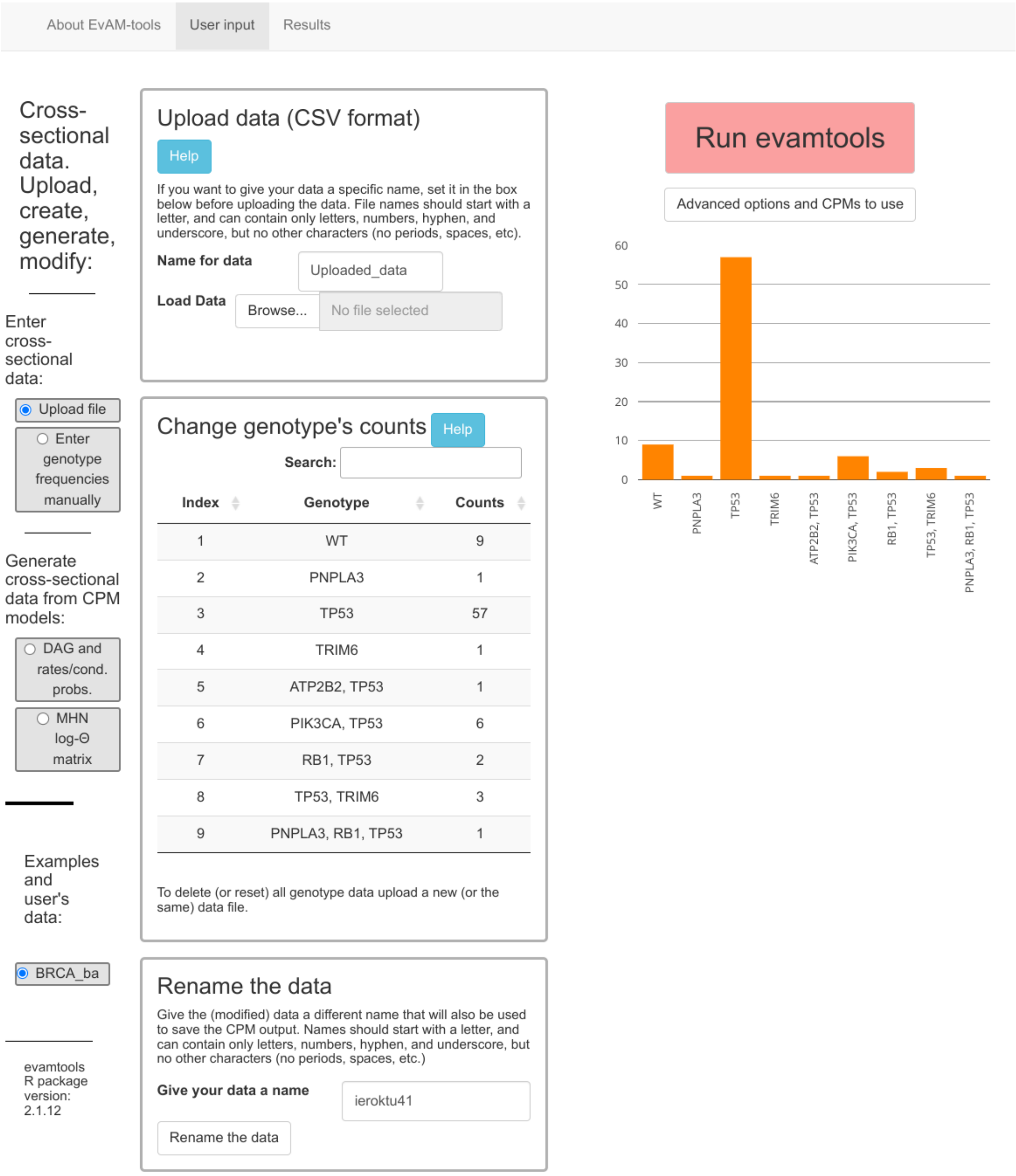

Before running the analysis, we select the unselected H-ESBCN as one of the methods to run (under “Advanced options and CPMs to use”). We also set the number of MCMC iterations to 500000, instead of 200000, for increased stability of results; this increase in iterations will of course result in longer running times.

We click “Run evamtools” and the output is shown in about 30 to 50 seconds[1]. For easier display of the figures in this document, we select first three of the methods to show, by clicking, on the left menu, under “Customize the visualization”:

[1]In this example, changing the setting for the number of MCMC iterations of H-ESBCN from 100000 to 500000 is responsible for increasing the total running time from about 30 to about 50 seconds.

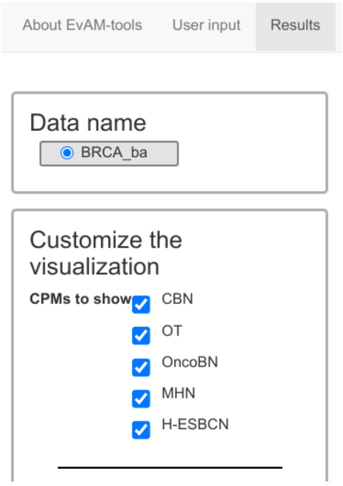

First on the first three (CBN, OT, OncoBN)

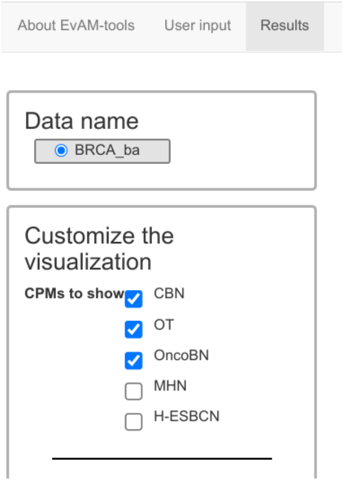

Then on the next two (MHN, H-ESBCN)

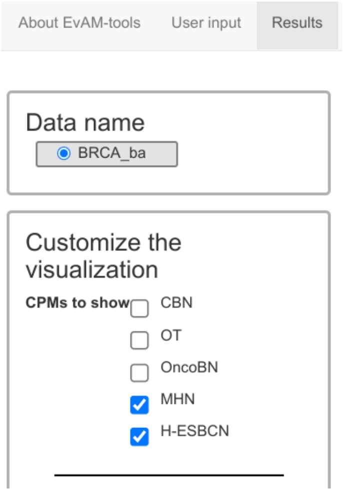

This is the output from the first three methods (CBN, OT, OncoBN):

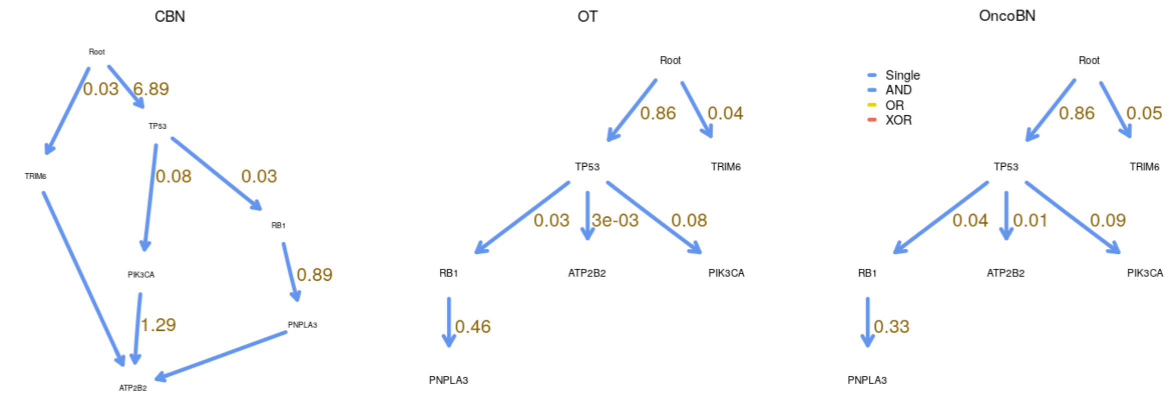

And this from the next two methods (MHN, H-ESBCN):

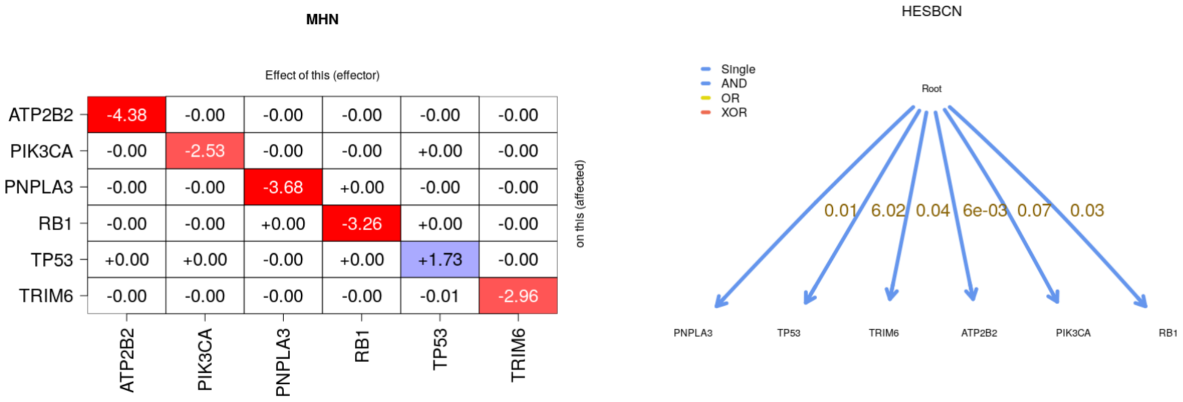

(The above screen-captures only show the DAGs/MHN matrix; we are not showing the figures of the predictions of the fitted models, such as transition probabilities or predicted genotype relative frequencies).

EvAM-Tools makes it immediate to see that:

- The output from OncoBn and OT is essentially identical.
- CBN and H-ESBCN give very different DAGs.
- OT and OncoBN differ from both CBN and H-ESBCN.

That OT and OncoBN give identical results is not surprising since OncoBN is using the disjunctive (OR relationships) model and has not found any disjunctive pattern. We can run OncoBN using the conjunctive model. Go back to “User input” and click on “Advanced options and CPMs to run” and set, for “OncoBN options”, the “Model” to “Conjunctive”:

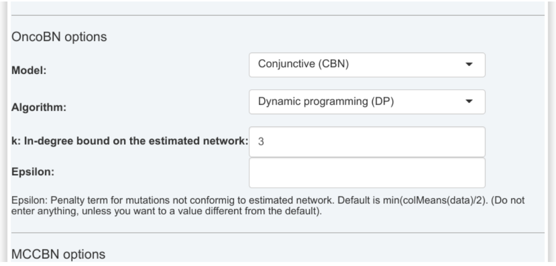

Click on “Run evamtools” to obtain the new fit (since we are only interested in rerunning

OncoBN, we might want to unclick the other methods, so as to make the run as fast as possible). Interestingly, if we run OncoBN with conjunctive pattern, it does not show any conjunctions either (the result is the same as shown above):

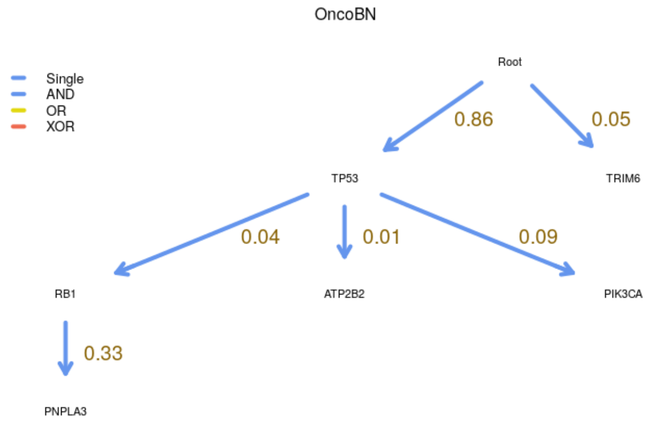

Thus, CBN seems to find support in the data for conjunctive dependencies that neither OncoBN (run using the “Conjunctive” model) nor H-ESBCN find.

EvAM-Tools’s output also displays, in the Results tab, the original data as well as transition probabilities, transition rates, and predicted genotype relative frequencies. We show the three differing DAGs, MHN’s output, and the data, when we select “Transition probabilities” (tabular output truncated in the screen capture):

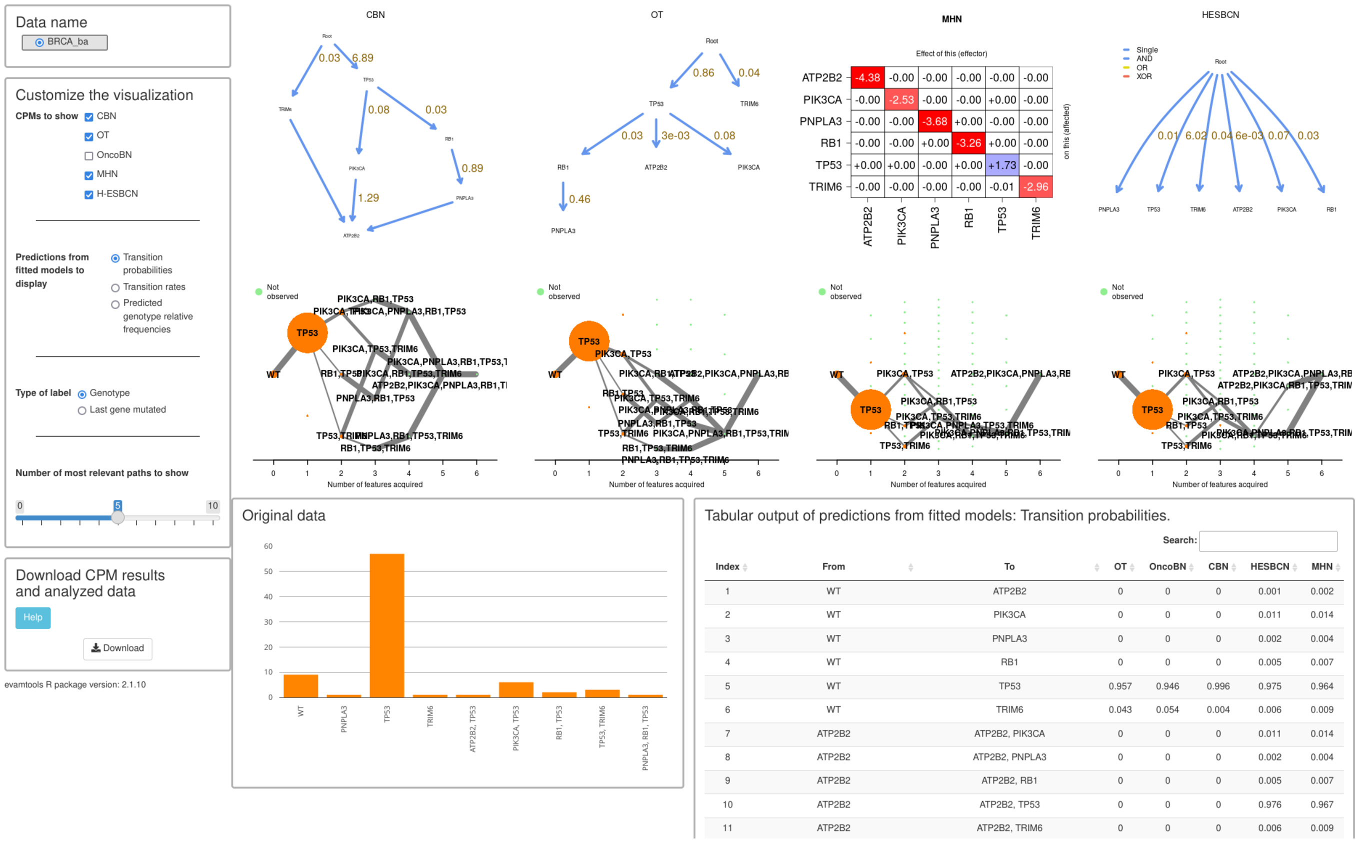

and when we select “Predicted genotype relative frequencies”:

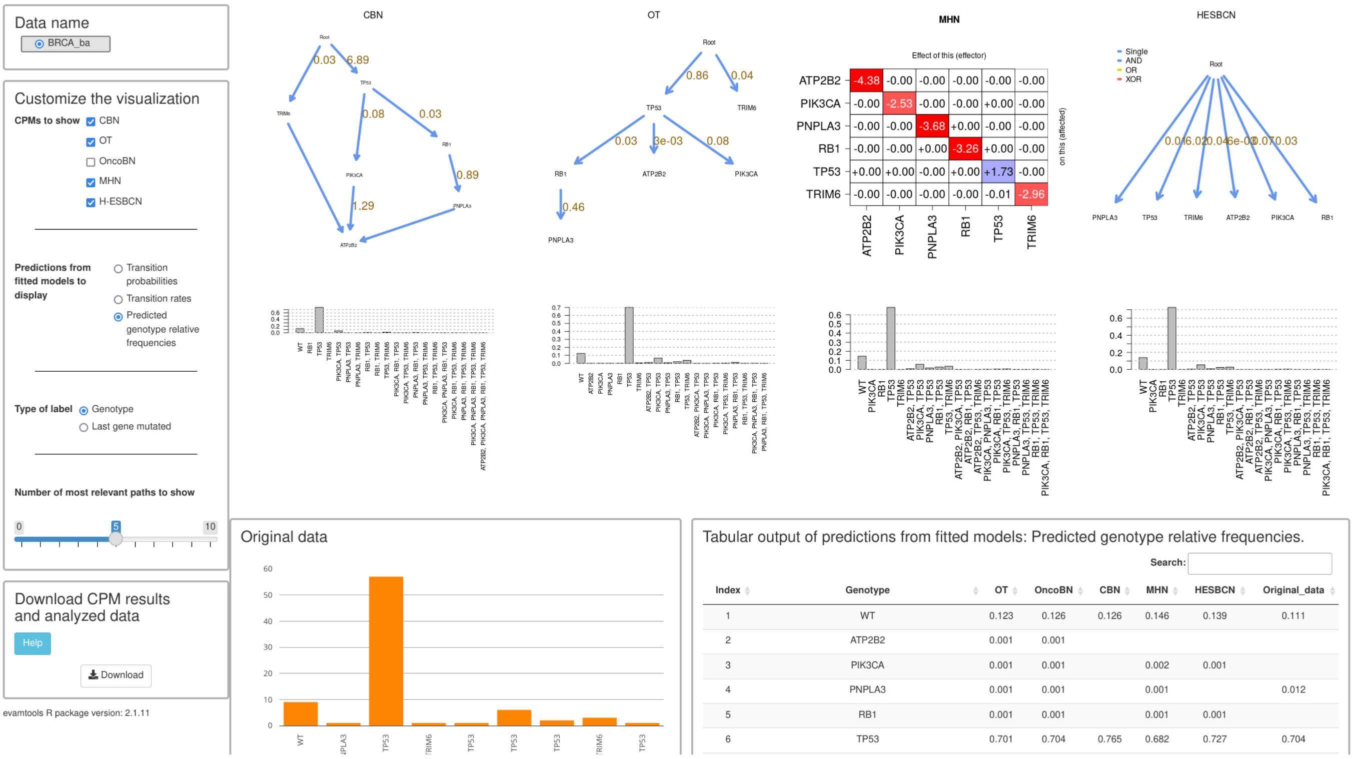

| Index | Genotype | OT | OncoBN | CBN | MHN | HESBCN | Original_data |
|---|---|---|---|---|---|---|---|
| 1 | WT | 0.123 | 0.126 | 0.126 | 0.146 | 0.139 | 0.111 |
| 2 | ATP2B2 | 0.001 | 0.001 | | | | |
| 3 | PIK3CA | 0.001 | 0.001 | | 0.002 | 0.001 | |
| 4 | PNPLA3 | 0.001 | 0.001 | | 0.001 | | 0.012 |
| 5 | RB1 | 0.001 | 0.001 | | 0.001 | 0.001 | |
| 6 | TP53 | 0.701 | 0.704 | 0.765 | 0.682 | 0.727 | 0.704 |

(This, incidentally, shows that we probably would have wanted to use shorter genes names as the histogram labels are too long).

From the above display we can conclude:

- The data contain very few cases where there are joint occurrences of two or more genes: most joint occurrences appear only once.
- The conditional probabilities from OncoBN (or OT) indicate that the really likely event is gaining TP53; the conditional probability of the remaining ones is very small.
- CBN leads to the same conclusion: the only large $\lambda$ is that for TP53.
- MHN’s output points in the same direction: except for TP53, the diagonal entries of the matrix are all negative and large in absolute value, and the off-diagonal entries are all essentially 0. Thus MHN’s model is saying that we can fit the data reasonably well without modeling inhibiting or facilitating relations between genes.

Therefore, we can conclude that the apparently different results are caused by differences in the weighting of evidence: H-ESBCN, given the very small frequencies of most genotypes with more than one mutation, is choosing not to take those as evidence of dependencies, and is instead returning a simpler model.

#### 2.2 Is it just sample size?

We can examine more carefully the conjecture above: would H-ESBCN return a different DAG if sample size were much larger but relative proportions were the same? We can do that easily with EvAM-Tools. We simply go to the “User input” tab and multiply the genotype counts, for example by 10 (which is trivially done by clicking on each cell and adding a 0; we can move between cells using Tab and click “Ctrl + Enter” when we are done).

We rename the data first, and then increase the sample size. This is what it looks like:

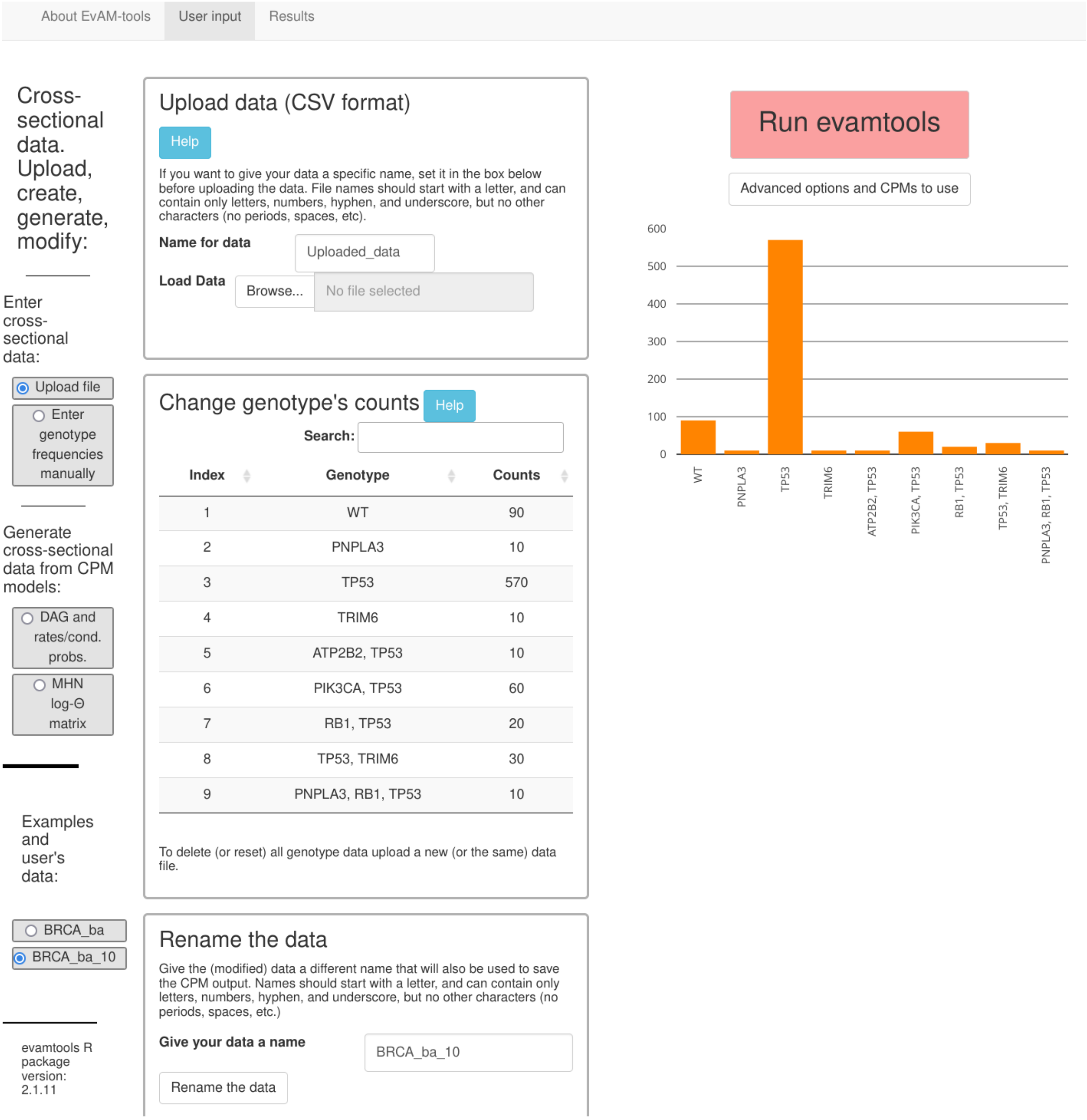

Now, we rerun the analyses:

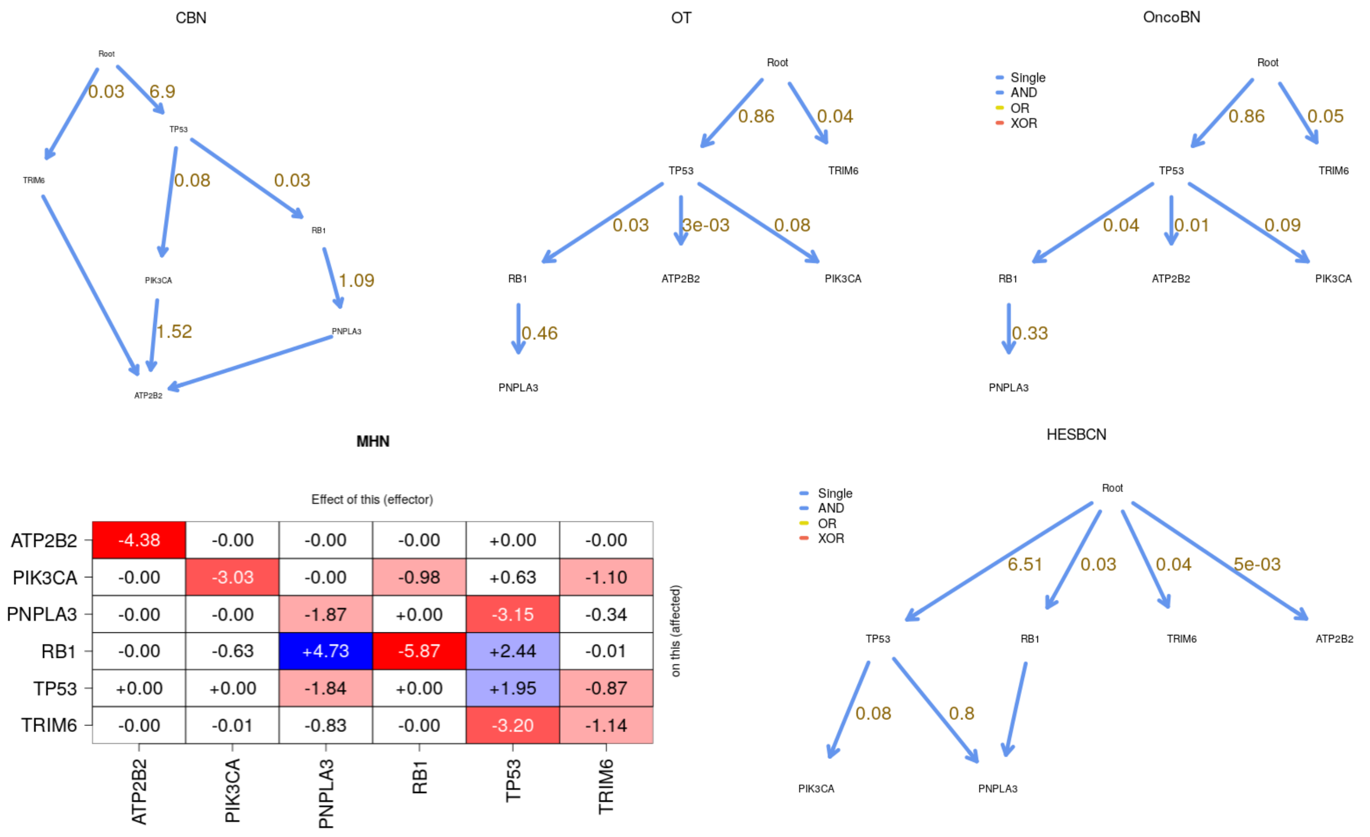

The DAGs and weights (lambdas, probabilities) are the same for OT, OncoBN and CBN. But the models inferred by MHN and H-ESBCN have both changed, and the second now includes dependencies between some of the genes. Some of these dependencies are similar to the ones in the CBN output (PNPLA3 depends on RB1 and TP53; PIK3CA depends on TP53).

Interestingly, increasing the sample size another 10 times results in additional changes in the MHN and H-ESBCN models (OT, OncoBN, and CBN only show minor changes, not shown below):

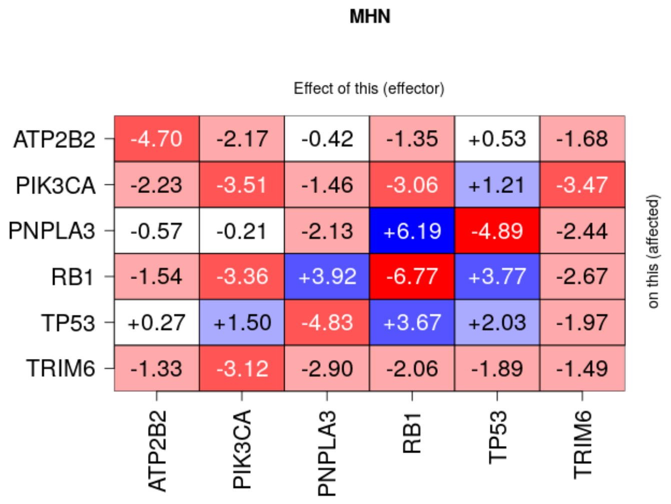

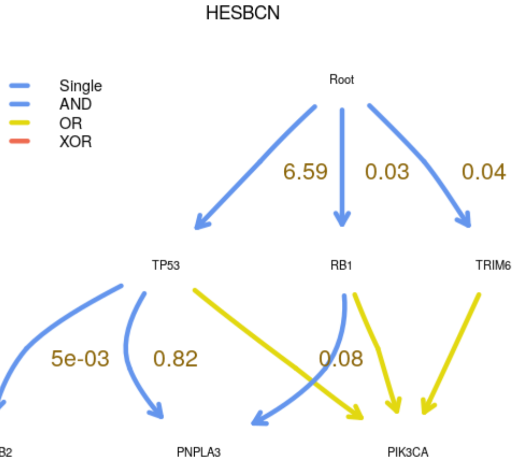

Note that in some runs, the output returned by H-ESBCN is different from the one above; with H-ESBCN different runs can sometimes lead to different results. You can fix the random number seed in “Advanced options” to prevent this, though this is not advisable, since fixing the seed would precisely prevent us from seeing the instability of the fitted models. For example, if you set the number of MCMC iterations (under “Advanced options”) to 100000 and the seed to 19, you will obtain a model with an XOR[2]. In the web app we use, by default, 200000 MCMC iterations, a number larger than the default in `https://github.com/danro9685/HESBCN`, precisely to minimize this instability. In the examples in this document we use an even larger number of 500000 MCMC iterations to obtain more robust results and because none of the examples shown take longer than about 1 minute to run. Variability of results from different runs can also be observed with CBN sometimes (though, in our experience, it is less common than with H-ESBCN with default parameters).

These results lead us to conclude tentatively that, compared to OT, CBN, and OncoBN, the penalties used in H-ESBCN and MHN seem to have a larger effect on models fitted to modest sample sizes.

[2]Note, though, that the output with the XOR also contains an edge with an extreme weight, which makes this model suspect; more detailed exploration would use a range of seeds, and possibly change also the number of MCMC iterations to run.

#### 2.3 Analyzing the ovarian CGH data

Lest readers think that the above (coincidence between OT and OncoBN, and H-ESBCN returning star models with moderate sample sizes) are general patterns, we now show, for a different example, the analyses of the ovarian CGH data. We upload the data `ov2.csv` and, as before, include H-ESBCN in the methods and set the number of MCMC iterations of H-ESBCN to 500000.

This is the output from the web app (run takes a little over one minute):

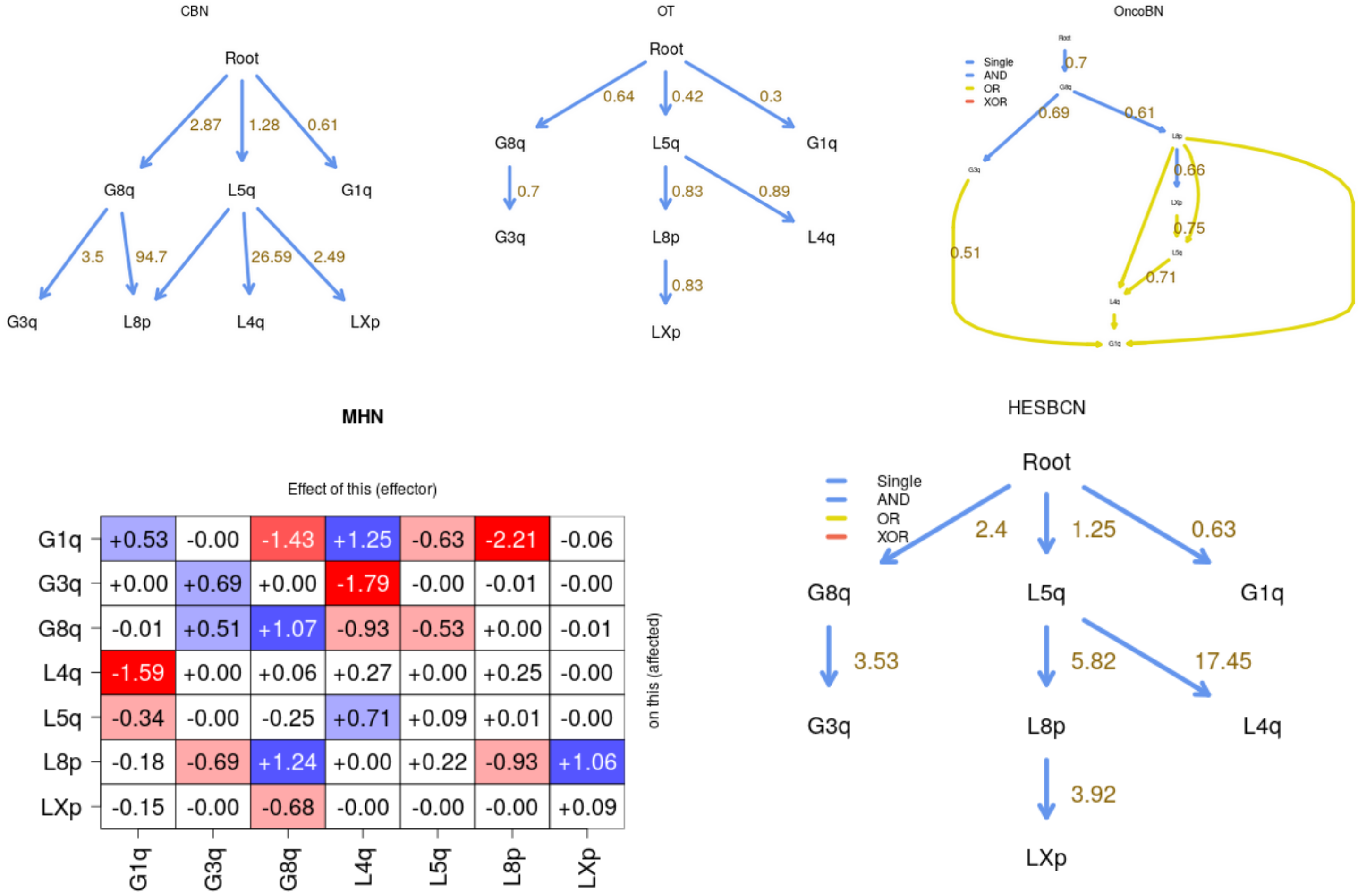

And we re-run the OncoBN model using the “Conjunctive” options (instead of the default disjunctive —we did this before in section *“Analyzing the BRCA data set”, section 2.1*, and it involves going back to “User input”, clicking on “Advanced options and CPMs to run” and setting, for “OncoBN options”, the “Model” to “Conjunctive”). This is the output:

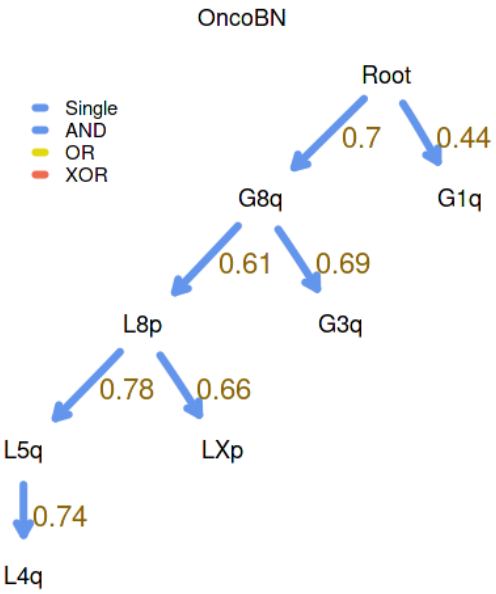

Interestingly, in the above run, H-ESBCN gives an identical structure to that of OT (parameters are, of course, different: OT’s weights are conditional probabilities and H-ESBCN $\lambda$s are rates). Different runs of H-ESBCN can give different results, such as the following ones, where we also ask the web app to display the predicted genotype frequencies:

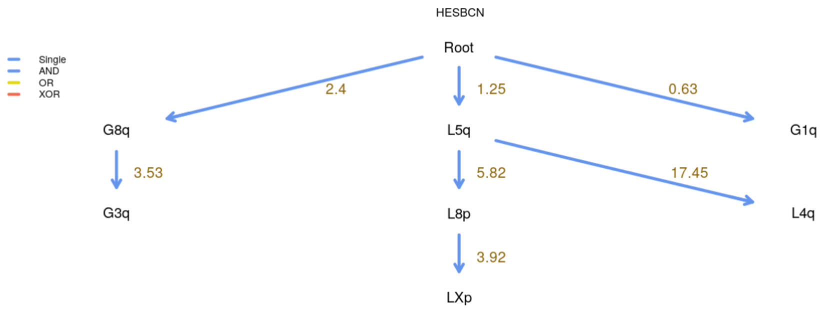

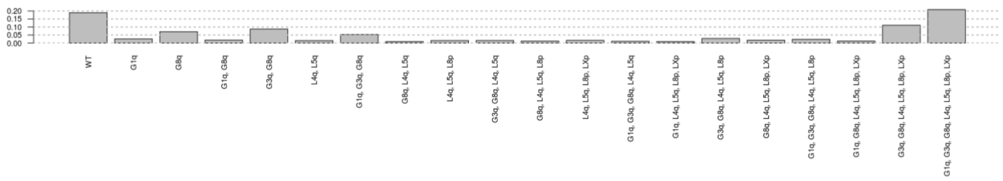

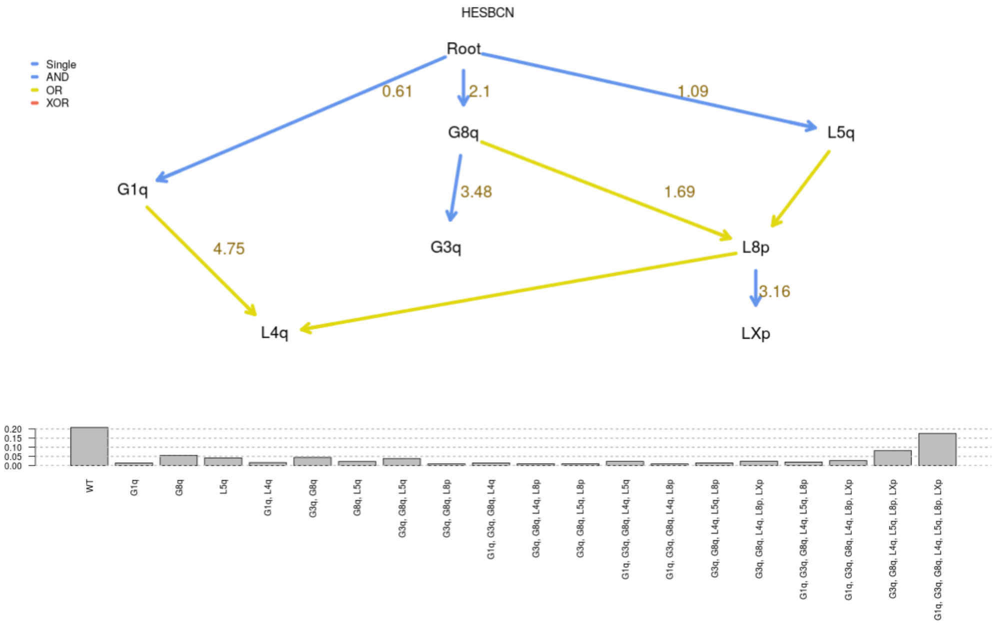

We can increase the number of MCMC iterations. When using 1000000 most runs tend to give this model:

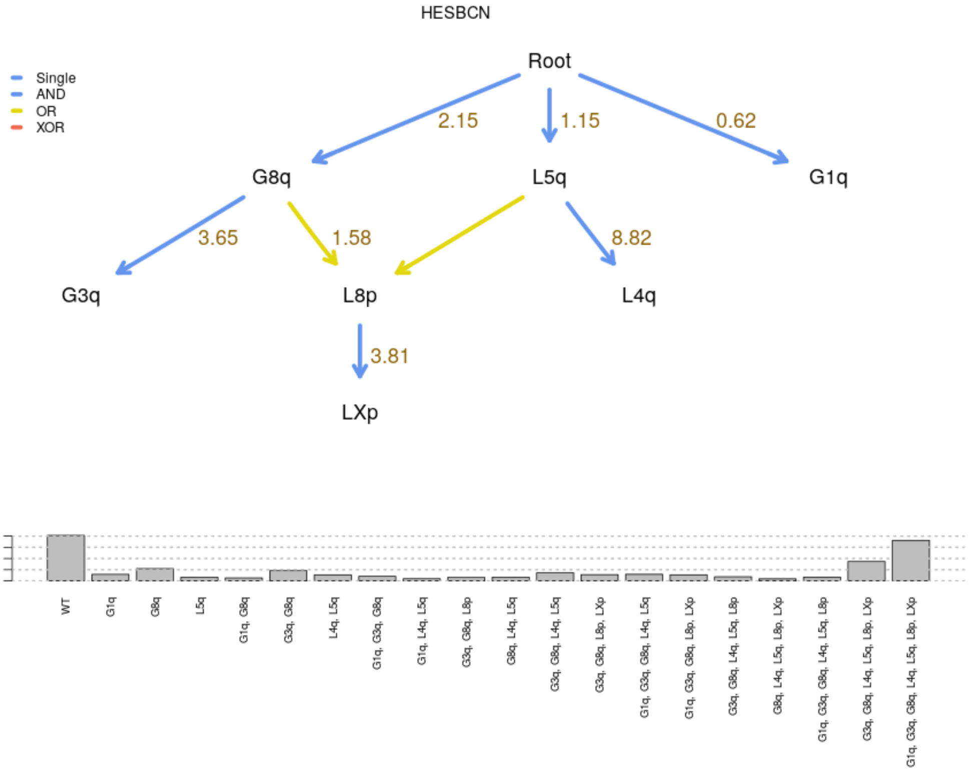

The variability between fitted H-ESBCN models might deserve a more careful exploration and, in "for real" analyses, additional runs with 1000000 iterations or even more.

Now there are large similarities are between OT and CBN (though, of course, there can be no conjunctions in OT). H-ESBCN finds identical restrictions for G3q (3q+) and G8q (8q+) as

OT and CBN; it also finds similar patters to OT for L5q and LXp; note that CBN shows L5q → L8p and L5q → LXp and that H-ESBCN shows an OR for the dependency of L8p on G8q and L5q, whereas CBN, which can only model ANDs, places an AND. OncoBN using the default disjunctive relationship (OR, but not XOR) seems to suggest quite a different model (note that H-ESBCN can also fit OR relationships). Interestingly, the conjunctive model for OncoBN is similar, but not identical, to the ones from OT and CBN.

#### 2.4 Using the web app for small computational experiments

We have shown the output of repeated runs of H-ESBCN, changing the number of MCMC iterations and possibly setting different random number seeds. GUIs and web apps are not the most appropriate tools for a systematic exploration; instead, properly documented code as an R script would be the preferred procedure. For small experiments, however, the web app is fine. A simple way to keep track of what is done is as follows:

1. Upload the data, giving it a meaningful name (under "Name for data"); for example, `ov`.

2. Go to the "Rename the data" box, and add the settings you will use to the name of the data; for example, for 500000 MCMC iterations with seed 14 for H-ESBCN enter `ov_500K_s14` in "Give your data a name", and **click on "Rename the data"**.

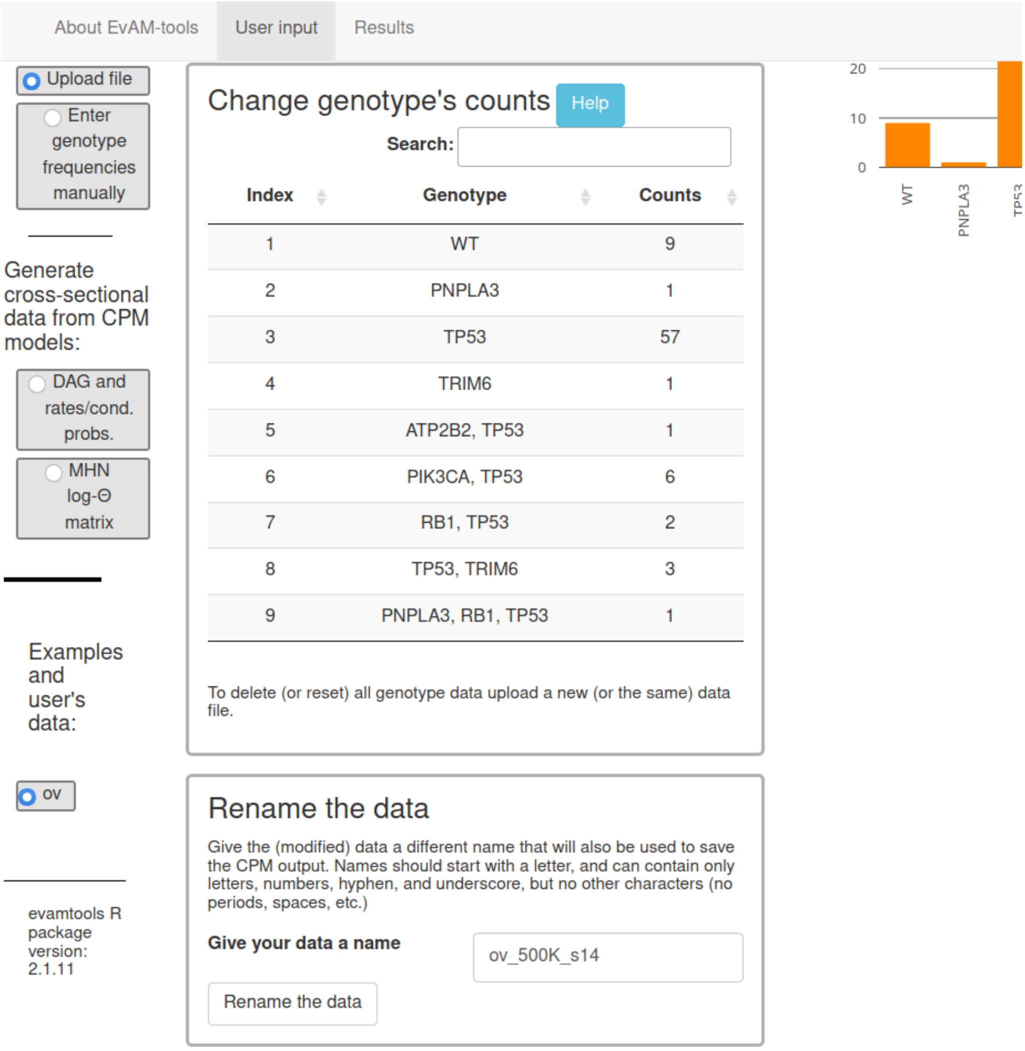

3. Under "Advanced options and CPMs to use" set the seed to 14 and the number of MCMC iterations to 500000, possible also setting H-ESBCN as the only method to run.

4. Click on "Run evamtools".

5. Repeat steps 2 to 4 as needed.

The "Results" tab will contain the output of the different runs, properly labeled so we can examine, at will, outputs from runs with different settings.

### 3 Analyzing manually constructed synthetic data

We will create some synthetic data to show the consequences of analyzing data where two events are indistinguishable, because they are completely aliased, i.e., indistinguishable, because they have identical patterns —identical columns in the data matrix—. Of course, manually constructed synthetic data can be used to explore or examine many other patterns, unrelated to aliased events.

From the “User input” tab we select the “Enter genotype frequencies manually”:

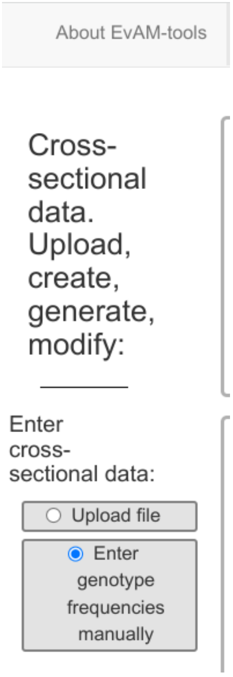

Now, we enter some WT, for example, 20 WT observations. We select no mutations, and type a “20” in “Counts”:

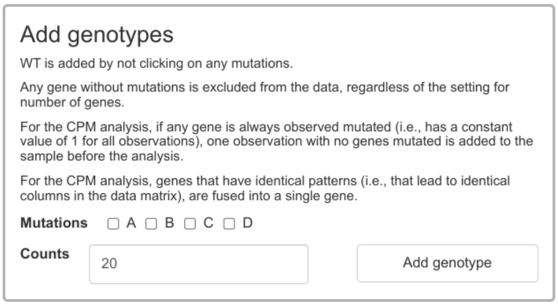

And when we click on “Add genotype” we see the histogram with 20 WT and the genotype table with the 20 WT:

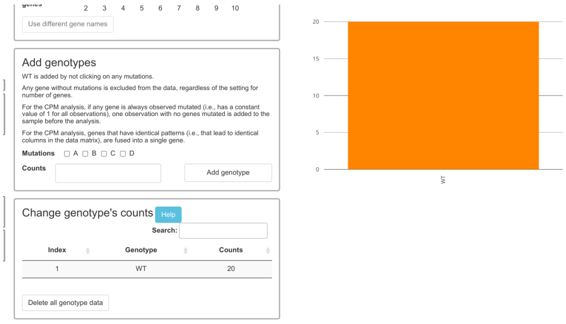

We now add 15 observations with only A mutated (i.e., 15 individuals of genotype A), and 12 with both B and C (i.e., 12 individuals with genotype "B, C"); we first click on "A" on "Mutations" putting a 15 in "Counts"

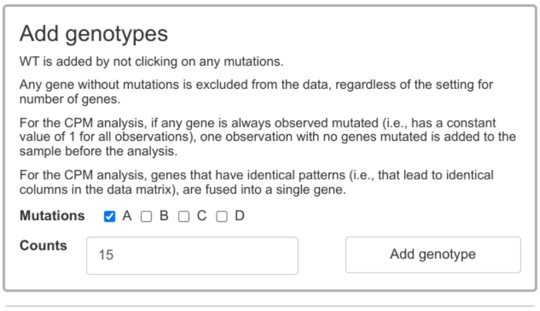

and then on "Add genotype". We next click on B and C in "Mutations" putting a 12 in Counts. We finally add 20 individuals with the "A, D" genotype (steps as before: click on A and D for mutations, and put a 20 in counts). After these steps, we can see the genotype composition as both a histogram and table:

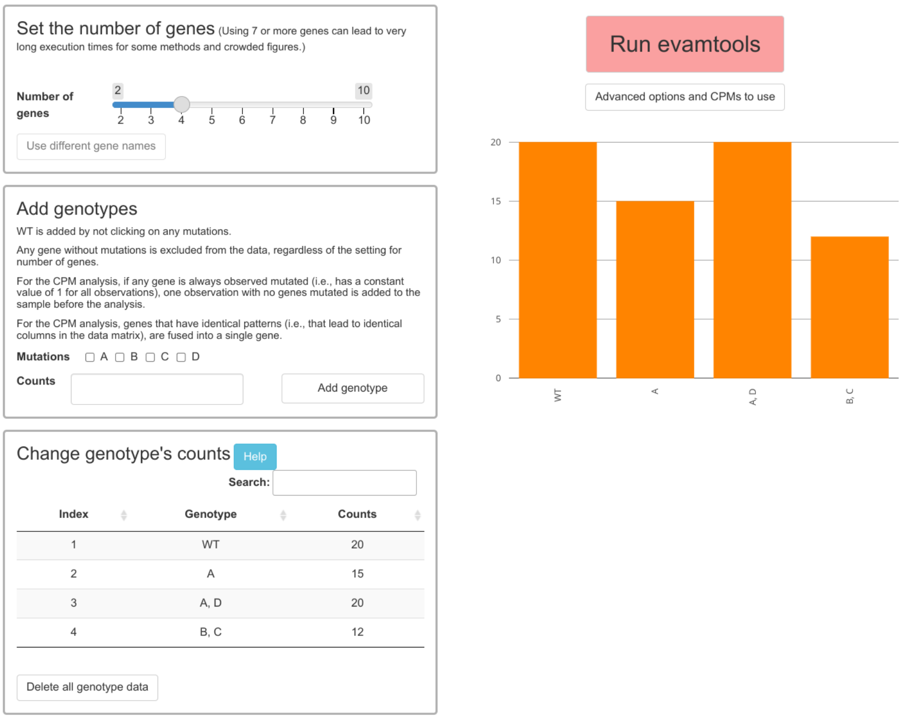

So that it is easier to go back to these data, we give this data set a name, for example "Aliased_1" under "Rename the data".

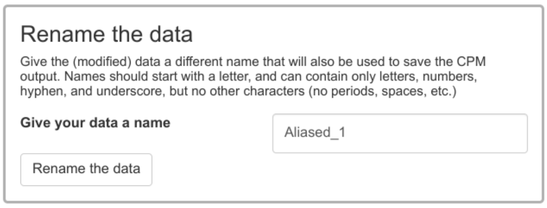

Click the "Rename the data" button, so the name is used and you will see it listed on the left, under “Examples and user’s data:”

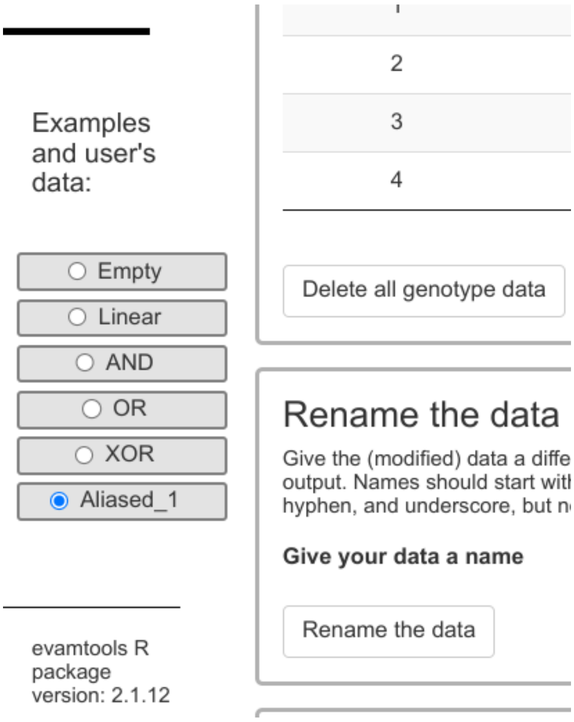

Now we click on “Run evamtools”. These are the fitted models (we did not use H-ESBCN here):

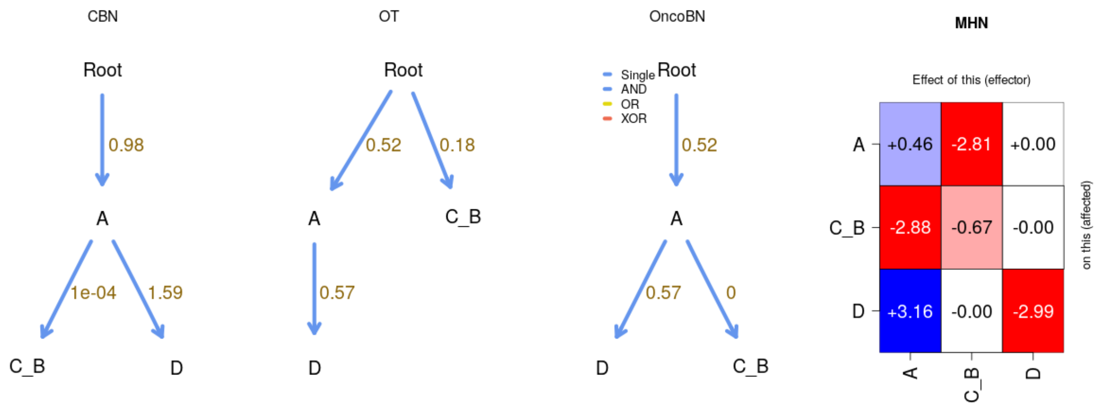

In the DAGs and the MHN log Θ matrix, as well as the transition probabilities, an event labelled “C_B”. “C_B” is the name of the event created automatically by EvAM-Tools by fusing the “C” and “B” events that are not distinguishable.

Note also that the models fitted by the DAGs, for example OncoBN or CBN, do not seem right when we look at the parameters of the “C_B” event. That is because there are no “A, B, C” events, and we would expect to see some if A on the one hand, and B and C on the other, are occurring independently. We will add some observations (eight, for example) with all of A, B, C. We go back to the “User input” tab, we “Rename the data” to “Aliased_2” (so that the new modifications we are about to make do not affect “Aliased_1”), and we add genoytpe “A, B, C” with a count of 8. And we click on “Run evamtools”. This is the output

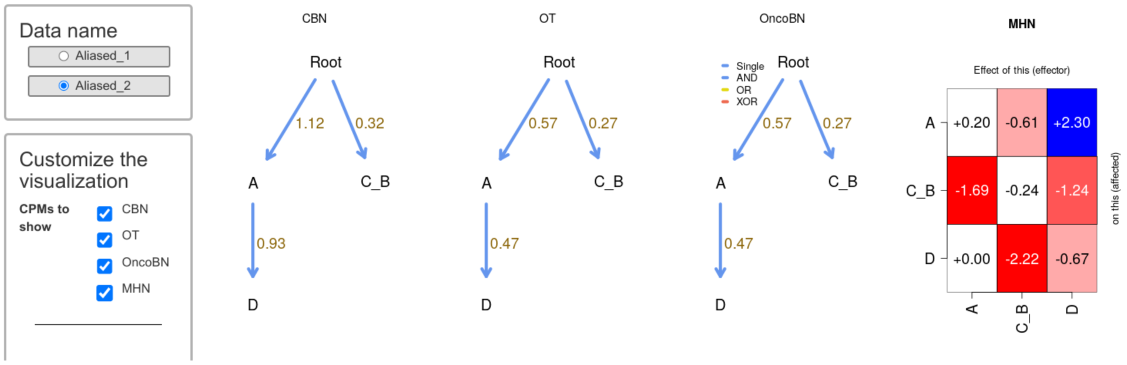

which is more sensible. Notice, however, that we still have “C_B” as an event because, in fact, they remain completely aliased, indistinguishable. This aliasing would be broken by adding just a single “C” or a single “B”, or a single “A, C”, or “A, B”, or “B, D” or “D, C” or any “A, B, D” or “A, C, D”.

### 4 Generating data from known models

The discussion in sections *"Analyzing the BRCA data set" (section 2.1)* and *"Analyzing the ovarian CGH data" (section 2.3)* has used CPMs on two cancer data sets for which the truth is unknown. To understand the differences between models, and the performance characteristics of different methods, we can simulate data under a known model and examine if the true pattern can be recovered. This is very easy to do with EvAM-Tools and addresses two commonly asked questions:

- Can we recover the true structure?
- How do different methods perform when data has been generated under the assumptions of another model?

This is what we will do below in examples *"A model with AND, XOR, OR" (section 4.2)* and *"A model with AND" (section 4.3)*. But EvAM-Tools is also useful to understand what different models imply in terms of the data we would observe, even without considering what each method would fit to a given observed data set; this is what we show in *"CPM models: what type of data they imply?" (section 4.1)*.

#### 4.1 CPM models: what type of data they imply?

##### 4.1.1 What happens if we increase $\epsilon$ for OncoBN?

We go to "User input" and, by default, the option "DAG and rates/cond. probs" is selected under "Generate cross-sectional data from CPM models". We can leave the default DAG in the selected "DAG_Fork_4". In "Type of model" under "Define DAG" we click on "OncoBN":

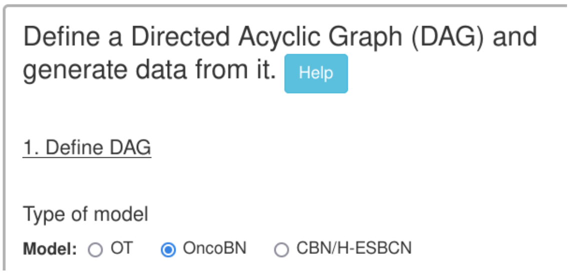

and you can see that the last column of the DAG table is now called "theta":

DAG table

Remember to hit Ctrl-Enter when you are done editing the DAG table for changes to take effect.

| From | To | Relation | theta |
|---|---|---|---|
| Root | A | Single | 0.5 |
| Root | B | Single | 0.7 |
| A | C | Single | 0.9 |
| B | D | Single | 0.3 |

Now, click on "Generate data from DAG"; to make patterns easier to observe, set the "Number of genotypes to sample” to a large number, such as 10000. If we look at the histogram we will see something similar to this one:

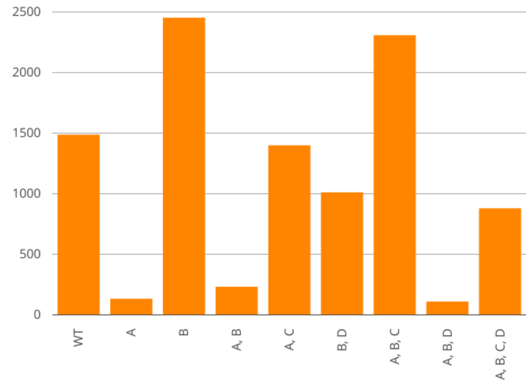

In particular, notice how the following genotypes are not observed: “A,D”, “B, C”, “B, C, D”, “A, C, D”, as they are not possible if the restrictions are completely respected. Now, increase “epos, $\epsilon$” to, say, 0.15, and click again on “Generate data from DAG”; now we will observe at least a few cases of all or most of the above four genotypes, as the predicted genotypes now incorporate deviations from the model.

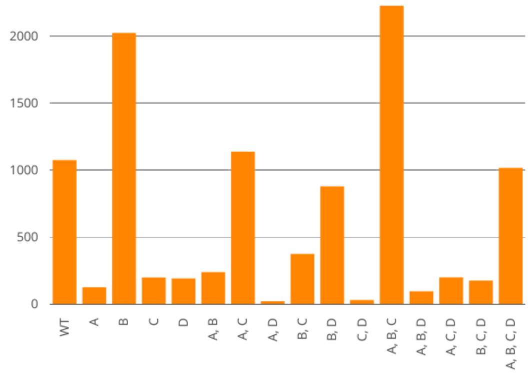

We could continue increasing the value of $\epsilon$; this will result in increasing frequencies of the above four genotypes, and a decrease in WT. Very large increases in $\epsilon$ will lead to a blurring of the signature of this DAG.

A more advanced exploration of the role of deviations from the model in OT and OncoBN compared to the role of observational noise would alter “epos, $\epsilon$” with and without simultaneously changing the value of “Observational noise”.

##### 4.1.2 A simple exploration of MHN

To try to gain a quick intuitive understanding of the multiplicative hazards model of MHN we go to “User input” and then click on “MHN log-Θ matrix”

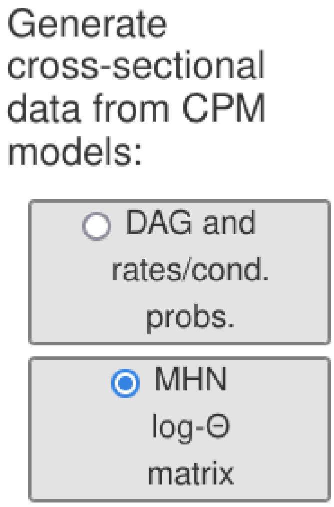

To start with a simple, yet not completely trivial, model, we set the number of genes to three:

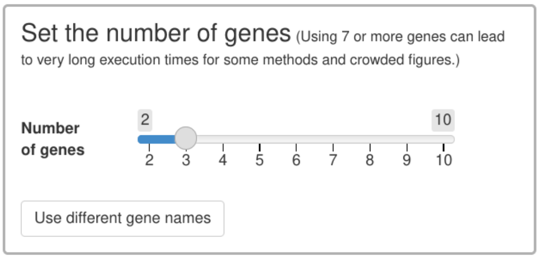

Now, we click on “Generate data from MHN model”:

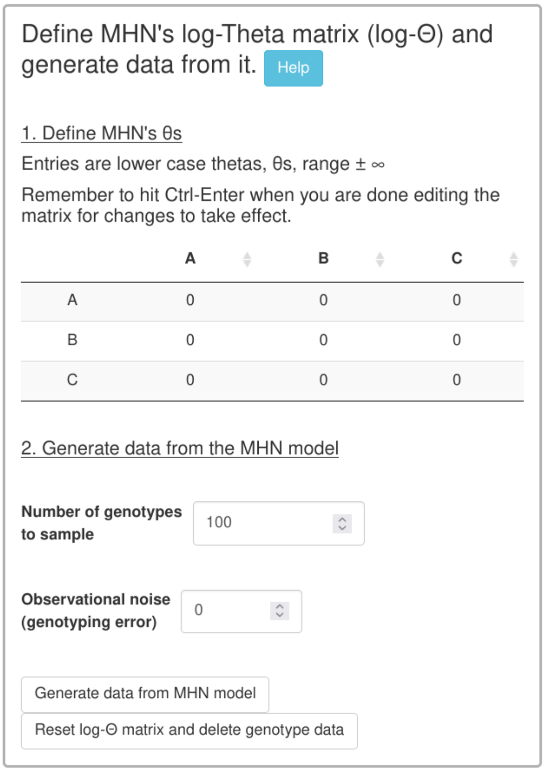

In this model, there are not multiplicative effects between genes. This is what we obtain (again, it might help to increase the Number of genotypes to sample to 10000 to decrease the role of random sampling noise and focus on the predicted genotype frequencies):

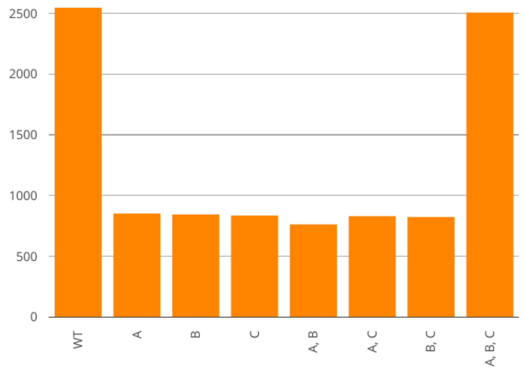

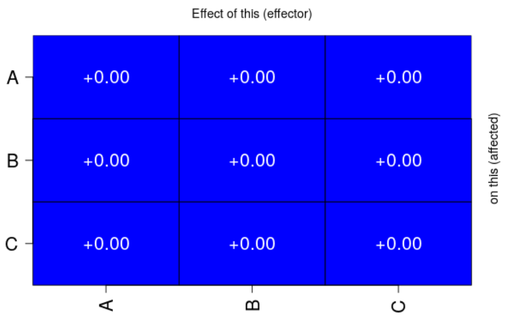

Let us now increase the baseline hazard of event B. For example, set log-$\Theta_{2,2}$ = 3. (Modify the entry, and click on Ctrl-Enter). The figure changes dramatically and all genotypes that have "B" have increased their frequency:

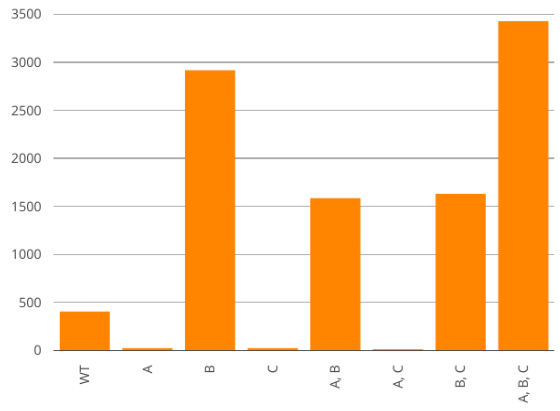

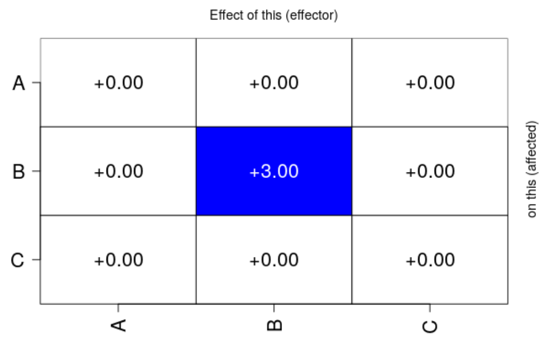

Let us now create a very large promoting effect of B on C; for that, we enter, for example, a 4 on the matrix entry (3, 2): log-$\Theta_{3,2} = 4$:

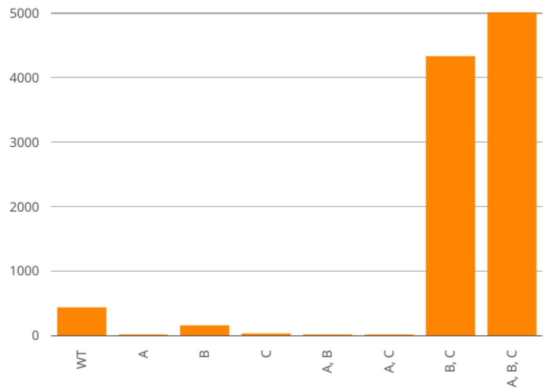

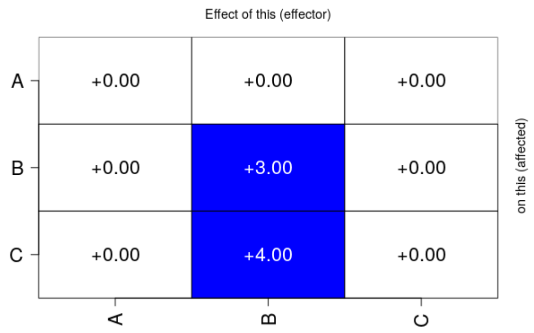

Notice how genotype B is more common than either A or C, but now whenever there is B it frequently in combination with C (much larger frequencies of genotypes B,C and A,B,C).

But what if B also an inhibiting effect on A? Set log-$\Theta_{1,2} = -2$:

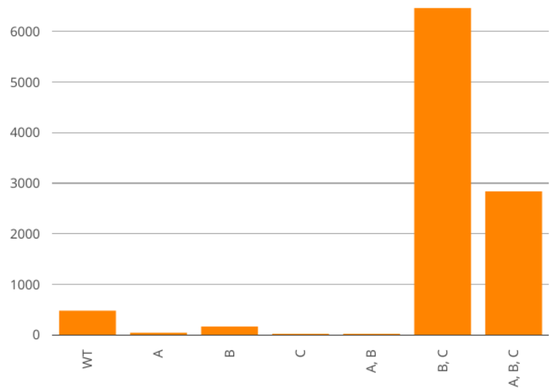

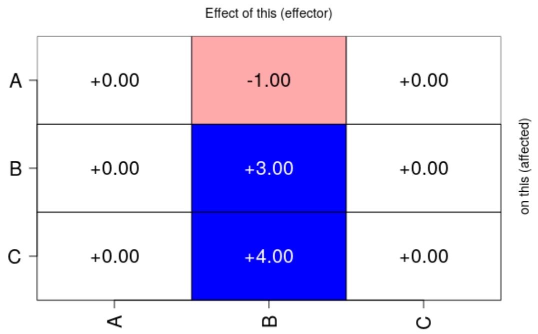

Notice how the frequency of genotype A,B,C has gone down (and also, though it is harder to appreciate, that of A,B).

#### 4.2 A model with AND, XOR, OR

Here, we will simulate data under a model that includes both AND, OR, and XOR relationships (e.g., H-ESBCN). EvAM-Tools, in its web app, already includes such a model for five genes:

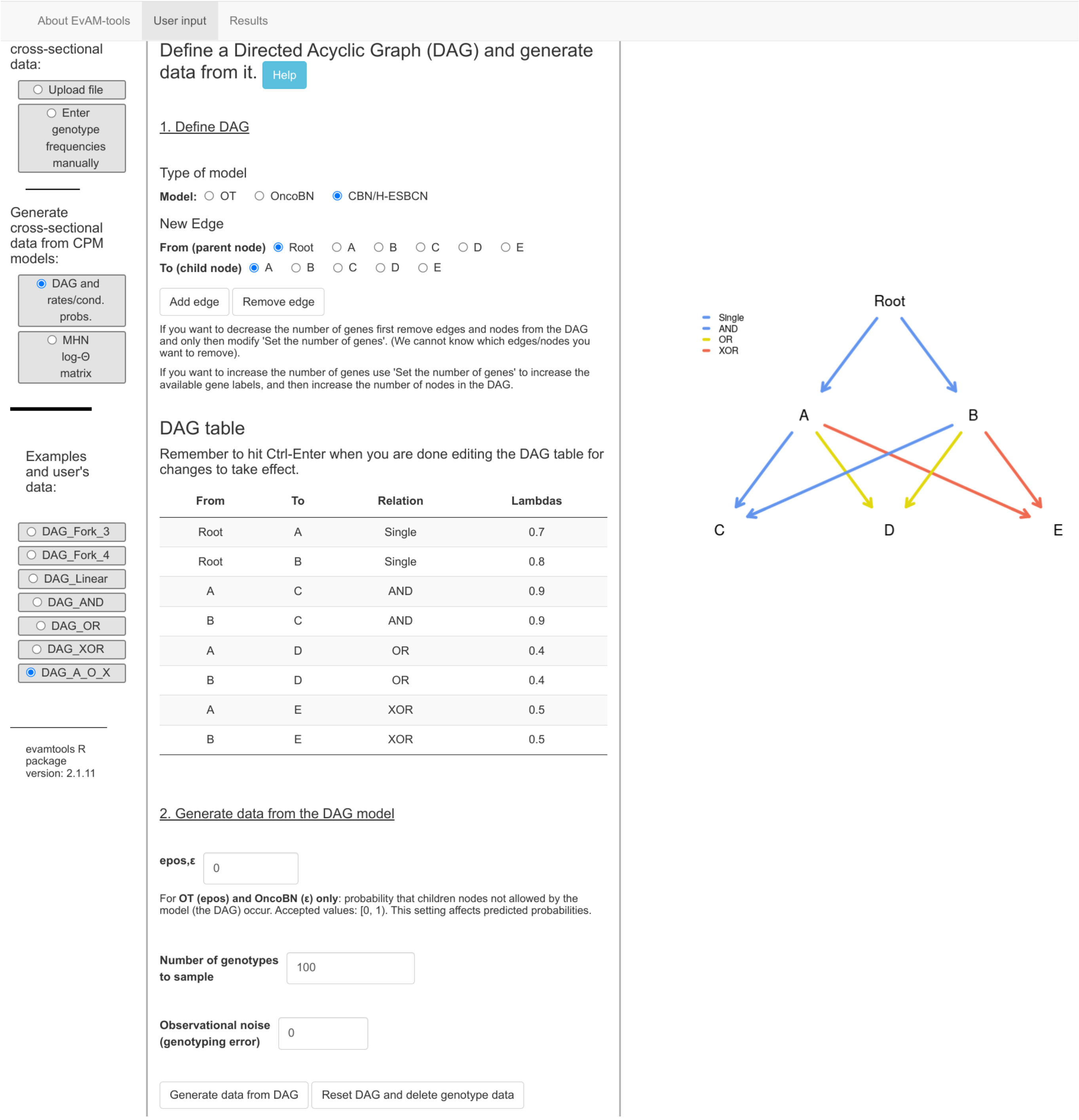

If we want, we can change values (rates, relationships, noise, etc). We will change the rates of A, B, D, and E, setting them to 3.2, 1.5, 2, and 1, respectively; we set the number of genotypes to sample to 1000 and we will add 1% of Observation noise (i.e., we will type 0.01 in the "Observational nose (genotyping error)" box). Then, we click on "Generate data from DAG", and obtain data simulated under that model:

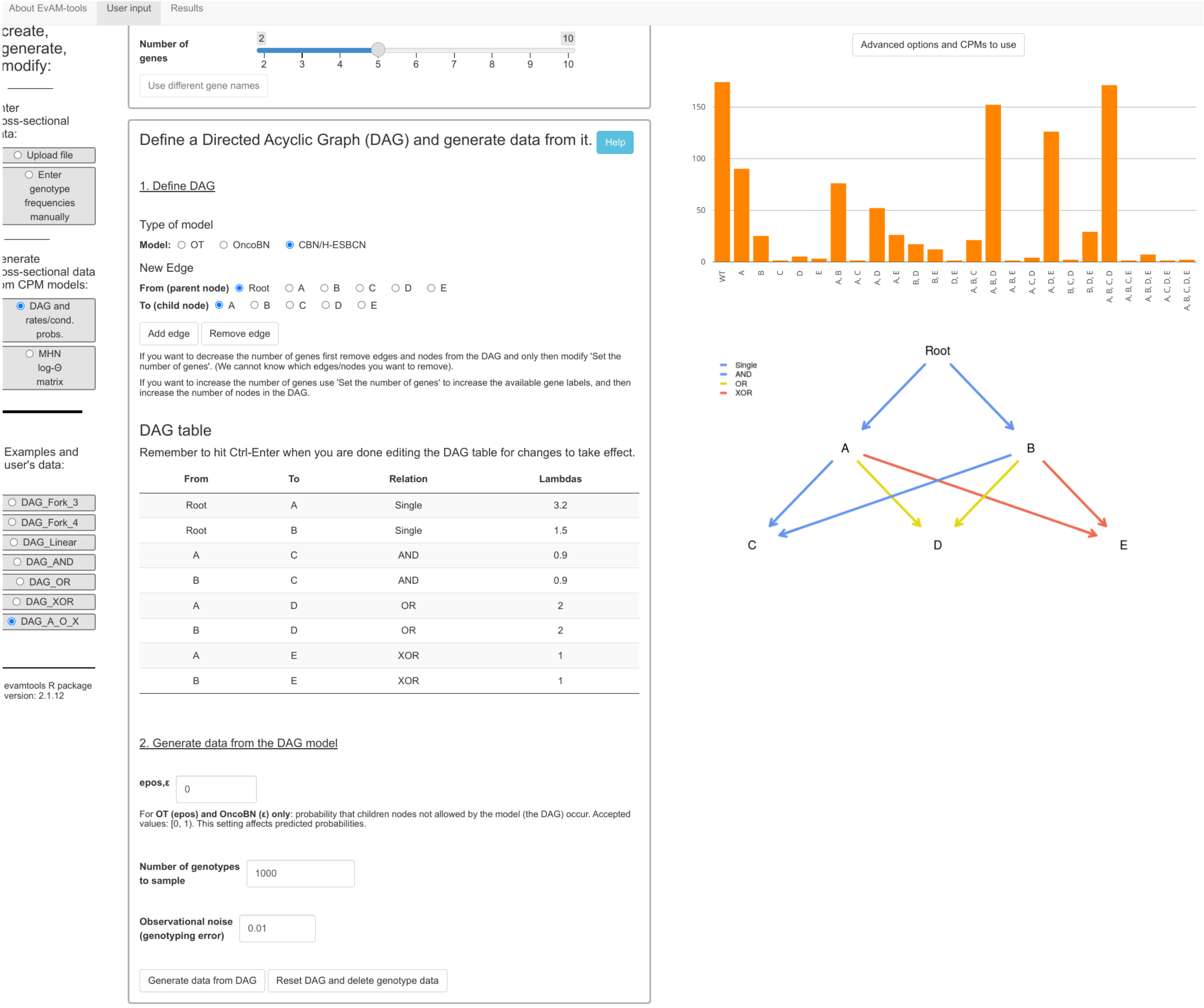

(Of course, the actual simulated data you are likely to obtain will differ from this one).

Now, we click “Run evamtools”, after adding H-ESBCN to the set of methods and setting its number of MCMC iterations to 500000 (again, this is done under “Advanced options and CPMs to use”). After about 30 seconds we obtain the output. In the plots below, and as we did before, we split it into three and two methods so it is easier to see. We show the models with two of the predictions: transition probabilities and predicted genotype relative frequencies (these predictions are also shown in the table, which we do not show below).

Note that the histograms of predicted genotype frequencies display, at most, the 20 most frequent genotypes (because of reasons of limited plotting space); all predicted genotypes are shown in the table.

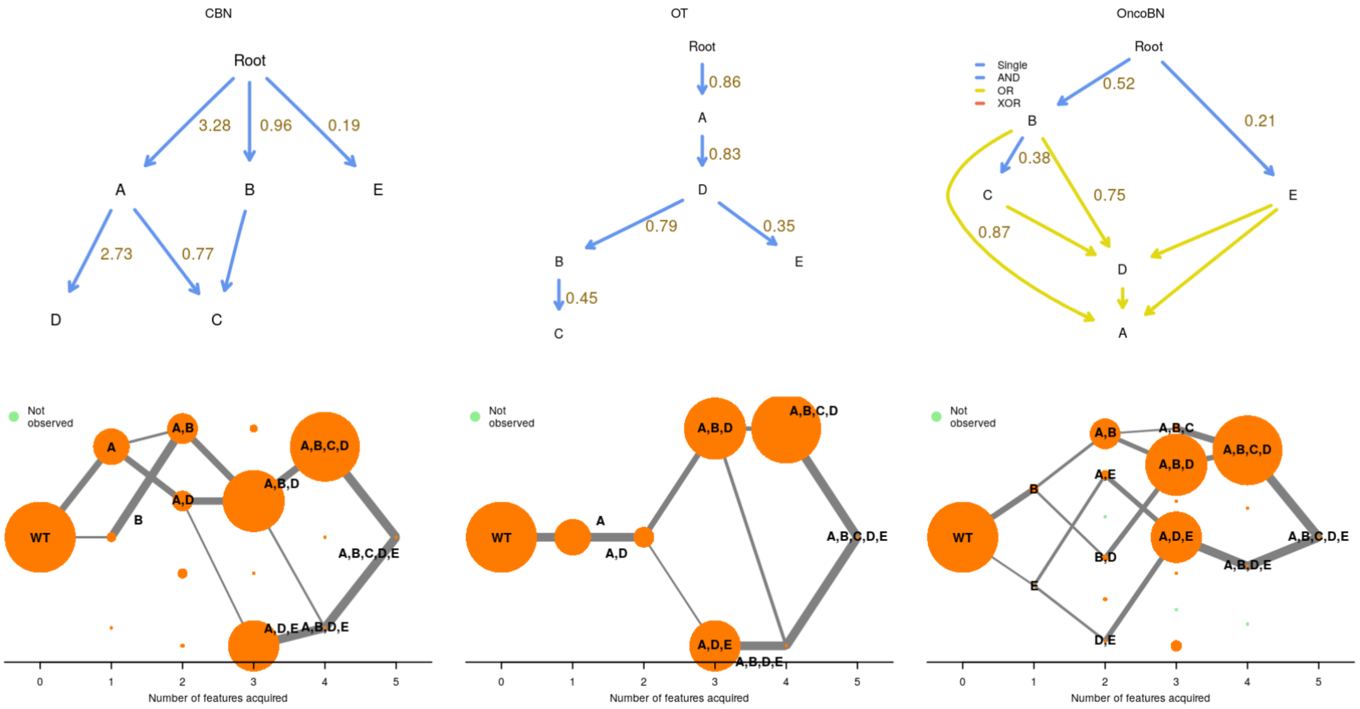
CBN
Root
3.28
0.96
0.19
A
B
E
2.73
0.77
D
C
OT
Root
0.86
A
0.83
D
0.79
0.35
B
E
0.45
C
OncoBN
Single
AND
OR
XOR
Root
0.52
0.21
B
0.38
0.75
C
E
0.87
D
A
Not observed
WT
A
B
A,B
A,D
A,B,D
A,B,C,D
A,B,C,D,E
A,D,E
A,B,D,E
Number of features acquired
A,B,C
A,E
B,D
E
D,E

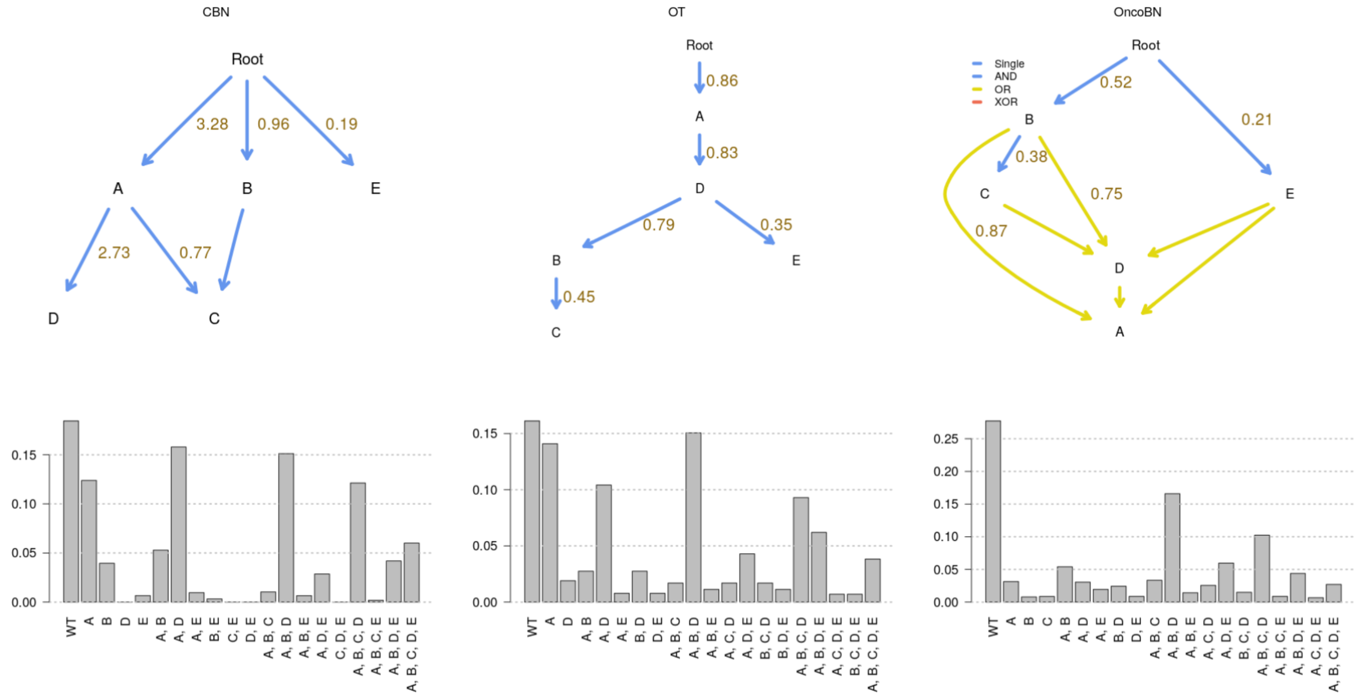
CBN
Root
3.28
0.96
0.19
A
B
E
2.73
0.77
D
C
OT
Root
0.86
A
0.83
D
0.79
0.35
B
E
0.45
C
OncoBN
Single
AND
OR
XOR
Root
0.52
0.21
B
0.38
0.75
C
E
0.87
D
A

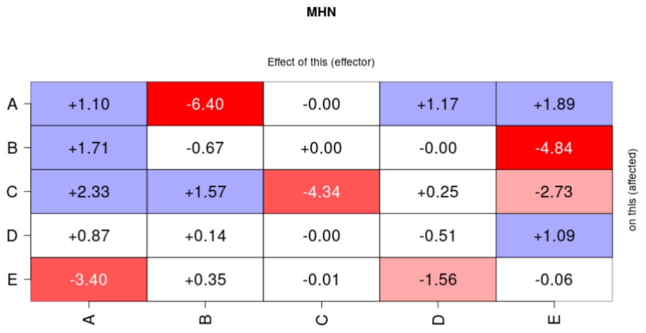

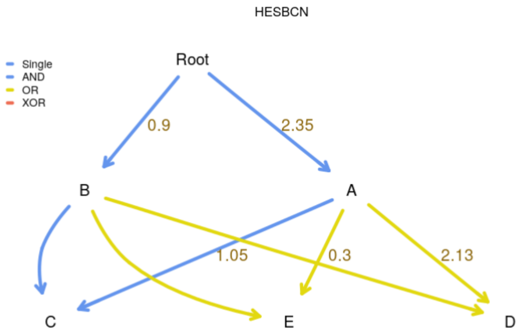

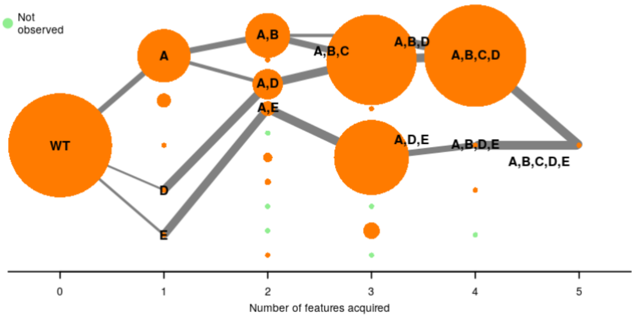

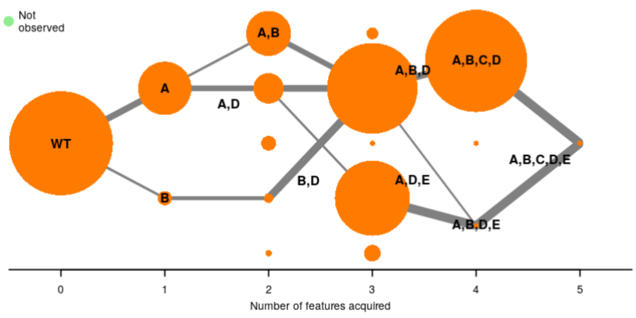

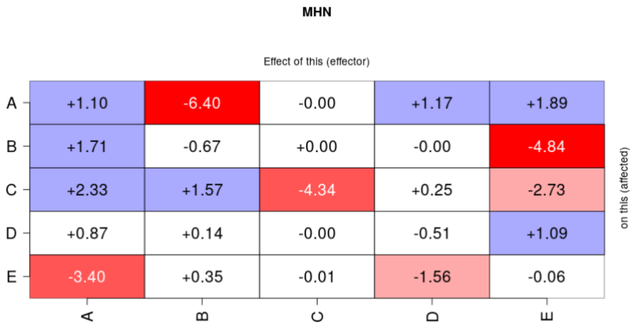

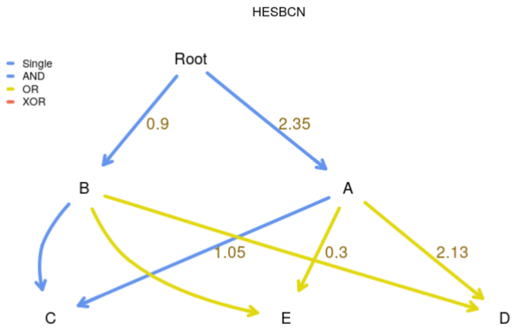

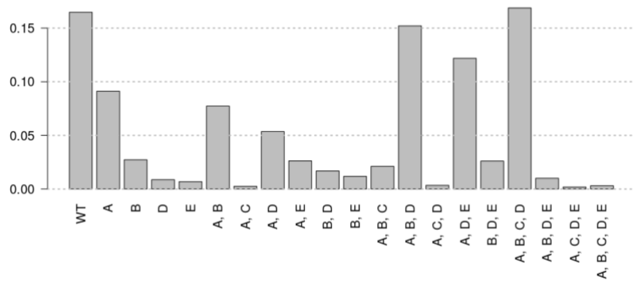

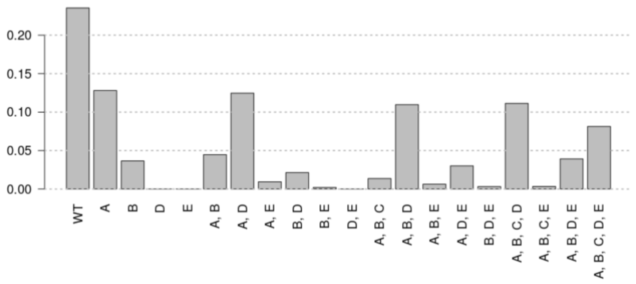

And, as has been the case before, repeated runs of H-ESBCN can lead to different results, for example:

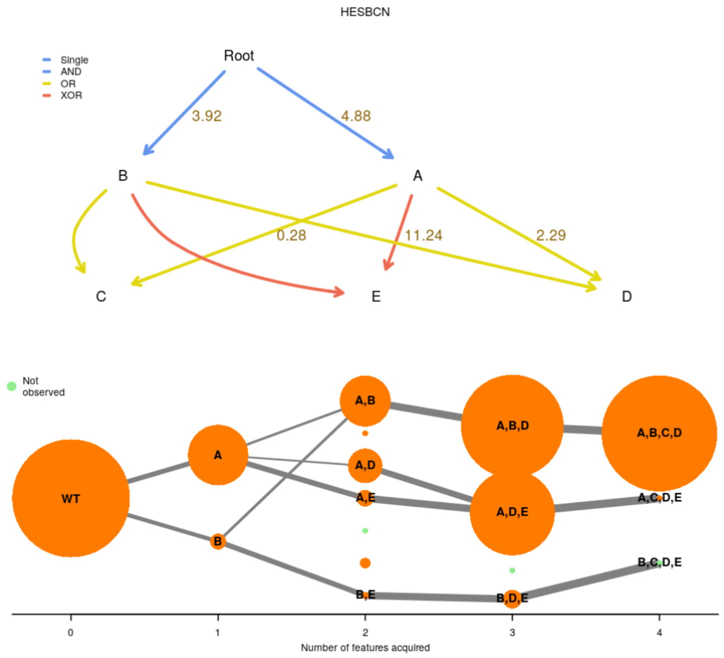

None of OT, CBN, or OncoBN can capture XOR relationships. But in this case, H-ESBCN incorrectly infers an XOR for D (it is really an OR) and an OR for C (it is really an AND).

We could increase the sample size by 10 by just setting “Number of genotypes to sample” to 10000 or add different levels of noise, etc.

We could also ask to get, as return, not only the predicted genotype relative frequencies, but sampled genotype counts with, possibly, some noise added. Even without noise added, the relative frequencies of the sampled genotype counts would differ from the predicted genotype relative frequencies just because of sampling noise. We can do that by going back to the “User input” tab, clicking on “Advanced options and CPMs to use” and setting “Sample genotypes” to TRUE and selecting the number of samples, which here we set equal to the sample size of original data set (i.e., 1000); we also set the observation noise to 0.01.

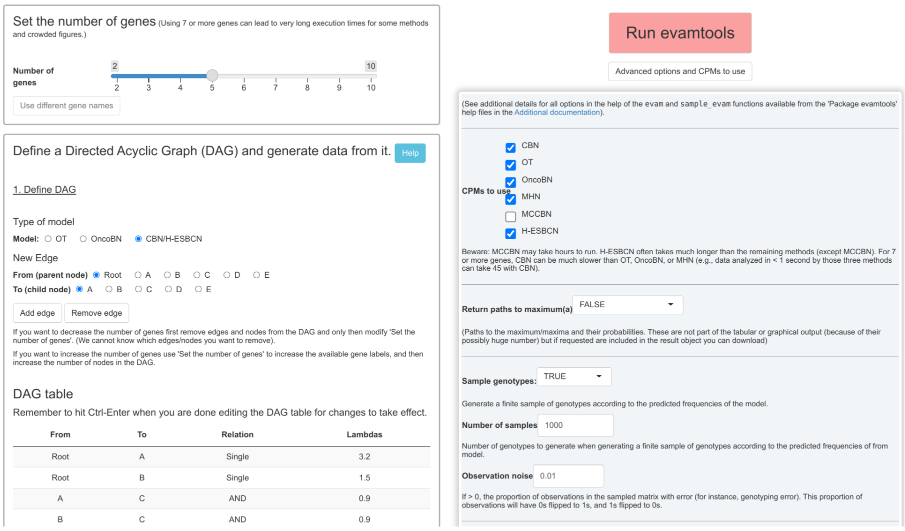

We hit on "Run evamtools" and, as before, we get the output but we now have one extra possible "Predictions from fitted models to display":

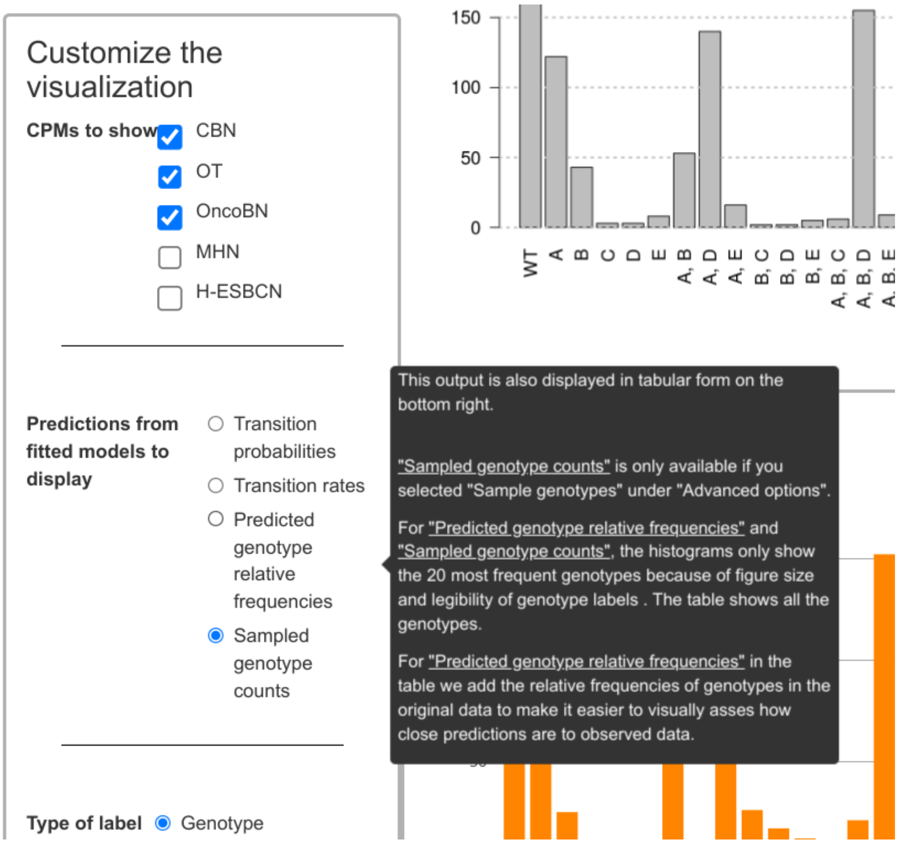

And this is the output for three of the models; notice the added variability (i.e., how the relative heights and even the actual genotypes present are not the same as in the predicted genotype counts):

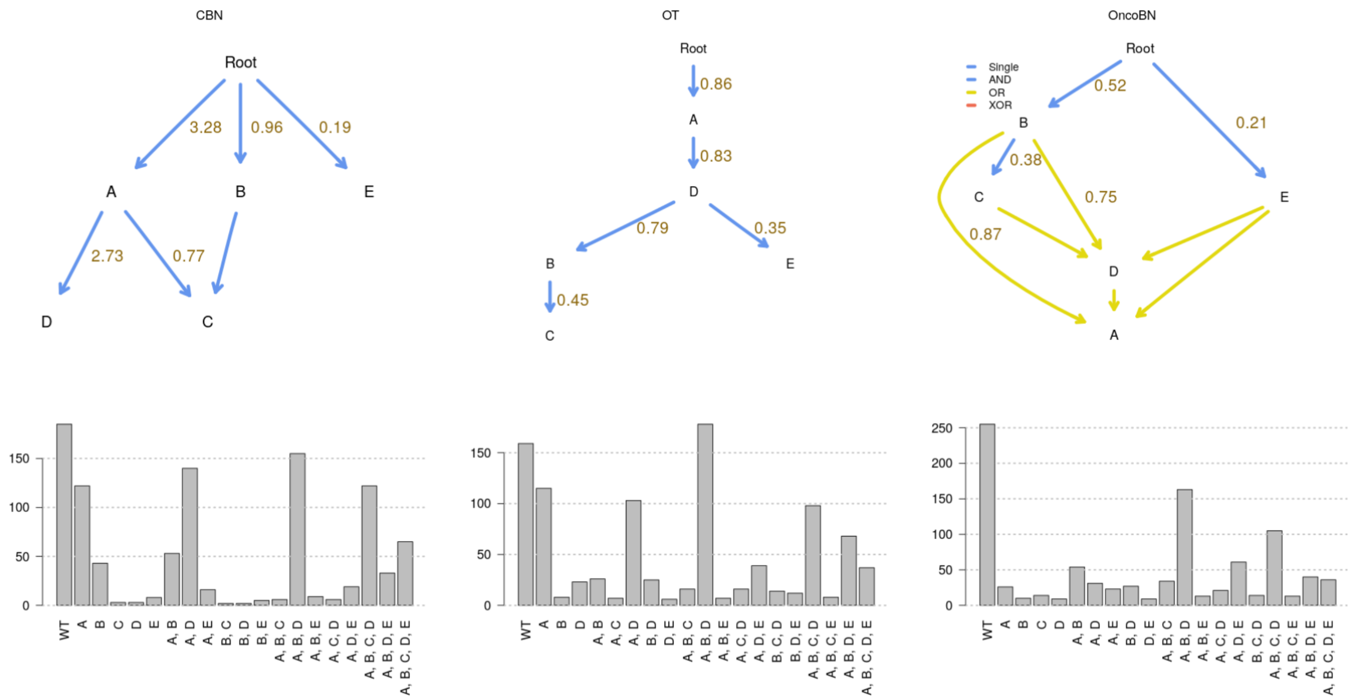

Of course, you can switch from displaying “Sampled genotype counts” to displaying “Predicted genotype counts” just by clicking on the button on the left.

##### 4.3 A model with AND

Let us use now a model with AND; we use the preselected "DAG_AND", with 1000 observations and a 5% (0.05) genotyping error. Remember to click on "Generate data from DAG" after changing the noise level:

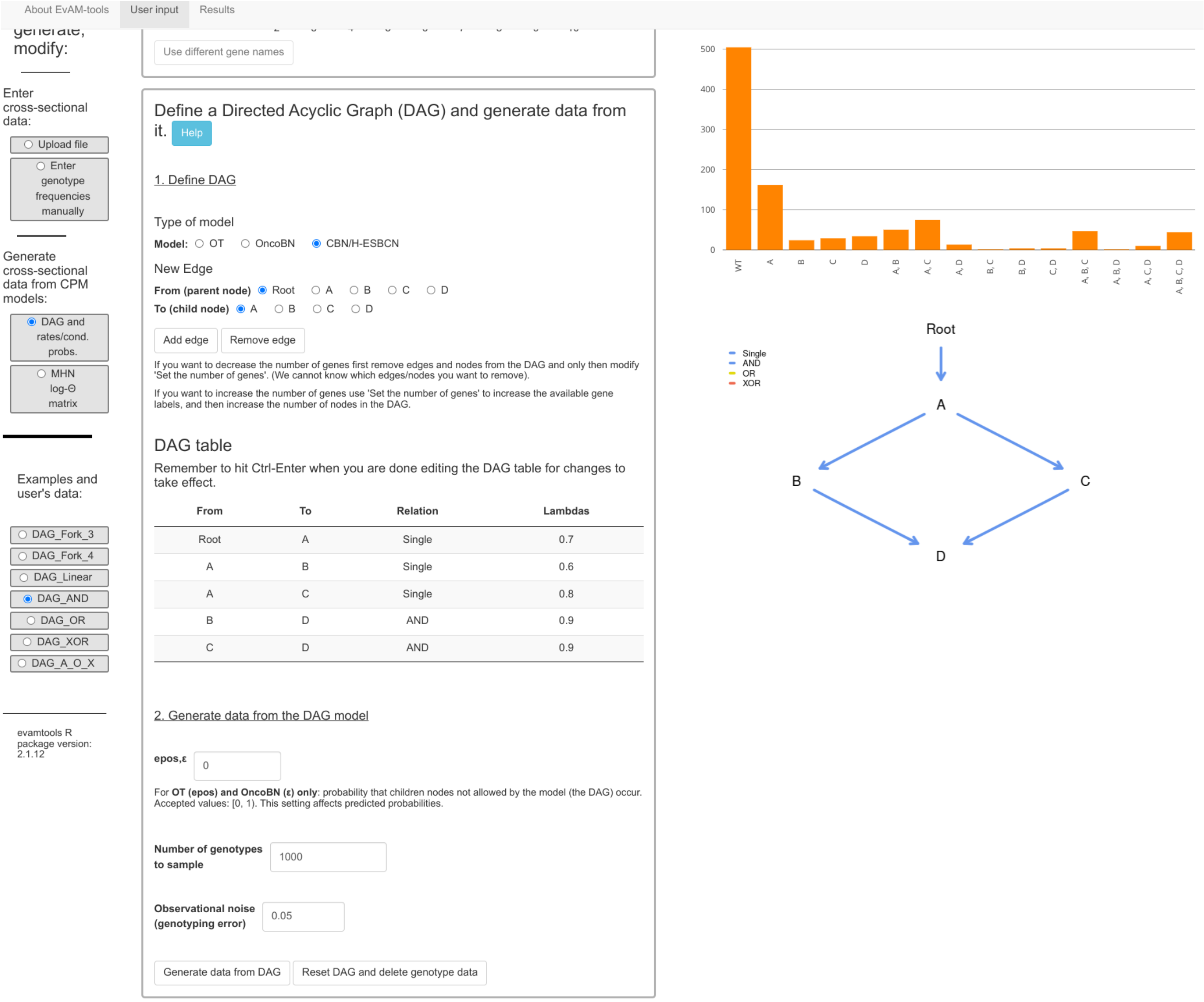

These are the fitted models:

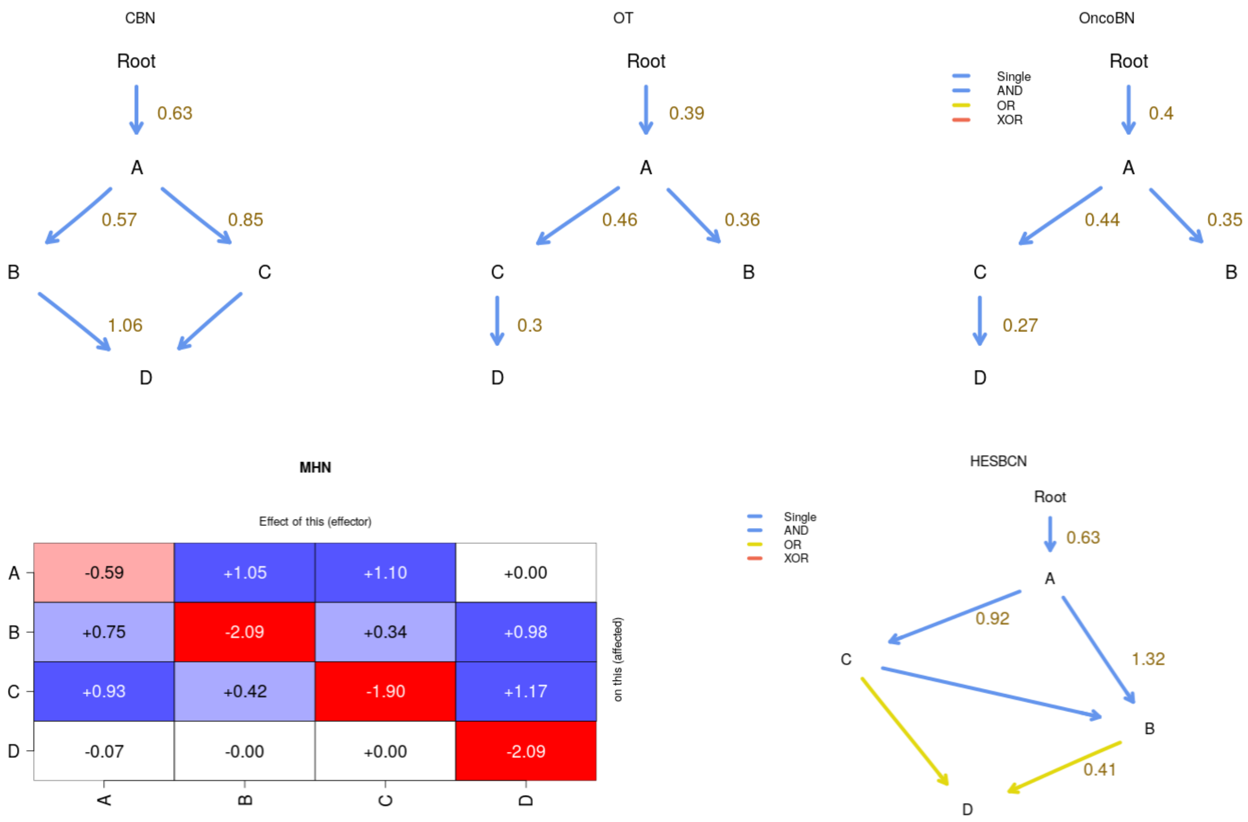

H-ESBCN incorrectly infers the AND as an OR. CBN correctly infers the underlying model and provides estimates of the parameters that are very close to the true ones. OT and OncoBN cannot infer the correct true dependencies: OT because it cannot fit DAGs, but only trees (i.e., each node has only one parent) and OncoBN because it was run in disjunctive mode (OR relationships). We can run OncoBN using the conjunctive model; we have done this before (sections *"Analyzing the BRCA data set", section 2.1* and *"Analyzing the ovarian CGH data", section 2.3*) and it involves going to "Advanced options and CPMs to run" and setting, for "OncoBN options", the "Model"

to “Conjunctive”:

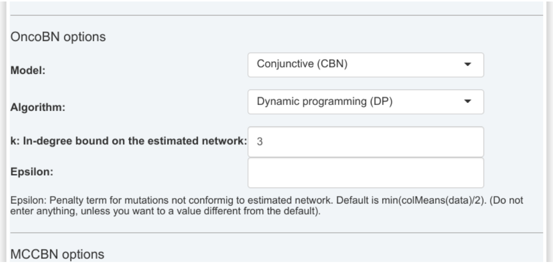

This is the output we get:

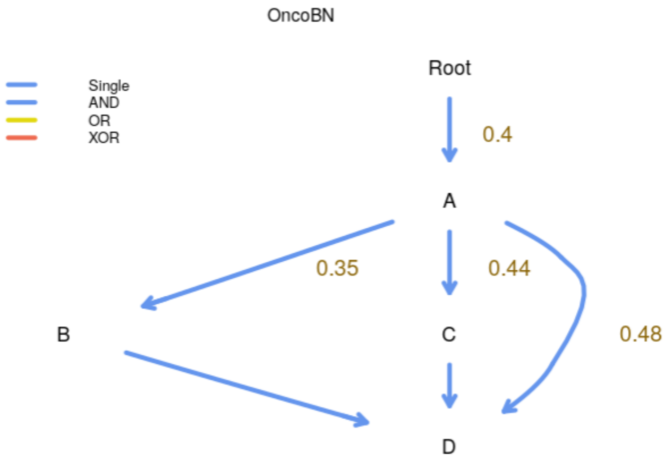

This correctly identifies the joint dependency of D on both B and C, but there is also and edge between A and D that we would never observe with CBN (as CBN always returns the transitively reduced DAGs); this is a technical issue beyond the scope of this document, but one that we have discussed in the OncoBN repo[3].

For H-ESBCN the returned model also includes an edge, the one between A and B, that is not necessary: since B depends on C and C depends on A, having B depend with an AND on both A and C is not needed (i.e., the transitive reduction of that DAG would remove the edge from A -> B). Note also that removing the A -> B arrow does not affect the relationships with D. Of course, for the relationships between C, B, and D we should not return the transitive reduction (as that would break the OR of D on either C or B). Thus, a model without the $A \rightarrow B$ would generate the exact same predictions as the model returned by H-ESBCN. (Why does this happen with H-ESBCN? It is a consequence of the heuristic search over DAG structures, which can occasionally return these topologies).

#### 4.4 Modifying data generated from a CPM model before analysis

What if the data came from a given model but some additional process had altered the genotype data? For example, suppose data really came from a CBN model, but the frequency of WT genotypes is too small because the data have been filtered to contain only genotypes with at least one driver mutation, or the data contain some contaminated samples that would never had

[3] https://github.com/phillipnicol/OncoBN/issues/5

become tumores and have an excess of WT. We can explore this by generating data from a CPM model and, then, modifying the genotype composition, as we did in section *"Is it just sample size?" (section 2.2)* when we modified uploaded data.

As an example, go to "User input", "DAG and rates/cond. probs", and use the default selected "DAG_Fork_4". Change lambdas to 2, 2.5, 1, 1.5, for A, B, C, D, respectively. Add 1% of observational noise. And set the "Number of genotypes to sample" to 5000, to ensure small sample sizes are not the culprit of different inferences. Click on "Generate data from DAG" and, for simplicity, then analyze the data with OT, OncoBN, CBN, MHN, and H-ESBCN.

This is the output:

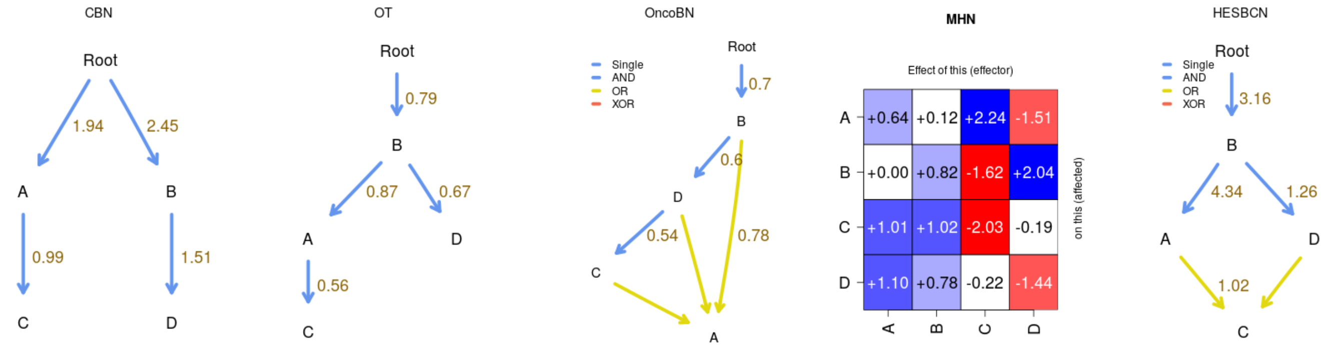

CBN correctly infers the DAG and the estimates of the $\lambda$s are close to the true ones. The other methods are not able to infer this model correctly.

Now, go back to the "User input". Before modifying the data, and to keep a copy of the originally generated one, on the "Rename data" type a name (e.g., "data_original") and click on "Rename data". Now, enter a new name in "Rename data", for example "data_few_wt", click on "Rename data", and modify the WT frequency; I change it from its original value of 912 to 9. And then, analyze the data clicking on "Run evamtools". This is the output:

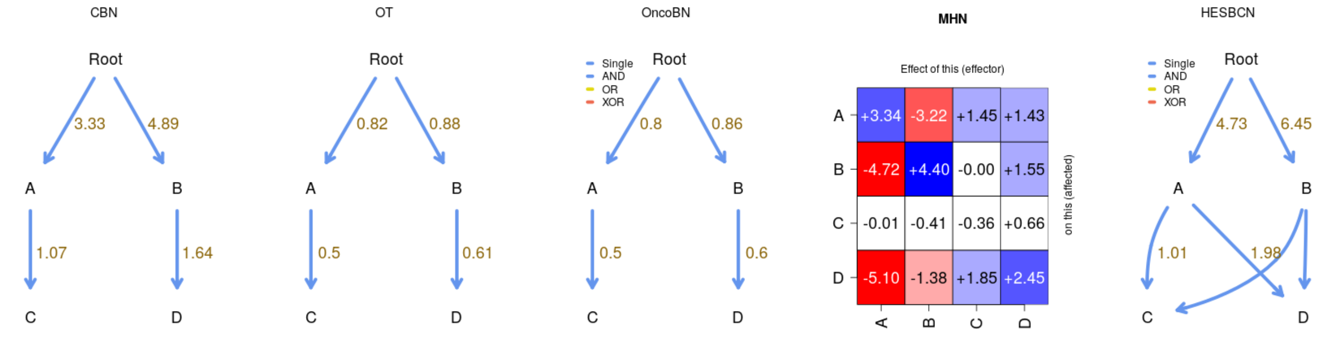

OT and OncoBN get the structure right, and for CBN the main consequence is altering the rates of A and B, increasing them (which is what we would expect). MHN also has increased estimates of log-$\Theta_{A,A}$ and log-$\Theta_{B,B}$ when we reduced the frequency of WT.

What if we removed the WT completely? We go back (and, if it is not the selected one, click, on the left, under "Examples and user's data", on "data_original"), and set WT to 0 (possibly after creating a new data set, “data_no_wt”) and analyze them:

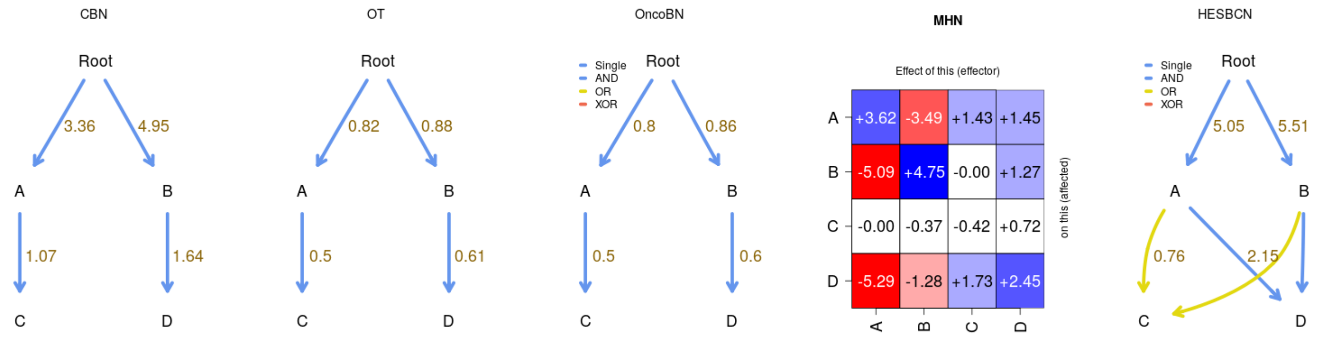

The effect is minor, which is not surprising, since the large cut in WT had been going from 912 to 9.

We could, instead, eliminate all the observations with the four mutations (e.g., maybe genotypes with three driver mutations are already generally lethal to the organism?). We go to “User input”, select, on the left, “data_original”, rename it (“no_all_mut”), and set to 0 the A,B,C,D genotype.

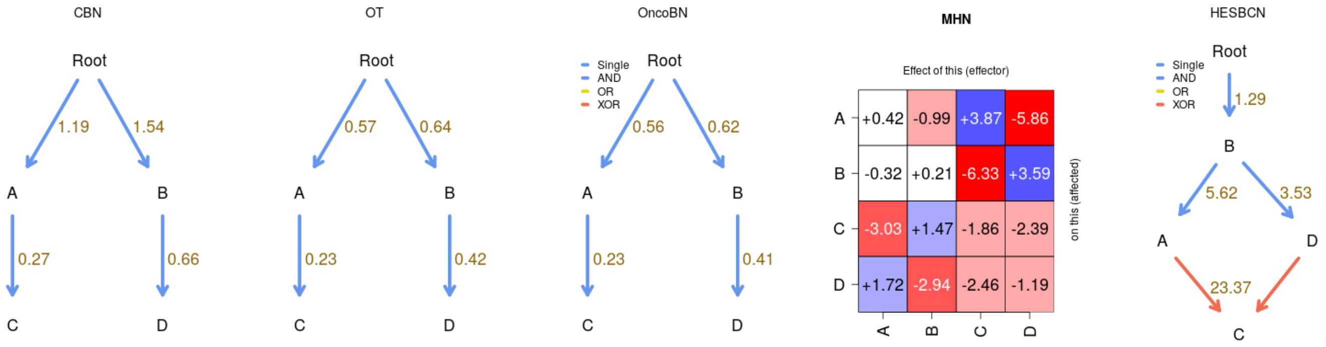

The DAG structures for CBN, OT, OncoBN are preserved, but now H-ESBCN has modeled an XOR; the XOR models is not actually present, but unless there are XOR or similar phenomena, models with AND and OR cannot model the absence of A,B,C,D, given the relatively high frequencies of the rest of the genotypes (the CBN model, for example, has decreased the estimates of all the $\lambda$s).

We could also, as mentioned at the beginning of this section, increase the number of WTs. Etc, etc. We will not pursue this any further here. The key message from this section is that EvAM-Tools makes it very simple to examine targeted, specific, deviations in genotype composition from the genotype composition generated by a CPM model.

### 5 Simulating random CPMs/evams

If we were interested in systematically examining the performance of the different methods under different models, simulating random CPM (or evam) models is crucial. This type of work (generating and analyzing large numbers of simulations) is not suited for a web app, but it can be easily done with the R package. The key function here is `random_evam`.

```
## Load the package
library(evamtools)
## For reproductibility
set.seed(3)
he_r1 <- random_evam(ngenes = 5, model = "HESBCN")
he_r1$HESBCN_model

##   From To      edge   Lambdas Relation
## 1 Root  A Root -> A 0.6656892   Single
## 2 Root  B Root -> B 1.1189358   Single
## 3    A  C    A -> C 1.8736265   Single
## 4    A  D    A -> D 2.0159447      XOR
## 5    A  E    A -> E 1.6987091   Single
## 6    B  D    B -> D 2.0159447      XOR

## Now, simulate a data set of size 200 from that model
## with 5% genotyping error

he_s1 <- sample_evam(he_r1, N = 200, obs_noise = 0.05)

## Analyze this data with all the methods except MCCBN (for speed)

he_s1_anal <- evam(he_s1$HESBCN_sampled_genotype_counts_as_data,
                   methods = c("CBN", "OT", "OncoBN",
                               "HESBCN", "MHN"))

## Show the fitted DAGs
he_s1_anal[grepl("_model", names(he_s1_anal))]

## $OT_model
##   From To      edge OT_edgeBootFreq OT_edgeWeight OT_obsMarginal
## A Root  A Root -> A              NA     0.3148883          0.345
## B Root  B Root -> B              NA     0.4069873          0.430
## D Root  D Root -> D              NA     0.3040531          0.335
## E    A  E    A -> E              NA     0.5851968          0.225
## C    E  C    E -> C              NA     0.9163967          0.210
##   OT_predMarginal
## A       0.3450000
## B       0.4300000
## D       0.3350000
## E       0.2244513
## C       0.2102331
##
```

```
## $CBN_model
##   From To       edge init_lambda final_lambda rerun_lambda CBN_edgeBootFreq
## A Root  A Root -> A    0.433470     0.438862     0.438863               NA
## B Root  B Root -> B    0.678177     0.677041     0.677041               NA
## C    A  C    A -> C   15.918836     2.134728     2.134609               NA
## D Root  D Root -> D    0.454241     0.457503     0.457502               NA
## E    C  E    C -> E    2.020337     8.245700     8.247622               NA
##
## $MCCBN_model
## [1] NA
##
## $OncoBN_model
##   From To       edge     theta Relation
## 2 Root  B Root -> B 0.4300000   Single
## 3 Root  C Root -> C 0.2100000   Single
## 1    C  A    C -> A 0.9285714   Single
## 4    B  D    B -> D 0.5045045       OR
## 5    C  D    C -> D 0.5045045       OR
## 6    C  E    C -> E 0.7857143   Single
##
## $OncoBN_fitted_model
## [1] "DBN"
##
## $HESBCN_model
##   From To       Edge    Lambdas Relation
## 1 Root  A Root -> A   1.028580   Single
## 2 Root  B Root -> B   1.097880   Single
## 3    A  C    A -> C 122.697000   Single
## 4    A  D    A -> D   7.867050      XOR
## 5    B  D    B -> D   7.867050      XOR
## 6    C  E    C -> E   0.564528       OR
## 7    D  E    D -> E   0.564528       OR
```

```
## Show MHN
he_s1_anal["MHN_theta"]
```

```
## $MHN_theta
##       A     B     C     D     E
## A -0.52  0.61  0.00 -2.48  0.00
## B -2.30 -0.23 -2.33  1.47 -0.01
## C  3.42 -0.31 -3.04 -0.24  0.00
## D  0.19 -3.53  1.23 -0.44  0.00
## E  1.31 -0.44  2.19 -0.32 -2.13
```

This is just one example; serious simulation studies would examine exhaustively a range of scenarios. And we could, of course, compare other output, such as the predicted genotype frequencies, or the probabilities of paths to the maximum (use argument `paths_max = TRUE` when calling `evam`), etc. But this should be enough to show you how EvAM-Tools can be used to systematically compare the performance of different methods in different scenarios.

### 6 Appendix: getting the BRCA and Ov data sets from the R console

Here we show how to obtain, from the R console, the two data sets used in section *"Analysis of cross-sectional data" (section 2)*. We simply export those data in a CSV format that can be uploaded to the web app.

```r
## Load the package to access the BRCA data
library(evamtools)
data(every_which_way_data)

## You can check the names here, which are the same
## as in Suppl File S5 Text of Diaz-Uriarte & Vasallo, 2019
##  names(every_which_way_data)

write.csv(every_which_way_data[["BRCA_ba_s"]],
          file = "BRCA_ba_s.csv", row.names = FALSE,
          quote = FALSE)

## Now export the ovarian cancer CGH data
library(Oncotree)

## Loading required package:  boot

data(ov.cgh)

## Rename column names: they start with a number and
## finish on a "+" (gain) or "-" (loss), so automatically
## reading these column names removes the +/- and adds an
## "X" as the first character. Let us have columns start
## with L or G for loss/gain.

new_cn <- stringi::stri_sub(colnames(ov.cgh), 1L, 2L)
new_cn <- paste(ifelse(grepl("+", colnames(ov.cgh), fixed = TRUE),
                            "G", "L"), new_cn, sep = "")
ov2 <- ov.cgh
colnames(ov2) <- new_cn
write.csv(ov2,
          file = "ov2.csv", row.names = FALSE, quote = FALSE)
```

### 7 License and copyright

This work is Copyright, ©, 2022, Ramon Diaz-Uriarte.

Like the rest of this package (EvAM-Tools), this work is licensed under the GNU Affero General Public License. You can redistribute it and/or modify it under the terms of the GNU Affero General Public License as published by the Free Software Foundation, either version 3 of the License, or (at your option) any later version.

This program is distributed in the hope that it will be useful, but WITHOUT ANY WARRANTY; without even the implied warranty of MERCHANTABILITY or FITNESS FOR A PARTICULAR PURPOSE. See the GNU Affero General Public License for more details.

You should have received a copy of the GNU Affero General Public License along with this program. If not, see https://www.gnu.org/licenses/.

The source of this document and the EvAM-Tools package is at https://github.com/rdiaz02/EvAM-Tools.
