## Supplementary material for "EvAM-Tools: tools for evolutionary accumulation and cancer progression models": EvAM-Tools methods' details and FAQ

2022-10-03
Version 44b7274

**9 License and copyright 35**

**10 References 35**

### 1 Introduction

This document provides additional details about EvAM-Tools and the methods included in it. Another file, evamtools_examples.pdf, from https://rdiaz02.github.io/EvAM-Tools/pdfs/evamtools_examples.pdf, includes commented examples, with both real and simulated data, that illustrate the use and utility of EvAM-Tools.

You can run the web app from https://iib.uam.es/evamtools/ or download a Docker image from https://hub.docker.com/r/rdiaz02/evamshiny; to run the R package download a Docker image from https://hub.docker.com/r/rdiaz02/evamrstudio.

### 2 Cancer Progression Models included in EvAM-Tool: details

#### 2.1 Cancer Progression Models and cross-sectional data: overview and type of input data

In cross-sectional data a single sample is obtained from each subject or patient. That single sample represents the "observed genotype" of, for example, the tumor of that patient. Genotype can refer to single point mutations, insertions, deletions, or any other genetic modification; in fact, these models have been used to analyze point mutations, gains and losses of CGH regions, SNP alterations, pathway alteration data, etc: the granularity of the data and level of analysis depend on the question addressed, and is not inherent to the models. As is often done by Cancer Progression Models (CPM) software, we think of the cross-sectional data as being stored in a matrix, where rows are patients or subjects, and columns are genes/CGH regions/SNPs/pathways/etc; the data is a 1 if the event (or alteration or mutation) was observed and 0 if it was not.

We have used expressions such as "genotype", "mutation" and other genetic- and genomic-related terms, but nothing prevents CPMs from being used with non-genetic, non-genomic data, and thus our preference for the expression "event accumulation models". The key features that the data must have to be properly analyzed with these methods are: a) that events or alterations are (or can be reasonably assumed to be) gained one by one; b) that once gained, they are not lost (e.g., there is no back mutation); c) that we can consider the different individuals/patients in the cross-sectional data as replicate evolutionary experiments or runs where all individuals are under the same constraints (e.g., genetic constraints if we are dealing with mutations); see further details below (section *2.2, "Cancer Progression Models (CPMs): assumptions"*).

Cancer progression models (CPMs) or, more generally, event accumulation models, use these cross-sectional data to try to infer restrictions in the order of the irreversible accumulation of discrete events; for example, that a mutation on gene B is always preceded by a mutation in gene A (maybe because mutating B when A is not mutated results in a lethal state for that cell). Inferring restrictions, in the sense just explained (B only if A), is what CBN, OT, OncoBN, and H-ESBCN do. Other cancer progression models, such as MHN, instead of modeling deterministic restrictions, model promoting/inhibiting interactions between genes, for example that having a mutation in gene A makes it very likely to gain a mutation in gene B.

#### 2.2 Cancer Progression Models (CPMs): assumptions

CPMs model the irreversible accumulation of discrete events. They assume that the observations in the cross-sectional data set are independent realizations of evolutionary processes where the same constraints hold for all tumors; therefore, a cross-sectional data set is considered a set of replicate evolutionary experiments where all individuals are under the same (genetic) constraints (Gerstung *et al.*, 2011; Beerenwinkel *et al.*, 2015, 2016; Diaz-Uriarte and Vasallo, 2019). The objective of CPMs is to infer these constraints. CPMs assume that events are gained one by one (no simultaneous acquisition of events) and that there is no back mutation so that once gained an event is not lost; CPMs also assume that the events that drive the process (driver genes if we are thinking about cancer) are known and present in the data set. Finally, CPMs assume that all subjects start the evolutionary process without any of the studied events (i.e., all subjects start the process with 0s in the matrix of subjects by alterations). If we think about cancer, this means that "CPMs assume that all tumors start cancer progression without any of the mutations considered in the study (the above matrix of subjects by driver alterations), but other mutations could be present that have caused the initial tumor growth" (Diaz-Uriarte and Vasallo, 2019); these other additional mutations that lead to the initiation of the process are absorbed in the root node from which cancer starts (Attolini *et al.*, 2010).

#### 2.3 Cancer Progression Models (CPMs): details

##### 2.3.1 Oncogenetic Trees (OT)

OTs are among the earliest formal models of accumulation of mutations in cancer. They were originally described in Desper *et al.* (1999) (see also Simon *et al.*, 2000; Radmacher *et al.*, 2001); additional references include Szabo and Boucher (2008); Szabo and Pappas (2022); Szabo and Boucher (2002). With OTs, restrictions in the accumulation of mutations (or events) are represented as a tree[1]. Hence, a parent node can have many children, but children have a single parent: therefore, an event can only directly depend on another event. As for all CPMs that use DAGs and trees, an edge from gene $i$ to gene $j$ means that a mutation in $i$ must occur before a mutation in $j$ can occur; an edge (or arrow from $i$ to $j$) indicates a direct dependency of a mutation in gene $j$ on a mutation in gene $i$.

OTs are untimed models (in contrast to, for example CBN, explained in section 2.3.3): weights along edges (the $\pi_{xy}$ we will use below) can be directly interpreted as probabilities of transition along the edges by the time of observation (Szabo and Boucher, 2008, p. 5). In other words, edge weights represent conditional probabilities of observing a given mutation, when the sample is taken, given the parents are observed.

As explained in (Szabo and Boucher, 2008, Definition 1, p. 4): "A pure untimed oncogenetic tree is a tree $T$ with a probability $\pi(e)$ attached to each edge $e$. This tree generates observations on mutation presence/absence the following way: each edge $e$ is independently retained with probability $\pi(e)$; the set of vertices that are still reachable from $M_0$ [the root of the tree, representing no alterations] gives the set of the observed genetic alterations."

To give an example, suppose a tree as follows:

[1] A tree is a Directed Acyclic Graph (DAG) where a child node can have only only one parent. Thus, trees are DAGs, but not all DAGs are trees.

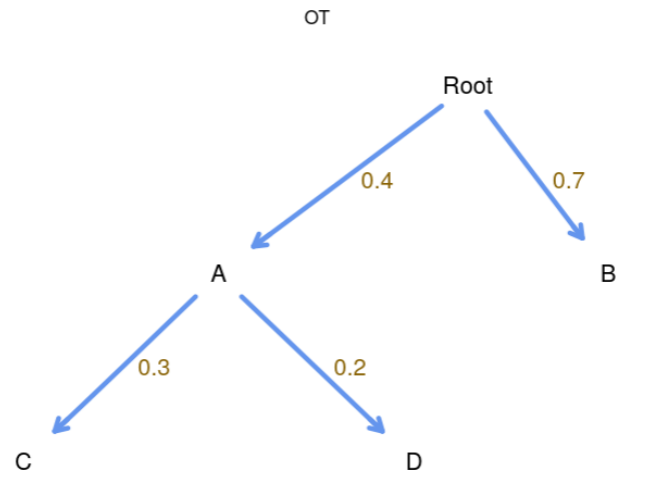

In this tree, events (mutations) A and B can be acquired independently, and depend on no one (Root is $M_0$ in the notation of Szabo and Boucher, 2008). C and D depend on A (and are independent of each other, conditional on A). The parameters of the model, shown in brown, are:

- Probability of acquiring $A$, $\pi_{0A} = 0.4$; $\pi_{0A}$ is the notation in Szabo and Boucher, 2008, and is the *weight* along the edge from $M_0$ (Root) to $A$.
- Probability of acquiring $B$, $\pi_{0B} = 0.7$.
- Probability of acquiring $C$, given $A$ has already been acquired, $\pi_{AC} = 0.3$ (again, $\pi_{AC}$ is the *weight* along the edge from $A$ to $C$).
- Probability of acquiring $D$, given $A$ has already been acquired, $\pi_{AD} = 0.2$.

According to the above model, the tumor develops as follows: starting from Root (or $M_0$), the tumor can gain $A$ and $B$, and these are independent events. If $A$ is gained, then the tumor can gain $C$ and $D$, and these two are again independent events (once $A$ has been gained). Therefore, the probabilities of the different genotypes or states of the tumor at the time of sampling are:

- Only Root or $M_0$, i.e., no events gained (i.e., only genotypes without any mutation, or "WT"): $(1 - \pi_{0A})(1 - \pi_{0B})$.
- Only $A$ occurs (i.e., genotype $A$): $\pi_{0A}(1 - \pi_{0B})(1 - \pi_{AC})(1 - \pi_{AD})$.
- Only $B$ occures (i.e., we observe genotype $B$): $\pi_{0B}(1 - \pi_{0A})$.
- Both $A$ and $B$ (but no $C$ or $D$), genotype $AB$: $\pi_{0A}\pi_{0B}(1 - \pi_{AC})(1 - \pi_{AD})$.
- $A$ and $C$, genotype $AC$: $\pi_{0A}(1 - \pi_{0B})\pi_{AC}(1 - \pi_{AD})$.
- Both $B$ and $C$ but no $A$, genotype $BC$: 0 (as $A$ needs to occur before $C$ can occur).
- ...

The above describes the ideal scenario, without errors. OT includes a model for errors from different sources: deviations from the model (i.e., events that do not respect the pure untimed model above) and observational (e.g., genotyping) errors. Together, these two types of error cause false positive and false negative observational errors $(\epsilon_+, \epsilon_-)$. These error rates are estimated by the OT algorithm and are incorporated in the computation of the predicted frequencies of genotypes according to OT (see details in *"CPMs: Error models", section 2.3.6*).

When using OT, as explained in Szabo and Boucher (2008, p. 5), the main objective is reconstructing the topology of the tree; the estimation of the edge probabilities (the weights or $\pi_{xy}$) and the error rates ($\epsilon_+, \epsilon_-$) is of secondary importance. As detailed in Szabo and Boucher (2008, p. 5), the estimation of the topology uses an "(...) algorithm [that] takes a greedy bottom-up approach: it assigns the parent of each node by finding the maximum-weight in-edge starting from the leaves." and that provides a computationally fast way of inferring the tree. The full algorithm for topology reconstruction is provided in Szabo and Boucher (2008, Section 3 and Fig. 2) (the algorithm is also provided in Figure 2 of file `ot.pdf`, part of the documentation of Szabo and Pappas, 2022); estimation of the weights is detailed in Szabo and Boucher (2008, p. 13). Sufficient conditions for the reconstruction of the true tree when there are false positive and false negative errors are given in Szabo and Boucher (2002) and sample size requirements in Szabo and Boucher (2008, p. 8)(see also Desper *et al.*, 1999).

##### 2.3.2 OncoBN

OncoBN, described in Nicol *et al.* (2021), is similar to OT in the sense of being an untimed oncogenetic model but, in contrast to OT, a node can have multiple parents (again, as for all CPMs that use DAGs and trees, an edge from gene $i$ to gene $j$ means that a mutation in $i$ must occur before a mutation in $j$ can occur; an edge —or arrow from $i$ to $j$— indicates a direct dependency of a mutation in gene $j$ on a mutation in gene $i$). When there are multiple parents the relationships and models can be of two different kinds:

- disjunctive (OR relationship): the DBN, Disjunctive Bayesian Network model;
- conjunctive (AND relationship): the CBN, Conjunctive Bayesian Network model.

A given OncoBN be either a DBN or a CBN, but not both: it can have conjunctive or disjunctive relationships, but not both. (And note that the CBN models fitted by OncoBN are untimed, and thus the parameters do not have the same interpretation as the parameters of the CBN models discussed below, *"Conjuntive Bayesian Networks (CBN)", section 2.3.3*).

As explained in Nicol *et al.* (2021, p. 2), a key difference between the conjunctive (AND) and the disjunctive (OR) model is that under the conjunctive model all parent alterations that constitute the AND relationship must be present in a cell for the child mutation to occur; the disjunctive model, in contrast, allows child event to occur when just one of the parent events has taken place. According to the authors, this might make the model better for modeling intra-tumor heterogeneity.

The following DAG shows a conjunctive model fitted with OncoBN:

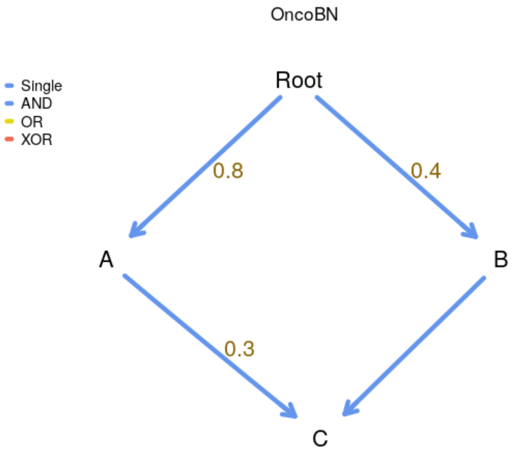

.

Note that the value of 0.3 is the value of the parameter $\theta_C$: this is the conditional probability of $C$ given its ancestors. (So, in contrast to OT, but similar to CBN and H-ESBCN, the parameters are not of edges, but of events). The values of $\theta$ are: $\theta_A = 0.8$, $\theta_B = 0.4$, $\theta_C = 0.3$. According to the OncoBN model the probabilities of the different genotypes are:

- Only Root (i.e., only genotypes without any mutation, or "WT"): $(1 - \theta_A)(1 - \theta_B)$.
- Only $A$, i.e., genotype $A$: $\theta_A(1 - \theta_B)$.
- $A$ and $C$, genotype $AC$: 0, since acquiring C requires also B.
- $A$ and $B$ (but not $C$), genotype $AB$: $\theta_A\theta_B(1 - \theta_C)$.
- All of $A$, $B$, $C$, genotype $ABC$: $\theta_A\theta_B\theta_C$.
- Only $C$: 0, since neither $A$ nor $B$ have occurred.
- ...

The next DAG is identical, except the model is a disjunctive one (notice the edges are OR edges):

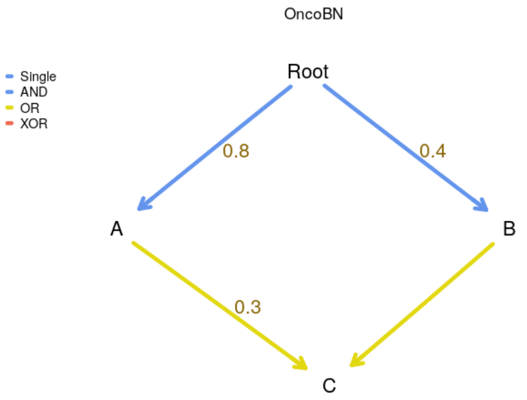

.

Now, $\theta_C$ is the probability of $C$ occurring if at least one of its ancestors has occurred. Therefore, we have the following probabilities of genotypes, where those that differ from the conjunctive case have been marked in bold with an initial asterisk:

- Only Root (i.e., only genotypes without any mutation, or "WT") : $(1 - \theta_A)(1 - \theta_B)$.

- *** Only $A$: $\theta_A(1-\theta_B)(1-\theta_C)$.**
- *** $A$ and $C$, genotype $AC$: $\theta_A(1-\theta_B)\theta_C$.**
- $A$ and $B$ (but not $C$), genotype $AB$: $\theta_A\theta_B(1-\theta_C)$.
- All of $A$, $B$, $C$, genotype $ABC$: $\theta_A\theta_B\theta_C$.
- Only $C$: 0, since neither $A$ nor $B$ have occurred.

The above represent the probabilities in a model without errors. OncoBN includes an error model, the "spontaneous activation model", where there is a non-zero probability of observing child events when restrictions in the DAG are not satisfied. The rate of spontaneous activation is part of the estimation procedure, and is included in the computed probabilities of the different genotypes (see details in section *2.3.6, "CPMs: Error models"*). For example, under the disjunctive model above, the probability of observing genotype $C$ would be $(1-\theta_A)(1-\theta_B)\epsilon$, where $\epsilon$ is the spontaneous activation probability, which is set as the same for all events (Nicol *et al.*, 2021, p. 5). (Figure 1 of Nicol *et al.*, 2021 provides another example of the role of $\epsilon$ in computing predicted probabilities).

For structure and model parameter learning (Nicol *et al.*, 2021, p. 5)), and to avoid overfitting and increase interpretability of the models, the authors use as the score the Bayesian Information Criterion (BIC), so the log-likelihood is penalized by the number of edges ($\log(N)|E|$, where $|E|$ the number of edges); in addition, the search space is restricted to DAGs with an in-degree bound that, by default, is set to 3 (this option can be changed in EvAM-Tools; in the web app, under "Advanced options", "OncoBN options", "k: In-degree bound of the estimated network"). Finally, for disjunctive models, there is an additional step of removing low confidence edges that are likely to be spurious.

To search for the best graph (the model with the best BIC) the authors provide two algorithms: an exact procedure that uses dynamic programming and an approximate structure learning algorithm that uses genetic programming (Nicol *et al.*, 2021, p. 5)). The authors recommend the dynamic programming procedure for less than 30 events, and the genetic algorithm for larger problems. (By default, EvAM-Tools uses the dynamic programming algorithm; this can be changed, for example in the web app, under "Advanced options", "OncoBN options", "Algorithm").

##### 2.3.3 Conjuntive Bayesian Networks (CBN)

In terms of the representation of the restrictions, CBN, like OncoBN, generalizes the tree-based restriction of OT to a directed acyclic graph (DAG): a node can have multiple parents. A node with multiple parents means that all of the parents have to be present (all of the parent events must have occurred) for the children to appear; therefore, relationships are conjuntive —AND relationships between the parents (recall OncoBN can model AND and OR relationships). CBN also differs from OT and OncoBN because the CBN model is a timed model: the $\lambda$s, the parameters of the models, are the rates of the exponentially distributed times to fixation of an event given that all parents of that event have been observed (i.e., given that the event restrictions, as specified in the DAG, are satisfied: Montazeri *et al.*, 2016, p. i729; Gerstung *et al.*, 2009, section 2.2).

Specifically, $T_i$, the waiting time for event $i$ to occur, is an exponentially distributed random variable with parameter $\lambda_i$ conditioned on all the parent mutations, $\text{pa}(i)$, having occurred (Gerstung *et al.*, 2009); thus, $T_i$ is defined recursively as (Gerstung *et al.*, 2009; Hosseini *et al.*, 2019):

$$T_i \sim \text{Exp}(\lambda_i) + \max_{j \in \text{pa}(i)} T_j \tag{1}$$

To give an example, suppose a DAG as follows:

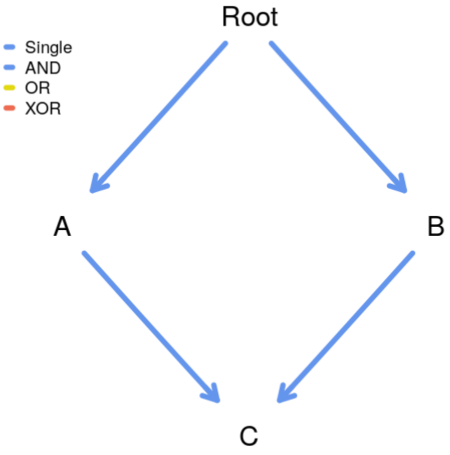

.

Then, the time to fixation of the three mutations (not genotypes) are:

- $T_A \sim \text{Exp}(\lambda_A)$
- $T_B \sim \text{Exp}(\lambda_B)$
- $T_C \sim \text{Exp}(\lambda_C) + \max(T_A, T_B)$

and we will not observe $C$ unless both $A$ and $B$ have occurred.

The $\lambda$ parameters of the CBN model define the transition rate matrix between genotypes (see also Montazeri *et al.*, 2016). For the example above we have:

- Rate from WT to genotype with A mutated: $\lambda_A$.
- Rate from WT to genotype with B mutated: $\lambda_B$.
- Rate from genotype with A mutated to genotype with both A and B mutated: $\lambda_B$.
- Rate from genotype with B mutated to genotype with both A and B mutated: $\lambda_A$.
- Rate from genotype with A and B mutated to genotype with A, B, C mutated: $\lambda_C$.

In other words, this is the transition rate matrix, where only genotypes that can appear are shown (i.e., genotypes $C$, $AC$, and $BC$ are not shown):

$$Q = \begin{array}{c} \\ WT \\ A \\ B \\ AB \\ ABC \end{array} \begin{array}{c} \begin{array}{ccccc} WT & A & B & AB & ABC \end{array} \\ \begin{pmatrix} -(\lambda_A + \lambda_B) & \lambda_A & \lambda_B & 0 & 0 \\ 0 & -\lambda_B & 0 & \lambda_B & 0 \\ 0 & 0 & -\lambda_A & -\lambda_A & 0 \\ 0 & 0 & 0 & -\lambda_C & \lambda_C \\ 0 & 0 & 0 & 0 & 0 \end{pmatrix} \end{array} \tag{2}$$

For parameter estimation, and since the observation times of the different individuals are unknown, it is assumed that observation time is exponentially distributed with parameter 1 (the probabilities of observing the different events are invariant under rescalings of the $\lambda_i$ and the $\lambda_s$, the rate of the time to observation — Gerstung *et al.*, 2009).

In EvAM-Tools we include two versions of CBN that differ in the algorithm (and, thus, in speed and in how many observations can be analyzed) and in the error model: H-CBN, described in Gerstung *et al.* (2009, 2011), and MC-CBN, described in Montazeri *et al.* (2016).

Unless qualified otherwise (i.e., saying MC-CBN), when we say "CBN" we refer to H-CBN. By default, MC-CBN is not selected as a method to be used in the web app because it is often much slower than any of the remaining methods; but H-CBN, although faster, can only handle, at most, 14 events whereas MC-CBN can handle hundreds of events.

H-CBN uses simulated annealing with a nested expectation-maximization (EM) algorithm for estimation: structure —DAG— learning is conducted with simulated annealing and parameters ($\lambda$s and $\epsilon$ —the error term; see next) are estimated using the EM algorithm (section 2.3, pp. 2810–2811 of Gerstung *et al.*, 2009). As is the case for most other methods, the key focus of the algorithm is inferring the DAG of restrictions (the poset); the selected DAG (poset) is the maximum likelihood one "(...) without additional model selection criterion such as the Akaike or Bayesian information criterion (AIC and BIC, respectively)" (Gerstung *et al.*, 2009, p. 2811). Briefly, the algorithm first finds the maximum likelihood estimates for $\lambda$s and $\epsilon$ of a given poset; a new poset is then generated from the previous one (after addition/removal of relations from the poset), the maximum likelihood estimates of $\lambda$s and $\epsilon$ computed for this new poset, and the new poset is accepted if its likelihood is larger or, if smaller, it is accepted with a probability that is a function the difference in likelihoods divided by the temperature (recall they use a simulated annealing algorithm). MC-CBN uses a Monte-Carlo EM algorithm (see Montazeri *et al.*, 2016, p. i731 for network —DAG— learning and p. i730 and Algorithm 1 in p. i731 for parameter estimation).

H-CBN and MC-CBN also differ in their error models. In H-CBN the $\lambda$s describe the true underlying model that produces the true, hidden genotypes, but the observed genotypes might differ from the true ones because of observation error; the observation error is a Bernoulli process, in which a mutation is falsely observed with probability $\epsilon$, which is assumed to be the same and independent across all sites (see also Sakoparnig and Beerenwinkel, 2012, p. 2319). In MC-CBN the model is a mixture between the CBN model and a noise component model, such as the independence model provided by a DAG where all mutations are direct descendants of the root (the empty poset; see details in Montazeri *et al.*, 2016, p. i731). The error models are, of course, part of the fitting algorithm.

##### 2.3.4 Hidden Extended Suppes-Bayes Causal Networks (H-ESBCN)

H-ESBCN (Hidden Extended Suppes-Bayes Causal Networks), described in Angaroni *et al.* (2021) (and used by its authors as part of Progression Models of Cancer Evolution, PMCE), is similar to CBN in that it is a timed model, where the parameters of the model, the $\lambda$s, are the rates of the exponentially distributed times to fixation of an event given that the parents of that event have been observed. In contrast to CBN, the dependency relationships are not limited to AND, and they can include OR and XOR. In contrast to OncoBN with respect to dependencies, H-ESBCN adds XOR relationships, but H-ESBCN allows the very same model to include AND, OR, and XOR relationships; the fitting algorithm includes automatic inference of logical formulas for these three different patterns, AND, OR, XOR.

To give an example, suppose the following DAG (we only show XOR and OR relationships, since we have already shown AND relationships in examples above, and there is nothing new with AND relationships); this example is discused, in another context, in *"An example with OR and XOR" (section 7.4)*:

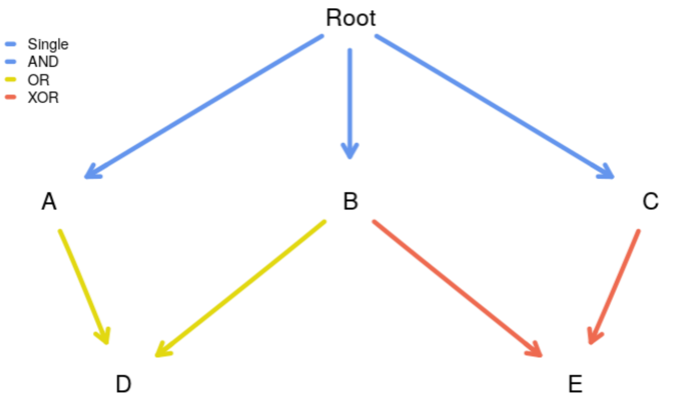

.

According to this DAG

- $A$, $B$, $C$ depend on none, and their rates are, respectively, $\lambda_A, \lambda_B, \lambda_C$.
- $D$ depends, with an OR, on both $A$ and $B$: the rate of fixation of $D$ given at least one of $A$ or $B$ have occurred is $\lambda_D$. Thus, we can observe genotypes $AD$, $BD$, $ABD$.
- $E$ depends, with an XOR, on $B$ and $C$: the rate of occurrence of $E$ given exactly one of $B$ XOR $C$ has occurred is $\lambda_E$. Thus, $E$ can only be observed in genotypes that show $B$ XOR $C$, such as genotypes $BE$, $CE$, $ABE$, $ACE$; genotypes $BCE$ or $ABCE$, in contrast, are not allowed because those genotypes have both $B$ and $C$ mutated.

The transition rate matrix between the genotypes that are possible under the model is shown below, where rows are origin, column destination (i.e., entries of $Q_{xy}$ are the transition rates from $x$ to $y$):

| | WT | A | B | C | AB | AC | AD | BC | BD | BE | CE | ABC | ABD | ABE | ACD | ACE | BCD | BDE | ABCD | ABDE | ACDE |
|---|---|---|---|---|---|---|---|---|---|---|---|---|---|---|---|---|---|---|---|---|---|
| WT | | $\lambda_A$ | $\lambda_B$ | $\lambda_C$ | | | | | | | | | | | | | | | | | |
| A | | | | | $\lambda_B$ | $\lambda_C$ | $\lambda_D$ | | | | | | | | | | | | | | |
| B | | | | | $\lambda_A$ | | | $\lambda_C$ | $\lambda_D$ | $\lambda_E$ | | | | | | | | | | | |
| C | | | | | | $\lambda_A$ | | $\lambda_B$ | | | $\lambda_E$ | | | | | | | | | | |
| AB | | | | | | | | | | | | $\lambda_C$ | $\lambda_D$ | $\lambda_E$ | | | | | | | |
| AC | | | | | | | | | | | | $\lambda_B$ | | | $\lambda_D$ | $\lambda_E$ | | | | | |
| AD | | | | | | | | | | | | | $\lambda_B$ | | $\lambda_C$ | | | | | | |
| BC | | | | | | | | | | | | $\lambda_A$ | | | | | $\lambda_D$ | | | | |
| BD | | | | | | | | | | | | | $\lambda_A$ | | | | $\lambda_C$ | $\lambda_E$ | | | |
| BE | | | | | | | | | | | | | | $\lambda_A$ | | | | $\lambda_D$ | | | |
| CE | | | | | | | | | | | | | | | | $\lambda_A$ | | | | | |
| ABC | | | | | | | | | | | | | | | | | | | $\lambda_D$ | | |
| ABD | | | | | | | | | | | | | | | | | | | $\lambda_C$ | | |
| ABE | | | | | | | | | | | | | | | | | | | | $\lambda_D$ | |
| ACD | | | | | | | | | | | | | | | | | | | $\lambda_B$ | | $\lambda_E$ |
| ACE | | | | | | | | | | | | | | | | | | | | | $\lambda_E$ |
| BCD | | | | | | | | | | | | | | | | | | | $\lambda_A$ | | |
| BDE | | | | | | | | | | | | | | | | | | | | $\lambda_A$ | |
| ABCD | | | | | | | | | | | | | | | | | | | | | |
| ABDE | | | | | | | | | | | | | | | | | | | | | |
| ACDE | | | | | | | | | | | | | | | | | | | | | |

Note that it is possible to have two (or more) parents to have dependents with different relationships. This, for example, is one of the pre-loaded DAGs in EvAM-Tools:

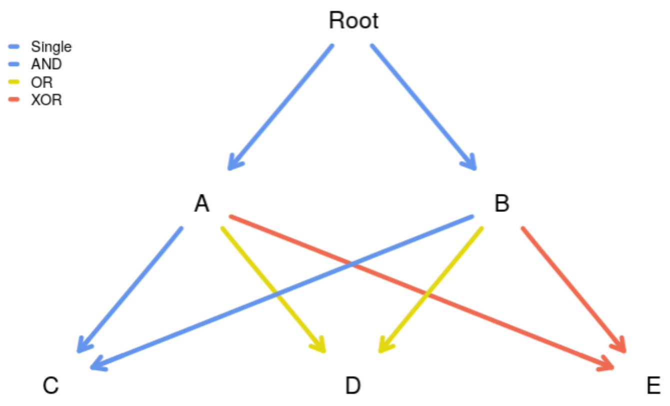

The error model is similar to the one of CBN *"Conjuntive Bayesian Networks (CBN)" (section 2.3.3)*, as described in Gerstung *et al.* (2009); Sakoparnig and Beerenwinkel (2012); see also *"CPMs: Error models" (section 2.3.6)*. The fitting algorithm is described in Angaroni *et al.* (2021, Sections 2.1, 2.2, pp. 756 and 757). As for other methods, its main focus is inferring the structure of the DAG, in this case the maximum a posteriori one in the framework of Suppe's probabilistic causation. A key feature of the algorithm is the attempt to automatically detect the correct logical formula (AND, OR, XOR) for the dependency. The structure searching algorithm uses MCMC from a randomly initialized structure which is modified according to eight different possible moves (Angaroni *et al.*, 2021, p. 756). To avoid fitting unneeded logic formulas, the structure learning algorithm includes regularization, which can be chosen by the user to be AIC or BIC. Estimation of the $\lambda$s (and error rate) for a fixed DAG structure is then done using an EM algorithm (Angaroni *et al.*, 2021, p.757).

##### 2.3.5 Mutual Hazard networks (MHN)

All of the methods described above share a model of deterministic dependencies for the accumulation of events (or mutations) (Schill *et al.*, 2020): an event (a mutation) can only occur if its dependencies are satisfied (though note that both OT and OncoBN, as well as MB-CBN allow for error deviations from this requirement — see *"CPMs: Error models", section 2.3.6*).

In contrast to the previous methods, with MHN (Schill *et al.*, 2020) dependencies are not deterministic and events can make other events more likely (promoting influence) or less likely (inhibiting influence). The rate of occurrence of events is modeled by a spontaneous rate of fixation and a multiplicative effect that each of these events can have on other events via pairwise interactions; these pairwise interactions are what allow MHN to model both promoting and inhibiting dependencies.

In more detail, the Markov process that governs the transition from a genotype $\mathbf{x}$ to a genotype with mutation $i$ added to genotype $\mathbf{x}$ is specified by (Schill *et al.*, 2020, eq. 2):

$$Q_{\mathbf{x}_{+i},\mathbf{x}} = \Theta_{ii} \prod_{x_j=1} \Theta_{ij} \tag{3}$$

where $x_j$ is 1 if gene $j$ is already mutated in genotype $\mathbf{x}$, and $Q_{y,x}$ is the transition rate from $x$ to $y$ (we are using the notation in Schill *et al.*, 2020, where transition rate matrices are transposed relative to the notation in Montazeri *et al.*, 2016 that we have used when describing CBN and H-ESBCN). $\Theta_{ii}$ is the baseline hazard or the rate of $i$ before any other events; $\Theta_{ij}$ is the multiplicative effect of event $j$ on the rate of event $i$. Therefore, equation 3

shows the transition rate as the product of the baseline hazard times the multiplicative effects of all the other mutated genes or events, $j$, on $i$.
To give a specific example, suppose the $\Theta$ matrix for a three-gene model[2] is:

$$\Theta = \begin{bmatrix} \Theta_{11} & \Theta_{12} & \Theta_{13} \\ \Theta_{21} & \Theta_{22} & \Theta_{23} \\ \Theta_{31} & \Theta_{32} & \Theta_{33} \end{bmatrix} \tag{4}$$

The following are the transition rates for some transitions:

- From WT to the genotype with the first event or gene: $\Theta_{11}$.
- From the genotype with the first event to the genotype with the first and the second events: $\Theta_{22}\Theta_{21}$.
- From the genotype with the first event and second events to the genotype with the third event: $\Theta_{33}(\Theta_{31}\Theta_{32})$.

Note that in EvAM-Tools we show the log-$\Theta$ matrix, the matrix of $\theta_{ij}$, where $\Theta_{ij} = e^{\theta_{ij}}$, because this makes it immediate to identify the inhibiting relationships as those with a negative sign, and it symmetrizes the effects around 0.
As can be seen, the relationships between events are inhibiting (event $j$ inhibits event $i$ if $\Theta_{ij} < 1$ or, equivalently, $\theta_{ij} < 0$) or promoting ($\Theta_{ij} > 1$ or, equivalently, $\theta_{ij} > 0$), but there are no deterministic restrictions (although MHN can be seen as a stochastic approximation to the deterministic dependencies of CBN: see the supplementary material of Schill *et al.*, 2020).
To fit the model, because observation time is unknown, and as is done by Gerstung *et al.* (2009), the authors assume that observation times are exponentially distributed with parameter 1. To prevent overfitting, the model fitting procedure maximizes the likelihood of the data minus an L1 penalty to try to avoid many interacting events (i.e., to promote sparsity of the fitted models): it uses a tunning parameter, $\lambda$[3] that multiplies the sum of the absolute values of the off-diagonal entries of the log $\Theta$ matrix (Schill *et al.*, 2020, eq. 6). The default value of $\lambda$ is 1/number of rows of the data set. The authors provide an efficient implementation of their method that uses a Quasi-Newton algorithm. There is no explicit error model for MHN.

##### 2.3.6 CPMs: Error models

We have mentioned error models when describing each procedure. We put together those details here, to allow for easier understanding of the similarities and differences between methods. (Methods are not ordered as above but, rather, by increasing complexity of the error model).

**MHN** There is no explicit error model (the simulation process described in p. 244 of Schill *et al.*, 2020 uses a scheme as the one in CBN, Gerstung *et al.*, 2009, explained below, but that is not part of the MHN model itself).

**CBN** In H-CBN the $\lambda$s describe the true underlying model that produces the true, hidden genotypes, but the observed genotypes might differ from the true ones because of observation error, for instance genotyping error (Gerstung *et al.*, 2009, p. 2810). The observation error is a Bernoulli process, in which a mutation is falsely observed with probability $\epsilon$, which is assumed to be the same and independent across all sites (see

[2] As a different example, see the set of transitions for a four-gene example in Schill *et al.*, 2020, Fig. 2.
[3] This $\lambda$ is different from the $\lambda$s of CBN and H-ESBCN

also Sakoparnig and Beerenwinkel, 2012, p. 2319); in other words, for all events of all subjects in the sample, if the true observation is a 0, it has a probability of being observed as a 1 of $\epsilon$, and similarly for an observation that is truly a 0.

**H-ESBCN** As for CBN (Angaroni *et al.*, 2021, p. 756).

**MC-CBN** With MC-CBN the model is a mixture between the CBN model and a noise component model, such as the independence model provided by a DAG where all mutations are direct descendants of the root (the empty poset; see details in Montazeri *et al.*, 2016, p. i730-i731). The simulations in `https://github.com/cbg-ethz/MC-CBN`, however, use a procedure where observations are generated from an underlying poset with a given set of lambdas, and symmetric error is then added (see the functions `mccbn:::random_poset` and `mccbn:::random_posets`), as for CBN above.

**OncoBN** The model includes a DBN (disjunctive) or CBN (conjunctive) model, as given by a DAG and a set of $\theta$s, and a "spontaneous activation model" (Nicol *et al.*, 2021, p. 3-4). The "spontaneous activation model", with parameter $\epsilon$, represents deviations from the model and allows child mutations to appear even if the parents in the DAG have not been mutated (i.e., even if the restrictions encoded in the DAG are not satisfied). This $\epsilon$, therefore, has a different meaning from the $\epsilon$ of CBN and H-ESBCN.

**OT** There are two sources of deviations from the OT model: a) those that result from observational (or genotyping) errors, that can lead to both false positive and false negative observational errors; b) events occurring that do not respect the OT model (Szabo and Boucher, 2002, 2008). The second source of errors would be the same as the "spontaneous activation" in OncoBN.

The `oncotree.fit` function in the `Oncotree` package returns a `eps` component with the estimated false positive, `epos` ($\epsilon_+$), and false negative, `eneg` ($\epsilon_-$), error rates. But these are the result of combining the two sources of error (Szabo and Boucher, 2008): observation errors and true deviations from the model. So observation error is reflected in both `eneg` ($\epsilon_-$), and `epos` ($\epsilon_+$), whereas true deviations from the model are only reflected in `epos` ($\epsilon_+$). In other words, the false negatives, as measured by the estimated `eneg`, are due purely to observation error. But the `epos` are not equivalent to the $\epsilon$ of OncoBN: `epos` includes both observation error (false positives) and true mutations that occur without respecting the restrictions of the OT DAG (tree).

So, when obtaining predicted frequencies under the model, for CBN, H-ESBCN, and MHN, we assume perfect compliance with the model; symmetric noise (e.g., genotyping noise) is added only when obtaining finite samples from the model. For OT and OncoBN the predicted frequencies from the model already include deviations from the "pure" model ("pure model": the model where the only possible genotypes are those that strictly respect the DAG or tree).

##### 2.3.7 CPMs: output

All methods provide directly, as output, estimates of the key constituents of their models, in particular:

**OT** Tree of restrictions, edge weights ($\pi$s), errors ($\epsilon_+, \epsilon_-$).

**OncoBN** DAG of restrictions, event $\theta$, spontaneous activation probability or error ($\epsilon$). (The type of model, conjunctive or disjunctive, is not estimated, but set by the user).

**CBN** DAG of restrictions, $\lambda$s, error rate.

**H-ESBCN** DAG of restrictions, including type of restriction (AND, OR, XOR), $\lambda$s, error rate.

**MHN** $\Theta$ matrix (or its equivalent $\theta$ or log-$\Theta$ matrices).

In addition, directly derived predictions, such as **predicted probabilities of genotypes** are provided by the original code/implementation (e.g., for OT, OncoBN, MHN) or can be obtained for CBN and H-ESBCN from the transition rate matrices (see details in *"Predicted genotype frequencies", section 3*). From the predicted probabilities of genotypes we can obtain **finite sampled genotype counts**, as explained in *"Error models and obtaining finite samples (or sampled genotype counts)" (section 5.2)*.
**Transition rate matrices** themselves are not part of the immediate output of any of the methods (except MHN[4]) but, as explained in *"Predicted genotype frequencies" (section 3)*, can be obtained from the DAG and the $\lambda$s, as we do in EvAM-Tools; we have already seen examples of the transition rate matrices for all of CBN ( *"Conjuntive Bayesian Networks (CBN)", section 2.3.3*), H-ESBCN ( *"Hidden Extended Suppes-Bayes Causal Networks (H-ESBCN)", section 2.3.4*), and MHN ( *"Mutual Hazard networks (MHN)", section 2.3.5*).
**Transition probabilities** can be computed from the transition rate matrices (for instance, using competing exponentials). And from the transition probabilities we can compute **probabilities of evolutionary paths** as the product of each transition along each possible path (see references and details in *"Probabilities of evolutionary paths and transition probabilities", section 4*).
All of this output is available from EvAM-Tools, and the web app shows most of them using both figures and tables. (Probabilities of evolutionary paths, even if asked to be computed, are not explicitly available from the web app, as they can be unwieldy to display; they are provided in the output one can download and are, of course, implicit from the transition probabilities between genotypes, and transition probabilities are displayed in the web app.)

##### 2.3.8 CPMs: summary

The following table provides a summary of the main features of each method.

[4] And we saw an example in *"Mutual Hazard networks (MHN)" (section 2.3.5)*

| Method | Timed/ untimed | Number and type of dependencies | Restrictions | Representation and output |
|---|---|---|---|---|
| OT | Untimed | Single | Deterministic | Tree with edge weights ($\pi$s) |
| OncoBN | Untimed | AND (CBN version), OR (DBN version); but a given model can only contain either AND xor OR, not both | Deterministic | DAG with event thetas ($\theta$s) |
| CBN | Timed | AND | Deterministic | DAG with event rates ($\lambda$s) |
| H-ESBCN | Timed | AND, OR, XOR | Deterministic | DAG with event rates ($\lambda$s) |
| MHN | Timed | Promoting and inhibiting (but only pairwise interactions) | Stochastic dependencies | $\Theta$ matrix (diagonal entries: baseline hazards; off-diagonal: multiplicative effects). |

### 3 Predicted genotype frequencies

#### 3.1 Predicted genotype frequencies for CBN, MCCBN, MHN, H-ESBCN

Briefly, for CBN, MCCBN, MHN, and H-ESBCN, the transition rate matrix describes the true process that generates genotypes and this matrix can be obtained from the parameters of the model ($\theta$s for MHN, $\lambda$s for the rest); we haven seen examples for all these methods in section *"Cancer Progression Models (CPMs): details" (section 2.3)*. Therefore, we can use the transition rate matrix to calculate the predicted probabilities of the different genotypes using standard results from continuous-time Markov Chains. In all cases here, we assume that the time of observation is exponentially distributed with rate 1 (as in Gerstung *et al.*, 2009 or Schill *et al.*, 2020)[5].
In more detail, obtaining the transition rate matrix from the model output is detailed in Montazeri *et al.* (2016) for CBN, and Schill *et al.* (2020) for MHN; for H-ESBCN see section 7, *"H-ESBCN: details and examples of using $\lambda$s and computing transition rate matrices and predicted genotype frequencies"*.
Once we have obtained the transition rate matrix, the fastest way to obtain the predicted genotype probabilities is using equation 4 in Schill *et al.* (2020):

$$\mathbf{p} = \int_0^\infty dt\ e^{-t}\ e^{tQ}\ \mathbf{p}_0 = [I - Q]^{-1}\mathbf{p}_0 \tag{5}$$

where $\mathbf{p}_0$ is the initial distribution (i.e., 1 for WT and 0 for the rest of the genotypes), $t$ is the time of observation (again, assumed to be exponentially distributed with parameter 1), and $Q$ is the transition rate matrix (beware: written here, as in Schill *et al.*, 2020, with $Q_{ij}$ meaning the transition rate from $j$ to $i$, in contrast to our expressions for transition rate matrices in equation 2 or the transition rate matrix in section 2.3.4). This is implemented in

[5]There is code in `evamtools`, in function `population_sample_from_trm`, to obtain samples at arbitrary collections of times —i.e., not limited to times exponentially distributed with rate 1.

the non-exported function probs_from_trm, and follows also what is done in the original Generate.pTh function from Schill *et al.* (2020). probs_from_trm is called from function evam.

Instead of using that expression, we can sample from the continuous-time Markov Chain using standard procedures (e.g., ch. 5 in Wilkinson, 2019 or Algorithm 1 in Gotovos *et al.*, 2021). Sampling is what we do in EvAM-Tools when you call sample_CPMs asking for obs_genotype_transitions or state_counts to be returned (and this sampling is implemented in the non-exported function population_sample_from_trm, and called, as needed, by function sample_CPMs).

##### 3.2 Predicted genotype frequencies for OT and OncoBN

OT and OncoBN do not return rates of a continuous-time Markov chain, but probabilities of seeing specific alterations at the time of observation. Predicted probabilities of genotypes for OT and OncoBN are obtained using the weights (OT) or $\theta$s (OncoBN), according to the expression for the probability of observing a genotype; these expressions incorporate, when predicting the genotypes, the estimated errors ($\epsilon_+, \epsilon_-$ for OT, $\epsilon$ for OncoBN; see section *"CPMs: Error models", section 2.3.6*). For example, see section 2.2 in Szabo and Boucher (2008) for OT and Figure 1 and section 2.1 in Nicol *et al.* (2021) for OncoBN or section *"OncoBN" (section 2.3.2)*. For OT we can use function distributiion.oncotree in package Oncotree and for OncoBN function Lik.genotype from package OncoBN[6].

For all methods, once we have the predicted probabilities, we can obtain a finite sample and, if we want, add observational (or genotyping) noise; see details in section *5.2, "Error models and obtaining finite samples (or sampled genotype counts)"*.

#### 4 Probabilities of evolutionary paths and transition probabilities

How to obtain probabilities of evolutionary paths for CBN and OT is detailed in Hosseini *et al.* (2019) and Diaz-Uriarte and Vasallo (2019) (see S4_Text: https://doi.org/10.1371/journal.pcbi.1007246.s006, section 3). Basically it involves computing the product of all the transition probabilities between genotypes along all paths from WT to the last possible genotype (for OT and CBN the genotype with all loci mutated). For how to obtain transition probabilities see also Diaz-Colunga and Diaz-Uriarte (2021) (specifically section 1 in S1 Appendix: https://doi.org/10.1371/journal.pcbi.1009055.s001).

The procedures to obtain transition probabilities and probabilities of evolutionary paths for H-ESBCN and MHN are similar to CBN: in all these methods we obtain probabilities of paths from the transition matrix, which is itself obtained from the transition rate matrix. The procedure with OncoBN is analogous to the one used with OT, both being untimed models (and, in both cases, obtaining probabilities of paths, as discussed in Diaz-Uriarte and Vasallo, 2019 is an abuse of the untimed model).

[6]Though for OncoBN we do not use Lik.genotype directly, as that would involve making the exact same repeated set of calls for every individual; see the non-exported function DBN_prob_genotypes in file onco-bn-process.R

### 5 Generating random CPM/EvAM models, obtaining finite samples from them, and error models

#### 5.1 Generating random CPM/EvAM models and sampling from them

We often want to generate data under the model of a CPM. Common use cases are:

- Understand what different models imply about how the cross-sectional data looks like.
- Examine how well a method can recover the true structure when the data fulfills the assumptions of a method. For instance, we would generate data under a particular model and see if the method that implements that model can recover the true structure under different sample sizes.
- Examine how a given method works, and what type of inferences it performs, when data are generated under the model of another method. For example, what is the output from MHN if the data are really coming from an H-ESBCN model?

Addressing the above needs involves:

1. Generating a random model.
2. Obtaining the predicted genotype frequencies from that model (see *"Predicted genotype frequencies", section 3*).
3. Obtaining a finite sample from the predicted frequencies of that model.
4. Using the data to answer whichever questions we had; for example, analyze the sampled data with another or the same method, plot the genotype frequencies, etc.

We explain each one in turn below, with reference to `evamtools` functions and arguments.

1. Generating a random model.

   Function `random_evam` generates random models for OT, OncoBN, CBN, MHN, OncoBN, and H-ESBCN. Details about the arguments of the function are provided in its help page. No specific provision is made for randomly generating from MCCBN, as the way to simulate is similar to CBN (generate a random poset and a random set of lambdas).

2. Obtaining the predicted genotype frequencies from that model.

   These are returned as part of the output of `random_evam` (as well as part of the output of `evam`). The predicted distribution of genotypes for a model is done assuming perfect compliance with the model; see *"Predicted genotype frequencies" (section 3)*. Remember that the model in OT and OncoBN already includes deviations from the "pure model" (see *"CPMs: Error models", section 2.3.6*).

3. Obtaining a finite sample from the predicted frequencies of that model.

   As the output from `random_evam` is the same (except for the data components) to that from `evam` we can pass the model to function `sample_CPMs`.

   When obtaining a finite sample, we can add sampling noise to the data. For example, noise due to genotyping errors; the probability of errors is controlled by argument `obs_noise` in the call to `sample_CPMs`.

   In more detail, the process involves:

(a) Obtaining a finite sample without errors from the predicted genotype frequencies.

(b) If requested (i.e., if `obs_noise > 0`), flipping a fraction `obs_noise` of the observations (i.e., turning 1s to 0s and 0s to 1s).

4. Using the data to answer whichever questions we had; for example, analyze the sampled data with another or the same method, plot the genotype frequencies, etc.

   To make this simpler, function `sample_CPMs` can return the finite sample (with or without observation noise) as a typical cross-sectional data set: a matrix where each row is a "sampled genotype", in which 0 denotes no alteration and 1 alteration in the gene of the corresponding column. This data matrix can be used directly as input for CPM methods, for instance as argument `x` (the cross-sectional data) to function `evam`.

#### 5.2 Error models and obtaining finite samples (or sampled genotype counts)

When obtaining a finite sample from a model, in all cases except OT, we have always followed the same procedure: we have first generated the predicted genotype frequencies under the model and, if requested, then added observational (e.g., genotyping) noise to the finite samples obtained from the predicted frequencies[7]. Recall that, for OncoBN, the fitted model (and, thus, the predicted frequencies under the model) already include deviations from the "pure model", as measured by $\epsilon$; see section *2.3.6, "CPMs: Error models"* and section *2.3.2, "OncoBN"*.
For OT, since `epos` ($\epsilon_+$) reflects both observation error and true deviations from the model, the above procedure is not possible. We need to introduce a difference between sampling from a model specified from scratch, such as a random model returned from function `random_evam`, and sampling from the predictions of a fitted model.
The fitted model for OT, when fitting a true data set, includes both `epos` ($\epsilon_+$) and `eneg` ($\epsilon_-$). Predicted genotype frequencies are obtained using function `Oncotree::distribution.oncotree` with, by default, argument `with.errors = TRUE`, which is what argument `with_errors_dist_ot = TRUE` to `evam` does. Therefore, from a fitted model, the predictions incorporate both false positive and false negative error rates, as estimated by `oncotree.fit`; as explained above, however, these estimated error rates are both the errors from the observational process (genotyping errors, for example) and true deviations from the model. When you later call `sample_CPMs` you can add an `obs_noise` with value larger than 0, but for OT, when sampling from the model fitted to observed data this might not make sense (since `epos` and `eneg` have been used already to produce the predicted genotype frequencies). Thus, if we use `with_errors_dist_ot = TRUE` in the `evam` call and then set `obs_noise = 0` when calling `sample_CPMs`, the observed data we generate should be the same (have the same distribution) as if we had used `Oncotree::generate.data` with `method = ''D1''`, `with.errors = TRUE` and `edge.weights = ''estimated''`.[8]
When sampling from a model specified from scratch, such as a random model returned from function `random_evam`, we generate the tree (the DAG) with density as given by argument

[7] Obtaining a finite sample of size $N$ given a vector of relative frequencies is done in R using the function `sample`.

[8] We can try to divide the `epos` component in a component like OncoBN's $\epsilon$ and another noise component. Then, we would first obtain the predicted distribution via `distribution.oncotree` with the $\epsilon$-like (and with `eneg` $= 0$), sample, and add noise. If `epos` $>$ `eneg` we can do this so that the noise added is symmetrical. This is shown in function `dot_noise_gd_3` in `inst/miscell/OT_generate_data_sample_CPMs.R`.

In that same file `inst/miscell/OT_generate_data_sample_CPMs.R` we show that a model obtained from `random_evam` with `ot_oncobn_eps` $= x$ and sampled using `sample_CPMs` with `obs_noise` $= y$ gives predictions with the same distribution as if we had used `Oncotree::generate.data` on that very same oncotree object but with `epos` $= x + y - (1/2)\ x\ y$ and `eneg` $= y$.

graph_density and the weights from uniform distributions with limits given by ot_oncobn_weight_min and ot_oncobn_weight_max. In addition, we can set a value larger than 0 for ot_oncobn_epos. This will be used as the epos, but not eneg, value of the OT model. When you sample and optionally add noise, with argument obs_noise to function sample_CPMs, noise is added symmetrically (as for the CBN model —section *2.3.6, "CPMs: Error models"*). Thus, we use a procedure where ot_oncobn_epos behaves as OncoBN's $\epsilon$ and obs_noise is purely symmetric observational error.
Why this difference? When you use a model fitted to real data, it is sensible to use Oncotree's inferential machinery to estimate the epos and eneg. If you later want to generate samples, these already include deviations from the model and noise. However, when you simulate a model, there is no data and thus no way to estimate epos and eneg. Therefore, it is sensible to split errors into two distinct pieces, which is also coherent with what we do with the rest of the methods: deviations from the model, and noise.

### 6 Random EvAM models and transitive reduction

As of now, the generation of random EvAM models uses transitively reduced graphs (we call mccbn::random_poset with argument trans_reduced = TRUE). This does not decrease the number of models that can be expressed when using CBN. However, it can limit the range of models when we can mix AND, OR, XOR in the same model. The following examples illustrate how this makes certain models impossible.

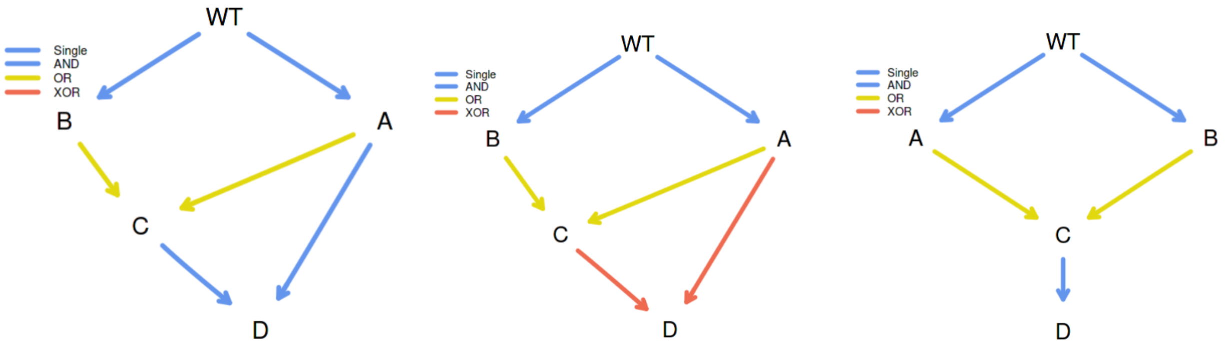

Figure 1: Non-transitively reduced DAG, OR and AND (left), non-transitively reduced DAG, OR and XOR (center), transitively reduced DAG.

- Under the left-most DAG in Fig. 1 we cannot observe genotype BCD.
- Under the center DAG in Fig. 1 we cannot observe genotype ACD.
- Under the right-most DAG, which is the transitive reduction of the above two graphs, we can observe both BCD and ACD.

Can we imagine biological scenarios where the left-most or center scenarios in Fig. 1 would apply? Yes. We don't recall seeing them in the literature, though. If this is deemed relevant, it is just a matter of changing trans_reduced = TRUE when we are simulating HESBCN models inside function random_evam.
You can of course construct the non-transitively reduced graphs "by hand" (creating the data frame with the appropriate structure) or, much simpler, using the Shiny web app.

### 7 H-ESBCN: details and examples of using $\lambda$s and computing transition rate matrices and predicted genotype frequencies

Here I provide full details about how we interpret and use the results from the method described in Angaroni *et al.* (2021). I do this here because, in contrast to CBN or MHN, there is no existing previous code or examples that do this, and we found some potentially confusing issues. I have turned this into a specific section so as not to break the flow of the former sections.

#### 7.1 Lambdas from the output: "Best Lambdas" and "lambdas_matrix"

The output returned by the H-ESBCN C code contains a "Best Lambdas" vector. The output returned by function `import.hesbcn` (that we have included in the code, in file `HESBCN__import.hesbcn.R`) has an object called "lambdas_matrix" where each of the lambdas for a gene is divided by the number of parents. This can be checked in any of the examples in the PMCE repository. Code that shows three examples, with XOR, OR, AND, is available under "inst/miscell/examples/HESBCN-lambdas-from-examples.R".
It is the output from "Best lambdas" (i.e., the undivided lambdas) that are "[the] rates of the Poisson processes of the continuous-time HMM, associated with the vertices of the model, which allow one to estimate the expected waiting time of a node, given that its predecessor has occurred." (p. 756). (What is the division? An operation that modifies an internal data structure, and just a temporary operation, done merely for implementation purposes. In line 95 of the code —as of current version, in https://github.com/BIMIB-DISCo/PMCE/blob/main/Utilities/R/utils.R— the divided lambdas are again summed, so the partition disappears: "curr_in_lambda = sum(hesbcn$lambdas_matrix[,curr_node])", and it is that value that is used in further downstream computations; email with the authors on 2021-07-09). The "Best lambdas" are returned by our modified `import.hesbcn` function.

#### 7.2 Interpreting OR and XOR (and AND)

I find Figure 1C of Angaroni *et al.* (2021) possibly confusing. First, the non-confusing part: node "D" has a rate when exactly one of B XOR C has occured, and node "G" some other rate when E or F or both E and F have occurred. (Note: the figure shows $\tau$s, not $\lambda$s. The comments here refer to the $\lambda$s).
Now the (for me, at least) possibly confusing part: it seems that the node called "B xor C" is such that B and C have the same rates of dependencies on A; in other words, it would seem to imply that $\lambda_B = \lambda_C$. Similarly, the node called "E or F" seems to indicate that both E and F have the same rate, so $\lambda_E = \lambda_F$. But this need not be so. In fact, virtually all of the examples we have looked at, and the examples in their output, do not satisfy that the rates to the genes that are part of a XOR, OR, or AND relationship are the same. For instance, in the example above of Bladder Urothelial Carcinoma (see "inst/miscell/examples/HESBCN-lambdas-from-examples.R"), KMT2D depends on KMT2C and TP53, but the rate for KMT2C, $\lambda_{KMT2C} = 0.1991$ and that for TP53, $\lambda_{TP53} = 0.8062$.
Remember that the $\lambda$ for a gene is the rate of the process until that mutation appears and is fixated, given all the dependencies of that gene are satisfied (which is, of course, the same interpretation as under CBN). Again: "[the] rates of the Poisson processes of the continuous-time HMM, associated with the vertices of the model, which allow one to

estimate the expected waiting time of a node, given that its predecessor has occurred." (p. 756).
But the rate at which the parents are satisfied can differ (as it was the case for CBN). A difference with respect to CBN is that, with CBN, if a gene D depends on three genes A, B, C, regardless of the lambdas of each of A, B, C, D can only happen once all of A, B, C are present. With H-ESBCN and with OR and XOR relationships this is no longer the case: one can see D with only A, for example.
What if some genes depend with and AND, others with a XOR and other with a OR? Just apply the rules to each type of dependency: in the HESBCN model if a gene depends on a set of genes, it has the same type of dependency on all the genes of that set.

#### 7.3 Predicted genotype frequencies

Once we have the transition rate matrix, obtaining the predicted genotype frequencies uses the same procedure as for CBN and MHN; see section *3.1, "Predicted genotype frequencies for CBN, MCCBN, MHN, H-ESBCN"*.

#### 7.4 An example with OR and XOR

In this example:

- A, B, C depend on none.
- D depends, with an OR, on both A and B
- E depends, with an XOR, on B and C
- Transition rate matrix is shown below: rows are origin, column destination. $\lambda$s are those from "Best Lambdas".

(This example is also shown in section *2.3.4, "Hidden Extended Suppes-Bayes Causal Networks (H-ESBCN)"*).

| | WT | A | B | C | AB | AC | AD | BC | BD | BE | CE | ABC | ABD | ABE | ACD | ACE | BCD | BDE | ABCD | ABDE | ACDE |
|---|---|---|---|---|---|---|---|---|---|---|---|---|---|---|---|---|---|---|---|---|---|
| WT | | $\lambda_A$ | $\lambda_B$ | $\lambda_C$ | | | | | | | | | | | | | | | | | |
| A | | | | | $\lambda_B$ | $\lambda_C$ | $\lambda_D$ | | | | | | | | | | | | | | |
| B | | | | | $\lambda_A$ | | | $\lambda_C$ | $\lambda_D$ | $\lambda_E$ | | | | | | | | | | | |
| C | | | | | | $\lambda_A$ | | $\lambda_B$ | | | $\lambda_E$ | | | | | | | | | | |
| AB | | | | | | | | | | | | $\lambda_C$ | $\lambda_D$ | $\lambda_E$ | | | | | | | |
| AC | | | | | | | | | | | | $\lambda_B$ | | | $\lambda_D$ | $\lambda_E$ | | | | | |
| AD | | | | | | | | | | | | | $\lambda_B$ | | $\lambda_C$ | | | | | | |
| BC | | | | | | | | | | | | $\lambda_A$ | | | | | $\lambda_D$ | | | | |
| BD | | | | | | | | | | | | | $\lambda_A$ | | | | $\lambda_C$ | $\lambda_E$ | | | |
| BE | | | | | | | | | | | | | | $\lambda_A$ | | | | $\lambda_D$ | | | |
| CE | | | | | | | | | | | | | | | | $\lambda_A$ | | | | | |
| ABC | | | | | | | | | | | | | | | | | | | $\lambda_D$ | | |
| ABD | | | | | | | | | | | | | | | | | | | $\lambda_C$ | | |
| ABE | | | | | | | | | | | | | | | | | | | | $\lambda_D$ | |
| ACD | | | | | | | | | | | | | | | | | | | $\lambda_B$ | | $\lambda_E$ |
| ACE | | | | | | | | | | | | | | | | | | | | | $\lambda_E$ |
| BCD | | | | | | | | | | | | | | | | | | | $\lambda_A$ | | |
| BDE | | | | | | | | | | | | | | | | | | | | $\lambda_A$ | |
| ABCD | | | | | | | | | | | | | | | | | | | | | |
| ABDE | | | | | | | | | | | | | | | | | | | | | |
| ACDE | | | | | | | | | | | | | | | | | | | | | |

##### 7.5 Three examples from actual analysis

- The code in inst/miscell/HESBCN-OR-XOR-AND-lambda-and-rates.R contains examples of how we use those lambdas (the `n`, number of steps, used is ridiculously small, and set to these tiny values just for the sake of speed).

1. OR

   - Suppose output such as this (again, see file inst/miscell/HESBCN-OR-XOR-AND-lambda-and-rates.R for how to reproduce it).

```
$adjacency_matrix
     Root A B C D
Root    0 1 1 0 0
A       0 0 0 1 1
B       0 0 0 1 1
C       0 0 0 0 0
D       0 0 0 0 0

$lambdas_matrix
     Root     A     B     C      D
Root    0 8.083 2.585 0.000 0.0000
A       0 0.000 0.000 8.914 0.2062
B       0 0.000 0.000 8.914 0.2062
C       0 0.000 0.000 0.000 0.0000
D       0 0.000 0.000 0.000 0.0000

$parent_set
       A        B        C        D
"Single" "Single"    "XOR"     "OR"

$lambdas
[1]  8.0833  2.5854 17.8277  0.4124

$edges
  From To      Edge Lambdas Relation
1 Root  A Root -> A  8.0833   Single
2 Root  B Root -> B  2.5854   Single
3    A  C    A -> C 17.8277      XOR
4    B  C    B -> C 17.8277      XOR
5    A  D    A -> D  0.4124       OR
6    B  D    B -> D  0.4124       OR
```

   - From the above output, these are the lambdas:
     $\lambda_A = 8.0833, \lambda_B = 2.5854, \lambda_C = 17.8277, \lambda_D = 0.4124$.
   - Focusing only on A, B, D, to see gene D we can follow four paths.
     - The first two involve only two mutations:

    - $WT \rightarrow A \rightarrow AD$
    - $WT \rightarrow B \rightarrow BD$
    - The first is much faster, since the rate for the transition from WT to A is 8.1 compared to 2.6 of the transition B to D (from competing exponentials, the probabilities of moving to A and B are 0.76 and 0.24, respectively).
  - In the other two paths D is the third gene to appear:
    - $WT \rightarrow A \rightarrow AB \rightarrow ABD$
    - $WT \rightarrow B \rightarrow AB \rightarrow ABD$
    - These two paths take the same time, on average: both A and B need to appear (with rates given by $\lambda_A$, $\lambda_B$) and then we need D to appear ($\lambda_D$).
  - Similarly, to get to genotype "A, B, D" we can follow these paths:
    - $WT \rightarrow A \rightarrow AB \rightarrow ABD$
    - $WT \rightarrow B \rightarrow AB \rightarrow ABD$
    - $WT \rightarrow A \rightarrow AD \rightarrow ABD$
    - $WT \rightarrow B \rightarrow BD \rightarrow ABD$
    - All of them take the same expected time, as we need A, B, and D to happen, each governed by $\lambda_A$, $\lambda_B$, $\lambda_D$, respectively.
- In terms of fitness, if we used OncoSimulR (see additional document "Using OncoSimulR to get accessible genotypes and transition matrices"), we would write, for the fitness of AB: $(1 + \lambda_A)(1 + \lambda_B)$, for AD $(1 + \lambda_A)(1 + \lambda_D)$, and for ABD $(1 + \lambda_A)(1 + \lambda_B)(1 + \lambda_D)$.
  - Note, specifically, that genotypes $AD$ and $BD$ are not fitness equivalent, unless $\lambda_A = \lambda_B$.

2. XOR

- Using the above example, and focusing only on A, B, C, these are the only ways of seeing a C:
  - $WT \rightarrow A \rightarrow AC$
  - $WT \rightarrow B \rightarrow BC$
  - As we have a XOR, no routes can go through AB.
  - The first is much faster and common than the second ($\lambda_A = 8.1; \lambda_B = 2.6$).
  - Fitness (again, this is relevant if using, for example, OncoSimulR) of $AC$ is $(1 + \lambda_A)(1 + \lambda_C)$ and of $BC$ $(1 + \lambda_B)(1 + \lambda_C)$.

3. Both OR and XOR

- There is nothing new. As an example, gaining both C and D mutations.
  - $WT \rightarrow A \rightarrow AC \rightarrow ACD$
  - $WT \rightarrow B \rightarrow BC \rightarrow BCD$
  - $WT \rightarrow A \rightarrow AD \rightarrow ACD$
  - $WT \rightarrow B \rightarrow BD \rightarrow BCD$
  - There is no path going through $AB$ since C has a XOR relationship on A and B.
  - In the first path we first need to wait for A to happen (rate $\lambda_A$) then C ($\lambda_C$) then D ($\lambda_D$).

- Same for the second, with B instead of A. The first path is much more common than the second.
- The third path transposes the order of occurrence of D and C, but takes the same average time as the third. Note that the fitness of the final genotype is the same through both routes, only the order of steps changes.
- The fourth path transposes the order of occurrence of D and C, but takes the same average time as the fourth. Note that the fitness of the final genotype is the same through both routes, only the order of steps changes.

##### 7.6 Combining AND, OR, XOR?

Nothing changes. Use the rules for AND where there is an AND, XOR where there is a XOR, OR where there is an OR. Again, in the HESBCN model if a gene depends on a set of genes, it has the same type of dependency on all the genes of that set.

### 8 FAQ

#### 8.1 Web app, figures

##### 8.1.1 In the figures, some times I get the error "Figure margins too large"

Solution: try to reduce the length of the gene names.
Longer explanation: It is impossible to accommodate, automatically, all possible use cases in terms of length of genotypes (e.g., you analyzed a data set with 10 genes, and some have names that are many characters long). We try to catch mistakes, but we might have missed some.

##### 8.1.2 In the figures, some times genotype names are truncated

This problem is related to the previous one: with very large genotype or gene names, sometimes the only way to prevent the "Figure margins too large" error is to make figure margins smaller, which can result in truncation.

##### 8.1.3 In the figures, some times the histograms are too tiny

Similar to previous problems: genotype or gene names are probably too large, so to accommodate them we need to make the rest of the plot smaller.

#### 8.2 Web app, saved output

##### 8.2.1 When data are saved, genes without mutations are excluded

When creating user data (for instance, when adding new genotypes), any gene that has no mutations is automatically excluded from the saved data, regardless of the setting for number of genes. This is a feature, not a bug. For example, suppose you set the number of genes to 3, but you only specify frequencies, or counts, for genotypes "A" and "A, B". The data set will only contain columns for genes A and B (since gene C has no mutations and it would be excluded during the analyses). See also 8.5.

#### 8.3 Web app: could we reduce the number of required clicks?

This issue was raised by one reviewer: Editing values inside the app typically must be confirmed by an additional button press or key combination. It would be more convenient if values updated automatically after pressing Enter or switching the input field.
We have taken the liberty of adding it to the FAQ because it clarifies our design decisions and provides additional information about the behavior of the GUI.
We have tried to minimize additional button presses. Below we provide a description of the current behavior, with detailed explanations of the reasons for the behavior. There are few remaining cases where additional button presses or key combinations could be avoided.

- "Set the number of genes": moving the slider has an immediate effect.
  - For "Enter genotype frequencies manually", new gene names are immediately added to "Mutations" in the "Add genotypes" box.
  - For DAG, new gene names are immediately added to the "To" and "From" lists under 1. Define DAG, "New edge".
  - For MHN, it immediately resizes the log-Θ matrix; if data have already been generated, and to prevent the data and the log-Θ matrix from being in

inconsistent states, on every change of number of genes a new data set is generated.

  - (This is not applicable to “Upload file”: setting the number of genes is not available here.)

- “Use different gene names” requires clicking on “Use these gene names” (on the popup box that is opened on clicking on “Use different gene names”). This is on purpose; first, forcing the change of gene names on switching input fields could lead to disconcerting behavior especially because the renaming is often an operation of renaming the complete set of genes, not just one of them; moreover, we try to convey that using this option carelessly will lead to confusion (several warnings are provided to minimize this careless use: on the box itself and on the tooltip). If users decide they do not want to use different names after all, they can abort the operation by clicking on “Dismiss”.

  Since this option will be used sparingly and consciously, we think forcing explicit clicks and not renaming on switching input field is the appropriate behavior.

- When uploading a file (under “Upload file”), there is no need to click on additional buttons after entering a name in “Name for data”. The user enters a string for the name, and then clicks on “Load data”; the entered string will become the name of the data when the upload is finished (and that data, with that name, will be shown on the left side, under “Examples and user’s data”).

- For DAG modification/creation, it is necessary to click on “Add edge” or “Remove edge” after selecting the “From” and “To” nodes. The alternative (adding a non-existent edge or removing an existing edge as soon as two nodes are selected), even if it removes one click, gives rise to non-obvious behavior that can be hard to understand. “Add/Remove edge” require an extra click but this extra click makes the behavior clear and explicit, and allow for correcting the From/To nodes before changing the DAG. Moreover, because we have separate buttons for “Add edge” and “Remove edge”, user errors such as trying to remove an edge that does not exist, or trying to add an edge that already exists, are much easier for users to understand: when the error message pops-up, the clicked button (“Add edge”, “Remove edge”) is still colored gray.

- For DAGs, when changing entries in the DAG table, as soon as “Ctrl + Enter” is pressed, new values of the data are generated according to the new parameters, without any need to press “Generate data from DAG model”.

  (Several parameters and/or relations can be changed without clicking “Ctrl + Enter” until all changes have been made: we move between entries of the table with Tab and “Shift + Tab”, and click “Ctrl + Enter” only at the end).

- For MHN, if a data set has been generated previously, when changing entries in the MHN table, as soon as “Ctrl + Enter” is pressed, new values of the data are generated according to the new parameters, without any need to press “Generate data from MHN model”. (As above, several entries of the table can be changed without clicking “Ctrl + Enter” until all changes have been made; we can move between cells of the table with Tab, and click “Ctrl + Enter” only at the end).

  Why generate new data for MHN on log-$\Theta$ matrix modification only if data had been previously generated, but generate it immediately for DAGs even if data had not been modified?

- We, and the users in our lab, seem to build DAG models step by step, and seem to appreciate the immediate feedback from adding/removing edges, changing the type of relationship, or modifying $\lambda$s or conditional probabilities.
- We, and the user in our lab, seem to build MHNs by first thinking about a pattern of relationships that involves more than one entry of the log-$\Theta$ matrix. Once the initial model is specified, "Generate data" is clicked. After that has happened, and to avoid the data and the log-$\Theta$ matrix from being in inconsistent states, we always force a resampling of data when an entry of the matrix is changed.
- The difference in behavior would, at most, involve one extra click with MHN.

• For DAGs, when changing the "Type of model" (OT, OncoBN, CBN/H-ESBCN), if a data set has been generated previously, new data are immediately generated, without any need to press "Generate data from MHN model". This both saves one click and prevents the model and the data from possibly being in inconsistent states.

  If no data have been generated we do not generate data. Why? For reasons similar to above. In our experience, if there is no data present, users are likely to change the model and then start modifying the DAG (adding/removing edges; changing type of relationships; changing parameters). It seems reasonable to delay the sampling until the sampling becomes necessary to prevent any possibly ambiguities or inconsistencies.

• For both DAG and MHN, switching from the input fields of "Number of genotypes to sample", "Observational noise", and "epos, $\epsilon$" (only for DAG) will not lead to generating new data unless "Generate data from DAG/MHN" is clicked. In our experience, these three parameters are often modified together; updating the data immediately after switching from the input field leads to intermediate data updates that get in the way of "modify this set of parameters, and then update according to the new set". Moreover, the operation could be slightly expensive, computationally, if "Number of genotypes to sample" is a very large number.

  Note, however, that we have changed the behavior of the app, so that changes to settings of "Number of genotypes to sample" and "Observational noise" are now preserved (within a session) and they are common to MHN and DAG. We think this will minimize needing to repeatedly change them to the desired settings (as we think it is reasonable that if a user is, say, using a sample size of 10000, this setting will be desired for both DAGs and MHN); this also prevents having to set them again to the desired value after, for example, moving to "Upload file" and back.

• When genotype data are modified (under "Change genotype's counts", and with the same behavior in "Upload file", "Enter genotype frequencies manually", "DAG", "MHN"), as soon as the user enters "Ctrl + Enter", the histogram displaying genotype frequencies is updated.

  Note, therefore, that a user can choose to modify the histogram with every change of a genotype, or modify several genotypes without clicking "Ctrl + Enter" until the end: modify the number, move with Tab to the next, modify, move with Tab to the next, etc, and only update the histogram when all modifications have been done with a single "Ctrl + Enter".

• The "Rename the data" box requires entering a name and then clicking on the button "Rename the data". This is also on purpose: we think renaming the data should be a very conscious action, and users will notice that, as soon as the "Rename the data" button is clicked, that name appears on the left side, with a blue button denoting it is the current, selected one on which modifications are being made. This should also prevent unwanted proliferation of data names that the user is not fully aware of having explicitly created. Asking for a click here, instead of renaming immediately after switching away from the input field, therefore, seems reasonable.

- Options under “Advanced options”: options are set just by changing them, without any additional clicks. For example, the “Number of MCMC iterations” (under “H-ESBCN options”) can be changed from the default 200000 to, say, 500000 just by deleting the 2 and putting a 5 without additional clicks. Likewise, changing the “Model” under “OncoBN options” requires clicking on the down arrow of the pull down menu and clicking on “Conjunctive”, without additional clicks.
- In the “Output” tab, virtually all operations do not require any additional clicks and are executed immediately. “CPMs to show” does not require extra clicks, but has a small lag of about 0.9 seconds from the first click: redrawing is a potentially expensive operation, and we do not start it until giving some reasonable time for the user to click/unclick methods to show. “Download CPM results and analyzed data” opens the standard popup box that allows to change the name and then asks for clicking on “Save”.

In summary, thus, in most cases, few or none additional button presses are needed. We think that those that remain fall into the following cases:

- The additional button or key press is unavoidable becaus of the way Shiny works.
- Not requiring this additional button or key press could lead to surprising and hard to understand behavior.
- Potentially expensive operations that we only want to execute when the user is done changing values.

#### 8.4 Web app: some genes have disappeared, and instead I see a name with an underscore (”_”)

Those events were indistinguishable, because they are completely aliased, i.e., indistinguishable, because they have identical patterns —identical columns in the data matrix—.
There are example in the additional examples files.

#### 8.5 Web app: the gene number slider changes automatically on changing data set name

This is on purpose to keep consistency and also so that we save only the needed data. See also 8.2.1.

#### 8.6 Web app: Names of genes under ”Mutations” in Create, are resorted on save/rename

This is on purpose. As explained by the tooltip, the list of genes next to ”Mutations” is kept sorted (”natural order”) showing first the genes in existing genotypes, and then other genes up to ””Number of genes”. The list will be resorted when new genes are added to genotypes. ”Mutations” is just a list of candidate gene names. You can see more (or fewer, up to the number of genes in your genotypes) by moving the slider of ”Number of genes”. On renaming of the data, we trigger a counting of the genes used, reset the slider of ”Number of genes”, and reorder the gene names next to ”Mutations”.

#### 8.7 Web app: I want to rename my genes, after I made a complex graph, and don't want to reset it. What can I do?

This is borderline behavior, and we do not encourage it, but you can do it if you set all genotypes to 0 via the "Change genotype's counts". Why is this not encouraged? Because the code is not intended for this.

#### 8.8 Web app: the names of data sets are separate for DAG, MHN, etc

This is on purpose: it should help users organize data and experiments.

#### 8.9 Web app: I do not find my data under 'Examples and user's data'

Maybe you are not looking in the right place? If you created your data using DAGs, it will not be shown under MHN, for example; see 8.8.

#### 8.10 Web app: Changing gene number with MHN and forced generation of data.

With MHN (but not DAG) , changing number of genes forces the generation of data from the model (as if you had clicked on "Generate data from MHN model") to prevent inconsistent states between the data and the model. This is on purpose.
Under MHN, adding a gene amounts to adding a row and a column, and removing a gene removing a row and a column; if we did not force a resample and you forgot to do it, the genotype data could be left in a state completely inconsistent with the model. In contrast, with DAGs, changing gene number has no effect on the model until you add/remove edges. (This is why adding/removing edges from a DAG forces data generation.)

#### 8.11 Why haven't you used method X?

We have included here what we believe are the current state-of-the-art methods that have existing public implementations that run in reasonable time. After searching the literature, we have included any method that could be deemed appropriate. We have, in fact, provided access to two very recent methods: H-ESBCN and OncoBN (and their github repos show we have contributed bug reports).
Among the remaining methods available, most of them do not seem to be developed nor used anymore. For some of these methods, their authors have developed newer methods that seem to have superseded the former methods. Some other methods have dependencies on external libraries that are not open source. And, of course, we cannot provide access to methods that have no software, or have software that will run only under proprietary systems. Some further comments are provided in S4_Text in Diaz-Uriarte and Vasallo (2019) (`https://doi.org/10.1371/journal.pcbi.1007246.s006`).
If you think we have overlooked a method that should be included, please let us know.

#### 8.12 With OncoBN sometimes I obtain DAGs that are not transitively reduced

Yes, that can happen. See details here `https://github.com/phillipnicol/OncoBN/issues/5`. There is an example of this in the additional examples.

##### 8.13 With H-ESBCN sometimes I obtain DAGs that are not transitively reduced

Yes, that can happen too. Again, there is an example of this in the additional examples.

##### 8.14 In the DAG figures, why do nodes with two or more incoming edges have only a single annotated edge with a number?

Because the number, which is the $\lambda$ (CBN, HESBCN) or $\theta$ (OncoBN) is the rate (CBN, HESBCN) or probability, conditional on the assumptions indicated by the DAG being satisfied. So the $\lambda$ or $\theta$ are per node, not per edge. For instance, suppose gene C depends on both A and B (there is an AND); and you see a number of 0.7. That is the $\lambda$ or $\theta$ for observing C mutated when both A and B are mutated.
And why then not annotate the nodes, instead of the edges? Because in our experience:

- Annotating nodes leads to more confusing figures.
- Annotating edges shows what transitions are likely/fast, an idea not conveyed by annotating nodes.

##### 8.15 Do sampled genotype frequencies and counts contain observation noise? And predicted genotype frequencies?

For all models except OT, predicted genotype frequencies do not have observation noise added. The OT model itself estimates noise, and thus predicted frequencies obtained from models fitted to observed data incorporate observation noise. See *"CPMs: Error models" (section 2.3.6)*.
When we obtain a finite sample from the predicted frequencies, you can decide to add observation noise with argument `obs_noise` to function `sample_CPMs`; what happens with OT depends on whether the predictions are from a simulated model or a model fit to observed data; see details in *"Error models and obtaining finite samples (or sampled genotype counts)" (section 5.2)*.

##### 8.16 Docker and setting up your own Shiny app

###### 8.16.1 I want to setup my own Shiny app with different default "Advanced options"

In file `EvAM-Tools/evamtools/inst/shiny-examples/evamtools/ui.R` search for "Advanced options" and modify the defaults to whatever you want.

###### 8.16.2 How can I use the Shiny app in a local intranet with load balancing using multiple Docker instances

This is well beyond the scope of this document and there are many options available. One that can work (and this is more or less what we actually do) is the following:

- Start multiple Docker instances (say, 20) by changing the range of ports, for example, 3010 to 3030.
- Use HAProxy (https://www.haproxy.org/) so that you have a single entry point for all requests to the service that are then distributed, with load balancing, to the 20 instances. You will want to use "sticky connections" (see the HAProxy documentation).

As said above, this is just a sketch of the basic procedure. There are many other options.

##### 8.16.3 If I use the Docker image for the package (rdiaz02/evamrstudio), can I run the Shiny app?

Yes, you can. Just start a browser as explained in the README (https://github.com/rdiaz02/EvAM-Tools#how-to-run-the-r-package-from-the-docker-image). Then, once in RStudio, in the R console type `runShiny()` and you will have the interactive Shiny app open. But even if you can do it, it is not clear why you'd want to do this: if you only want to run the Shiny app, the Docker image rdiaz02/evamshiny is lighter and the steps to launch it much faster.

##### 8.16.4 Why aren't you using Shiny Server?

Because we did not see it as necessary or convenient. If you want to run Shiny interactively from an R session load `evamtools` and call function `runShiny`; no need for Shiny Server.
If we want to run Shiny as a service, with Docker images it is rather straightforward to launch a bunch Docker instances and use HAProxy to access to them using load-balancing (see 8.16.2) or any other such similar solution.
Moreover, notice this in the Shiny Server documentation (https://shiny.rstudio.com/articles/shiny-server.html): "Shiny Server will host each app at its own web address and automatically start the app when a user visits the address. When the user leaves, Shiny Server will automatically stop the app. " That is not exactly what we want. We want the containers to be up and running, ready to answer requests as they come with minimal latency. Moreover, a single Shiny Server would have given access to a single instance of the app (so that if two or more users access the app, one of the users has to wait while R is busy executing what the other user is running); to give users access to multiple simultaneous instances we would have needed, for example, multiple Docker images each with its own Shiny Server.
However, we might be missing something; if you think Shiny Server would allow or ease some use cases, please let us know.

##### 8.16.5 I want to build my own Docker images

If you want to modify the Docker images, modify the Dockerfiles: Dockerfile-evam-rstudio (for the RStudio Dockerfile that launches RStudio) or Dockerfile-evam-shiny (well, for the Dockerfile that creates the container to run shiny).
Then, from the 'EvAM-Tools' directory run one or both of:

```
docker build -f Dockerfile-evam-shiny  --tag somename .
docker build -f Dockerfile-evam-rstudio  --tag somename .
```

You can now run these images, as explained in the README file..
Note: it is possible, and actually a better idea, to run docker without sudo; look a the Docker documentation: https://docs.docker.com/engine/security/rootless/).

**What if creating the image fails because of no internet connection from the container** Creating the above image requires installing R packages and that might fail because the Docker container cannot connect with the internet. The following might help: https://superuser.com/a/1582710, https://superuser.com/a/1619378. In many cases, doing `sudo systemctl restart docker` might be enough.

**Cleaning the build cache and stale old images** Sometimes (e.g., if the base containers change or you want to remove build cache) you might want to issue

```
docker builder prune
```

or the much more drastic

```
docker system prune -a
```

Please, read the documentation for both.

**Copying docker images from one machine to another** Yes, that can be done. See here, for example: https://stackoverflow.com/a/23938978

#### 9 License and copyright

This work is Copyright, ©, 2022, Ramon Diaz-Uriarte.
Like the rest of this package (EvAM-Tools), this work is licensed under the GNU Affero General Public License. You can redistribute it and/or modify it under the terms of the GNU Affero General Public License as published by the Free Software Foundation, either version 3 of the License, or (at your option) any later version.
This program is distributed in the hope that it will be useful, but WITHOUT ANY WARRANTY; without even the implied warranty of MERCHANTABILITY or FITNESS FOR A PARTICULAR PURPOSE. See the GNU Affero General Public License for more details.
You should have received a copy of the GNU Affero General Public License along with this program. If not, see https://www.gnu.org/licenses/.
The source of this document and the EvAM-Tools package is at https://github.com/rdiaz02/EvAM-Tools.
